# A quantum compatible classical continuum model explains mechanical responses of cell membranes and membrane crosslinkers

**DOI:** 10.1101/2024.12.20.628332

**Authors:** Jichul Kim

## Abstract

Both classical mechanics and quantum mechanics explain the Brownian motion. However, it is unclear whether they are compatible with each other as the physical and mathematical identity of the wavefunction in quantum mechanics has been elusive. Here, a continuum theory using grammars in classical mechanics modeling but compatible with the quantum wavefunction is introduced. The theory explains the confined Brownian motion of cell membrane inclusions interacting with extracellular matrices or cytoskeletons via elastic molecular crosslinkers. This crosslinker theory is combined into the Canham-Helfrich-Evans model for fluid membranes. Calculations through the provision of a finite element method for the combined theory reproduced measured data from adhesion molecular machineries and cell membranes. Overall, by providing physical and mathematical interpretations of the quantum wavefunction, the presented theoretical model provides improved capabilities for the realistic simulation of classical and quantum biomechanical aspects of cell membranes and membrane linker proteins.

## 1. Introduction

Understanding interactions between cell membranes and membrane crosslinkers to extracellular matrices (ECMs) or cytoskeletons is crucial for the study of cellular signaling across cell membranes. A variety of adhesion receptors and cytoplasmic adaptors that act as crosslinkers have been identified. For example, activated integrins inserted in the membrane mechanically interact with ECMs to function for migration, expression, and homeostasis (1). The talin adaptor that interacts with the membrane-inserted integrin forms physical connections between membranes and cytoskeletons to function for various physiological processes (2, 3).

As another example, CD44 membrane receptors are linked to underlying cytoskeletons to function for cellular signaling (4, 5). Despite rapid advances in experimental technologies and the accumulation of a huge amount of measured data on the membrane-crosslinker interaction, current theoretical continuum models do not fully incorporate the experimental developments.

The membrane-crosslinker interaction is complex. One end of the crosslinker is bound to a membrane-inclusion molecule or directly inserted in the membrane, while the other end is fixed on ECMs or cytoskeletons. Therefore, one end of the crosslinker is mobile by showing confined Brownian motion on the surface of deforming fluid membranes. At the same time, the spring-like crosslinker can be stretched and relaxed due to applied forces. In addition to these physical complexities, there is a viewpoint considering the Brownian motion as a quantum phenomenon while cell membranes have been explained by classical continuum mechanics. A unified theoretical and mathematical framework for classical mechanics and quantum mechanics is thus required for realistic calculations of the membrane-crosslinker complex.

In this work, simple but novel principles are addressed to incorporate all these complexities and discrepancies into a single variational framework. First, the membrane crosslinker can be considered as an elastic constraint collectively applied on a continuous area rather than a point on the membrane (Fig. 1A). It was required to assume that forces at the fixed end of the crosslinker are constant if all the other boundaries in the membrane-crosslinker complex are also fixed (even though the mobile end is under confined diffusion). Here the assumption is in line with that the membrane curvature and lateral stretching are stationary because a mechanical change of the membrane can alter the force. In this case, the constant crosslinker force might be calculated by finding the average configuration for the crosslinker under the confined diffusion, which is equivalent to finding the elastic modulus density profile on the membrane for the crosslinker (Fig. 1B). The approach is also equivalent of finding a continuous trajectory (i.e., area) for the membrane-targeting end of the crosslinker that satisfies the minimum energy principle for a long enough time span (without the membrane deformation).

**Fig. 1.**
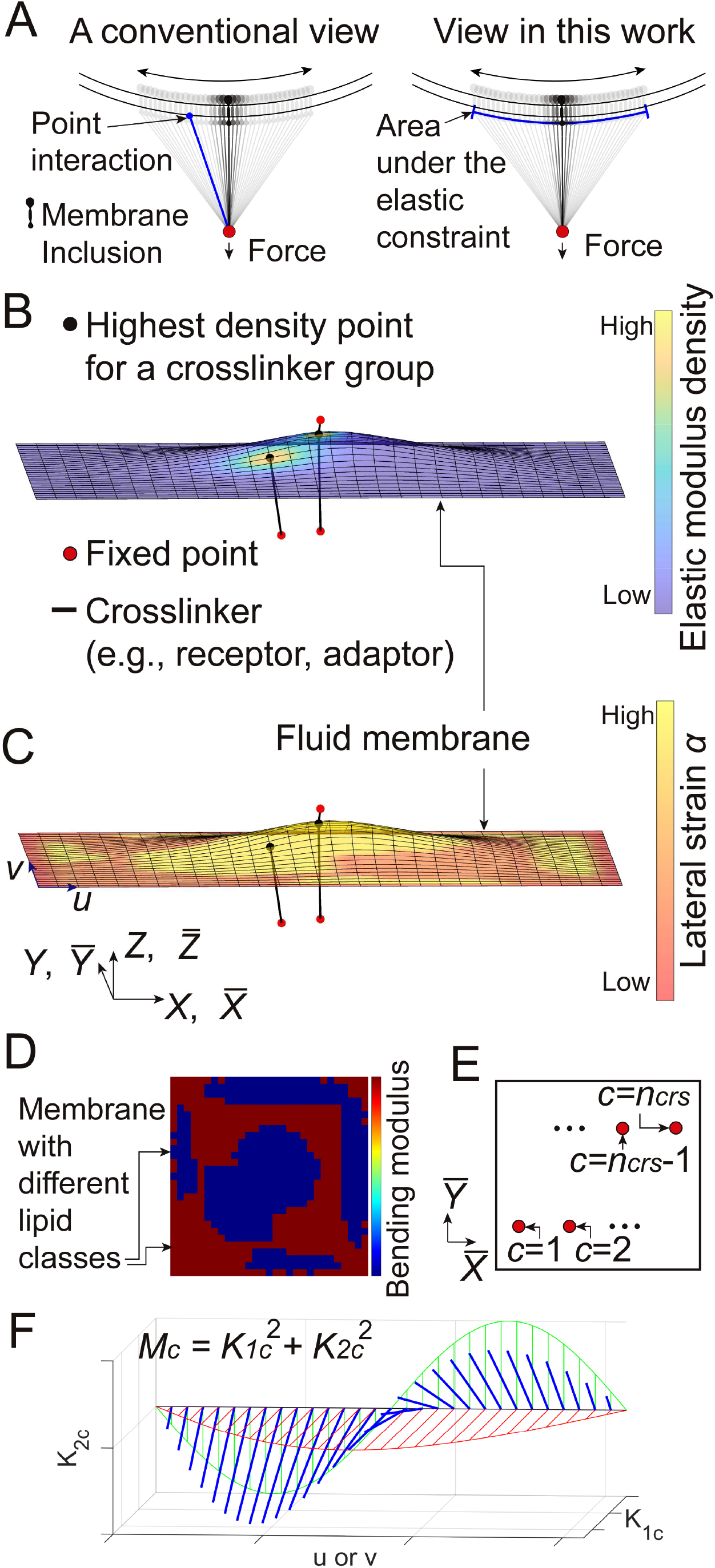
Descriptions for the membrane-crosslinker model. (**A**) Two different views for the confined Brownian motion of crosslinkers based on current classical elasticity (left) and quantum-compatible classical mechanics model in this work (right). (**A left**) The conventional viewpoint. Similar to an elastic spring on the solid membrane, a crosslinker force applied exclusively on a point in the fluid membrane can be considered. However, each point interaction during the confined diffusion may not fully consider the mobility of the crosslinker in this viewpoint. (**A right**) The alternative quantum-compatible viewpoint (in this work). With a force on the crosslinker, chances for the application of the force at multiple places on the fluid membrane can arise instantaneously with lateral slips of the crosslinker. In this case, the crosslinker might not follow the current classical elasticity universally. A more generalized model can be developed by assuming the crosslinker as an elastic constraint collectively applied on a continuous area during the confined diffusion. (**B-D**) An example configuration calculated from the finite element model of membrane-crosslinker complexes. (**B**) The elastic modulus density is shown in the surface map. (**C**) Lateral lipid number strain of the membrane is shown in the surface map. (**D**) The bending moduli of different lipid classes in the membrane are shown in the surface map. (**E**) Description of how to index the group of crosslinkers in this model. (**F**) Description of how two DOFs *K*_1*c*_ and *K*_2*c*_ define the elastic modulus density *M*_*c*_. The length of the blue line is 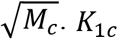 and *K*_2*c*_ satisfy ∫(*K*_1*c*_ + *K*_2*c*_) *dA* = φ_*c*_ (see Equation 1).

Second, there is a certain intrinsic material constant for the crosslinker connected to a mobile membrane inclusion. In the variational framework, this constant is linked mathematically to the ability of providing all possible configurations for the elastic modulus density profile on the membrane. This was achieved by defining the modulus density with two degrees of freedom (DOFs) (Fig. 1F). Here the two DOFs correspond to real and imaginary parts of the wavefunction in quantum mechanics. The area integral of these two variables can serve as the intrinsic constant for the crosslinker with the membrane inclusion.

The crosslinker theory is coupled into continuum models for lipid membranes (6-10). Calculations by providing a finite element method for the combined model (Fig. 1B-D) directly reproduced numerous experimental data on adhesion molecular machineries and cell membranes. Furthermore, the model provided biological insight into the following: how cell membranes can be compartmentalized with a minimal number of membrane-cytoskeleton crosslinkers; how applied forces on the crosslinker can differ depending on the mobility condition at its one end; how the presence of different lipid classes and their sorting affect the size of lipid reservoirs in mechanical responses of membranes; and how the law of current classical elasticity might be insufficient to explain the crosslinker protein interacting with the fluid membrane. Overall, the presented theoretical model not only provides invaluable biological insights into cell membranes and their interaction with crosslinkers but also suggests potential answers for the physical and mathematical identity of the quantum wavefunction.

## 2. Results

### A combined continuum theory for cell membranes and membrane crosslinkers

Functional *Π*^*tot*^ to describe mechanical responses of a cell membrane as well as elastic crosslinkers connected to mobile membrane inclusions is defined in Equation 1.

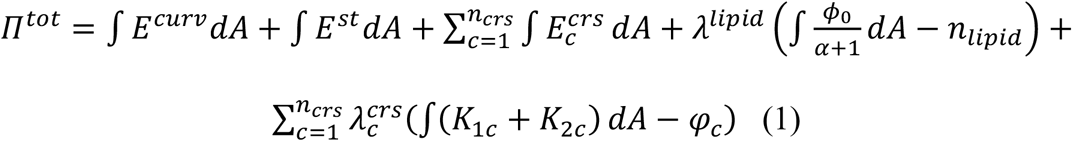

where

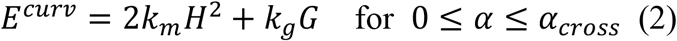

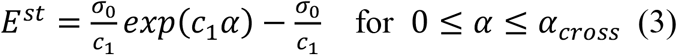

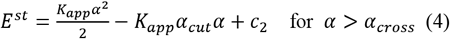

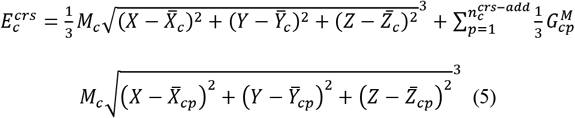

and

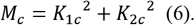

In Equation 1, the first term denotes membrane energies generated from the mean curvature *H* and the Gaussian curvature *G* that can be expressed with displacement functions *X, Y*, and *Z* in parametric coordinates *u* and *v* (6, 7, 11) (Fig. 1C, and Fig. S1). The second term provides energy due to the lateral strain *α* of membranes (8-10) (Fig. 1C). Details on membrane modeling are described in Materials and Methods and Supporting Information (SI).

Briefly, *α*_*cross*_ and *α*_*cut*_ are the crossover strain and the cut-off strain, respectively. *k*_*m*_ and *k*_*g*_ are the bending modulus and the Gaussian curvature modulus. *K*_*app*_ is the apparent area stretching modulus. *c*_1_ and *c*_2_ are constants (see Materials and Methods). *dA* is the area element. Throughout this article, all calculations were performed in the range 0 ≤ *α* ≤ *α*_*cross*_. The Gaussian curvature energy contribution in Equation 2 was omitted in this work based on the Gauss-Bonnet theorem. See Table S1 for a summary of the membrane parameters used in this work.

The expression in the third term of Equation 1 denotes elastic energies from the crosslinkers that are mobile with the membrane inclusion at one end and fixed in the three-dimensional space at the other end (Fig. 1B and C). The energy density for a group of crosslinkers is written as given in Equation 5. It was defined to have forces in a quadratic form when both ends of a single crosslinker are fixed (see Equation 7). *c* indexes the group of crosslinkers that share the membrane-targeting point (Fig. 1E). *c* also globally indexes the first crosslinker in each group. It is possible that the group is composed of a single crosslinker (Fig. 1B and C). *p* indexes additional crosslinkers within the group indexed with *c. n*_*crs*_ is the number of crosslinker groups (Fig. 1E). 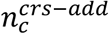 is the number of additional crosslinkers in the group indexed with *c*.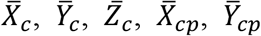, and 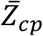 are fixed point values for the crosslinkers (red beads in Fig. 1B and C). *M*_*c*_ and 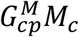 in Equation 5 are the elastic modulus density for the crosslinkers indexed with *c* and 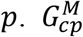 is the unitless gain. *K* and *K*_2*c*_ are two orthogonal DOFs. As illustrated in Fig. 1F, by introducing *K*_1*c*_ and *K*_2*c*_, smooth positive values for *M*_*c*_ = *K*_1*c*_^2^ + *K*_2*c*_^2^ along the membrane surface can be defined. Using only one DOF *K*_1*c*_ may limit the smoothness and variability of *M*_*c*_ in certain cases. The *M*_*c*_ value at a certain point on the membrane may provide a combined quantity for the relative residence frequency and elastic property of the mobile crosslinker. *M*_*c*_ has the unit of Newton per quartic meter i.e., N⁄m^4^. Note that for all calculated solutions in this article, *K*_1*c*_ was almost equal to *K*_2*c*_ i.e., *K*_1*c*_ ≈ *K*_2*c*_. In this case, 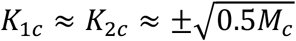 can be held.

The fourth term of Equation 1 defines a constraint to preserve the total number of lipids *n*_*lipid*_ by introducing a Lagrange multiplier *λ*^*lipid*^. Another Lagrange multiplier 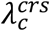 in the fifth term of Equation 1 constrains the total value of *K*_1*c*_ and *K*_2*c*_ i.e., ∫(*K*_1*c*_ + *K*_2*c*_) *dA*, to be a constant value φ_*c*_ 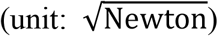 φ_*c*_ determines the distribution and magnitude of the elastic modulus density, and these reflect the confined diffusion area, residence frequency, and elastic property of the mobile crosslinker. Therefore, φ_*c*_ is an intrinsic constant that collectively characterizes the elastic property of the crosslinker as well as the interface viscosity between the fluid membrane and the membrane inclusion connected to the crosslinker. *K*_1*c*_ and *K*_2*c*_ can have both positive and negative values. Infinitely many possibilities for the positive and smooth *M*_*c*_ profile on the membrane can be made by satisfying the conservation condition ∫(*K*_1*c*_ + *K*_2*c*_) *dA* = φ_*c*_ with a given φ_*c*_ value. Among the possibilities, however, the variational principle gives us the right solution. Note that even though each crosslinker-inclusion complex with a different φ_*c*_ value can generate a set of infinitely many *M*_*c*_ profiles, the sets from different complexes are not exactly the same as each other. This might explain how φ_*c*_ can serve as an intrinsic material constant.

It is worth noting that the expressions for the crosslinker are similar to quantum mechanical descriptions―scaled versions of *K*_1_, *K*_2_, and *M* i.e., 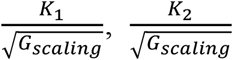, and 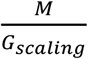 where *G*_*scaling*_ is the positive scaling factor with the unit of N⁄m^2^, may correspond to the real part of the wavefunction *Ψ* (*Re*[*Ψ*]), the imaginary part of *Ψ* (*Im*[*Ψ*]), and the probability density, i.e. the square of the modulus of *Ψ* (|*Ψ*|^2^), respectively. According to the given notions, 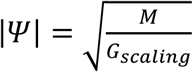 as well as 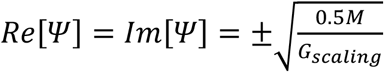 when *Re*[*Ψ*] = *Im*[*Ψ*] can be held. In this model paradigm, therefore, the wavefunction in quantum mechanics might be an energetically determined function to calculate the combined quantity for the relative spatial distribution i.e. residence frequency of a mobile object (the mobile end of a crosslinker in this work) and its attraction property with respect to its conjugate (the fixed end of the crosslinker in this work) by considering given physical constraints (the membrane deformability in this work) and the conservation condition ∫(*K*_1_ + *K*_2_) *dA* = *φ*. Overall, it is required to apply the theory to other quantum mechanical systems such as the particle in a box problem (12, 13) and atomic orbitals to demonstrate its general applicability (see Discussion). It is also worth investigating how solutions of the Schrödinger equation and the presented variational theory of the modulus density are similar or different from each other.

With given Equations 1-6, a finite element method was developed, and its full description is summarized in Materials and Methods and SI. An analysis regarding the possibility of the generation of strain gradients around membrane-inserted proteins is presented in Fig. S2 (also see SI). Finally, the method was used to investigate: (1) lipid sorting in the deformed membrane; (2) the interaction between the cell membrane and the adhesion protein integrin and talin; (3) nanomechanical responses of cell membranes; and (4) the validity of current classical elasticity for the crosslinker interacting with the fluid membrane.

### Lipid sorting and the formation of lipid nanodomains

In Fig. 2, a 1.547 × 1.547 μm^2^ planer membrane and regularly defined nine crosslinkers 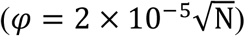 were prepared to investigate lipid sorting due to membrane curvature (14) and stretching (Fig. 2A and B). The distribution of crosslinkers was determined by simplifying recent experimental data on CD44 (4, 15). Eight lipid classes were assumed in the square membrane (see SI). For each lipid class, the total number of lipids was the same while the bending modulus *k*_*m*_ was different. With systematic mechanical inputs, the eight different lipid classes were sorted and lipid nanodomains were generated as shown in Fig. 2A (see SI for details on simulation methods). The calculated membrane configuration is reminiscent of a mosaic made by different lipid nanodomains as described previously (16). The lateral strain profile shown in Fig. 2B demonstrates that the lipid domain with a lower bending modulus is more stretched.

**Fig. 2.**
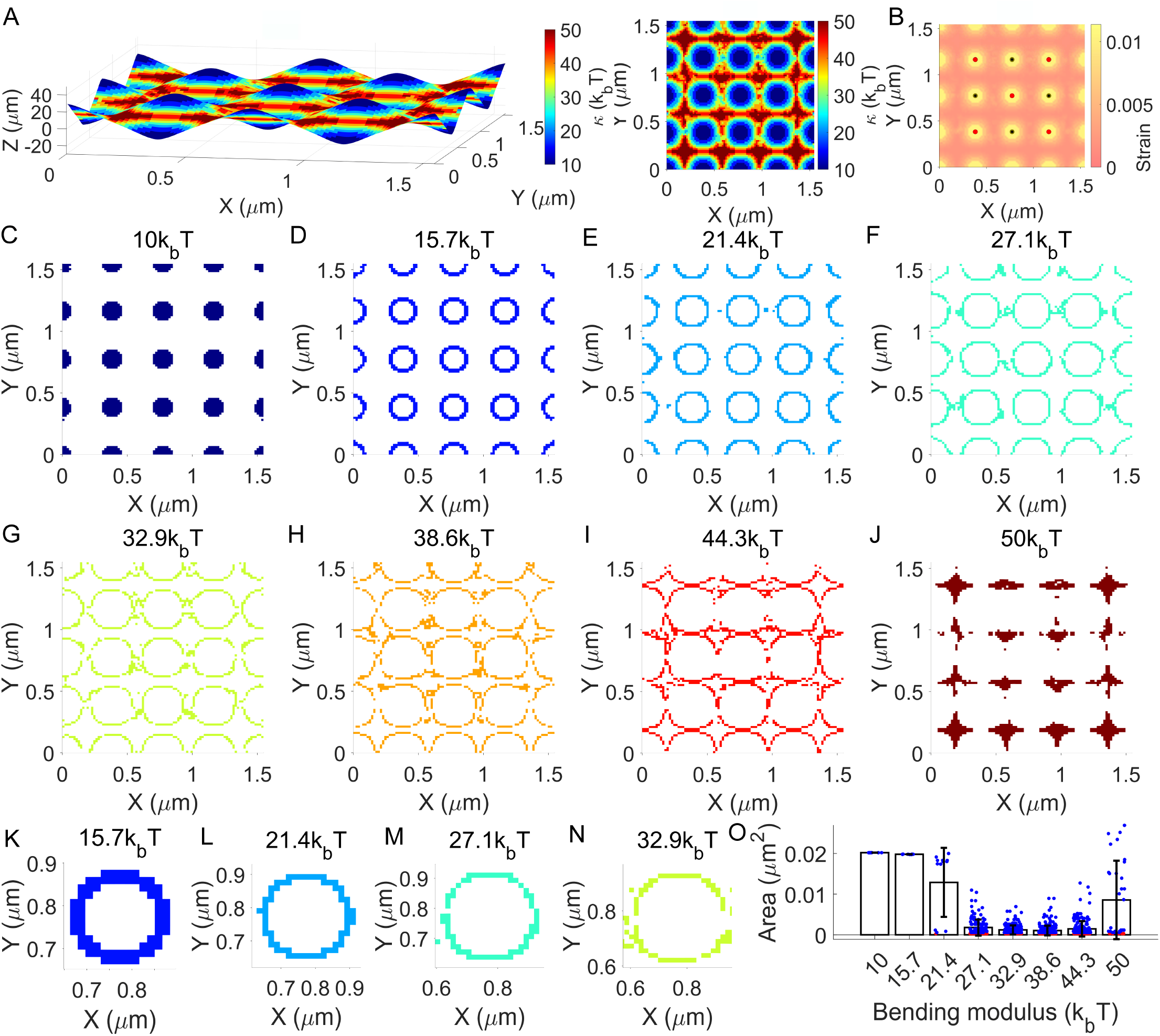
Lipid sorting and the formation of lipid nanodomains. (**A**) Three-dimensional deformation of a planar membrane with the surface map for sorting of eight different lipid classes with different bending moduli. Crosslinkers are not visualized. Left: angled view. Right: plane view. (**B**) Lipid number strain α profile for the membrane in (A). Crosslinkers are visualized. (**C-J**) The pattern of lipid nanodomains shown separately for each lipid class. (**K-N**) Expanded views for patterns in (D-G). (**O**) Plots for the mean and standard deviation (SD) of the area of the nanodomains. Mean values are 20183 nm^2^ (10*k*_*b*_*T*, SD: 11 nm^2^, n: 9), 19785 nm^2^ (15.7*k*_*b*_*T*, SD: 39 nm^2^, n: 9), 12897 nm^2^ (21.4*k*_*b*_*T*, SD: 8476 nm^2^, n: 13), 1786 nm^2^ (27.1*k*_*b*_*T*, SD: 1975 nm^2^, n: 146), 1209 nm^2^ (32.9*k*_*b*_*T*, SD: 1117 nm^2^, n: 222), 1060 nm^2^ (38.6*k*_*b*_*T*, SD: 1194 nm^2^, n: 262), 1480 nm^2^ (44.3*k*_*b*_*T*, SD: 1886 nm^2^, n: 194), and 8571 nm^2^ (50*k*_*b*_*T*, 9642 nm^2^, n: 35). Individual domain areas are shown with colored dots. Blue: multi-element domain. Red: single-element domain. See Fig. S15 for the indication of DOFs for each element. See Movie S1 for all membrane-crosslinker configurations for the analyses in this figure.

How the lipid sorting calculation can be correlated to measured compartmentalized diffusion of lipids (17-19) was also investigated. To this end, the lipid domains with different bending moduli are separately plotted in Fig. 2C-J. Fig. 2C shows the disk-shape domain, and this is similar to the area made by the confined diffusion of lipids (as an example see Fig. 4B in the reference (18)). In Fig. 2F and G (and Fig. 2M and N), donut patterns for the lipid nanodomain are demonstrated. Here multiple lipid nanodomains collectively form a single donut shape. This donut shape may explain measured hop diffusion trajectories in a closed form (see an example in Fig. 4C of the reference (18)). The donut shape with a single lipid domain in Fig. 2D and E (and Fig. 2K and L) might be consistent with the orbital trajectory of the lipid observed from supported lipid bilayers (20) and living cell membranes (21). The other patterns in Fig. 2H-J may explain other more randomized hop diffusion trajectories reported in numerous previous experiments with an assumption that a certain lipid can only diffuse to energetically favored lipid nanodomains, i.e. the domains with similar lipid parameter values. For a more direct comparison, areas of the nanodomains were calculated and plotted in Fig. 2O. These areas in the length scale of 32.6–142 *nm* were similar to measured values for the hop diffusion compartment (22).

### Mechanical responses of talin and integrins and their interaction with cell membranes

Integrins are focal adhesion receptors that interact with cytoplasmic adaptor talin (1-3). As the first step to investigate the formation of adhesions mediated by integrins, the lowest possible φ value for single monomeric talin was estimated by using previous magnetic tweezer data (23). In Fig. S3A, the experimental force vs. extension data, obtained from single full length talin monomers fixed at both ends and interacting with vinculin molecules, were compared to a calculated curve by using the following Equation 7.

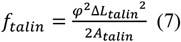

where Δ*L*_*talin*_ and *A*_*talin*_ are the talin extension and the area of the fixed region for the talin monomer, respectively (see SI for the derivation of Equation 7). In typical magnetic tweezers, forces on a magnetic bead are used to estimate forces on a target molecule fixed on the surface of the magnetic bead. Therefore, the estimated φ value here is in part based on the force calibration for the magnetic bead in the reference (23).

The *φ* value of the talin monomer was 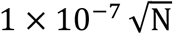 when A_talin_ = 10.2 nm^2^ (Fig. S3A). This value can be used for the talin monomer connected to the mobile integrin in a frictionless fluid membrane since the theoretical limit for the tiny area (i.e., A_talin_) is considered in Equation 7. Using 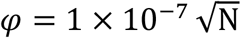 to investigate the integrin-talin complex with a 2.32 × 2.32 μm^2^ frictionless membrane resulted in much smaller talin force values as shown in Fig. S3B. For a cell membrane generating a certain viscosity, a value larger than 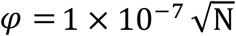 can be used for the integrin-talin complex. Overall, the results suggest that mechanical responses of a certain crosslinker can be different depending on the mobility condition applied at its one end even with the same extension.

The 2.32 × 2.32 μm^2^ membrane and two crosslinker groups with a total of three crosslinkers around the center region of the membrane were prepared as a basis complex for adhesion molecular machineries. Here the crosslinker with *c* = 1 represents a talin monomer without additional crosslinkers sharing the membrane-targeting end (see inset diagram in Fig. 3,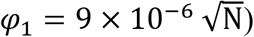 The crosslinker with *c* = 2 represents a complex of a single integrin heterodimer and its extracellular ligand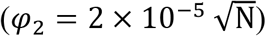. The crosslinker with (*c, p*) = (2,1) represents another talin monomer connected to the integrin-ligand complex 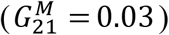. For this additional crosslinker with 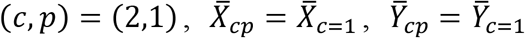, and 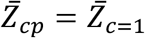 were defined initially to simulate a talin dimer whose two monomers share a fixed point on actin cytoskeletons.

**Fig. 3.**
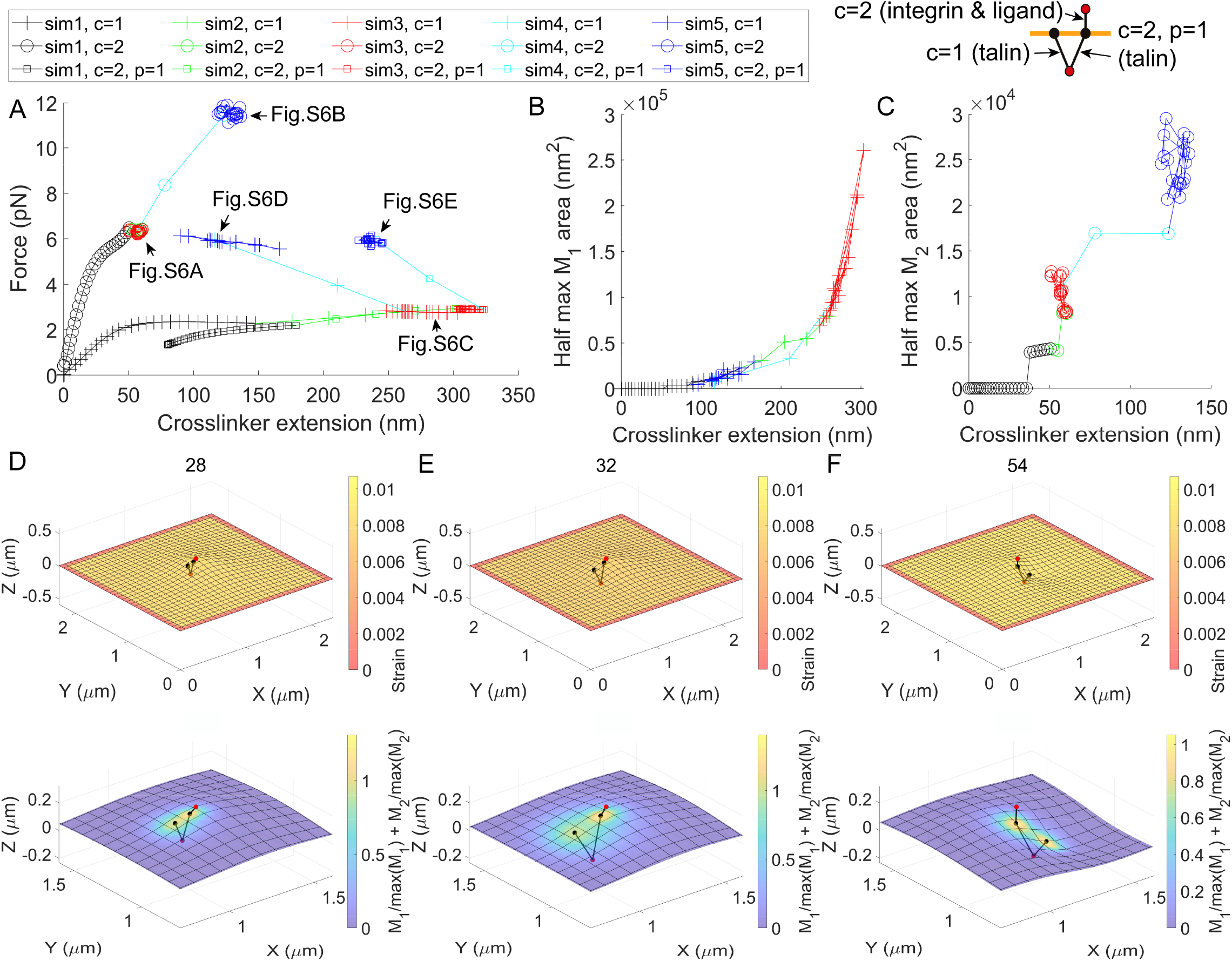
Calculations for the interaction among integrins, talin, and membranes in the formation of nascent adhesions. (**A**) Force vs. extension responses with respect to inputs in Fig. S4. The extension of the crosslinker was defined from its fixed point (red bead) to the maximum *M* point on the membrane (black bead). Expanded views for indicated regions (arrows) are provided in Fig. S6. (**B, C**) Half max *M*_*c*_ area vs. extension responses for the crosslinkers. (**D-F**) Deformed shape of the membrane-crosslinker complex. Surface maps indicate the membrane strain *α* and the sum of normalized elastic modulus densities at the top and bottom, respectively. See Fig. S5 for the membrane-crosslinker complex from a different viewpoint. See Fig. S7 for the surface map of *M*_*c*_ for *c* = 1 and *c* = 2. See Fig. S16 for the indication of DOFs. See Movie S2 for all membrane-crosslinker configurations for the analyses in this figure.

**Fig. 4.**
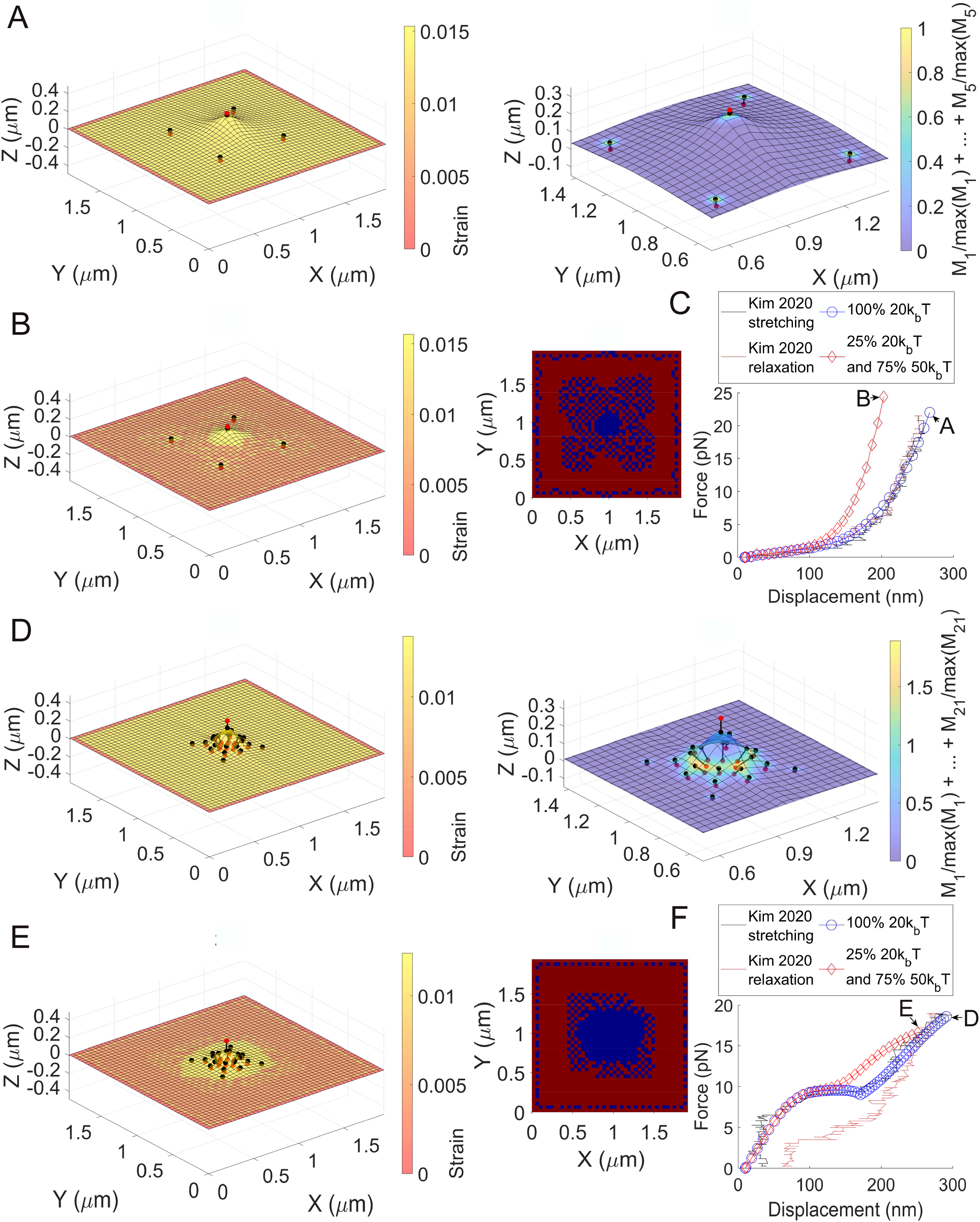
The effect of the number and distribution of crosslinkers and the composition of lipids in nanomechanical responses of membrane-crosslinker complexes. (**A**) The deformed configuration with the surface strain map for a 1.944 × 1.944 *μm*^2^ membrane (*k*_*m*_ = 20*k*_*b*_*T*) and five crosslinkers when the center crosslinker (*c* = 3) was pulled with the 249 nm displacement (left). An expanded view for the membrane-crosslinker complex with the surface map for the sum of normalized *M*_*c*_ (right). (**B**) A deformed configuration with the surface strain map for the membrane-crosslinker complex in (A) by considering 75% 50*k*_*b*_*T* lipids and 25% 20*k*_*b*_*T* lipids. The corresponding lipid sorting map is shown at right. (**C**) The force vs. displacement of the red bead at the center (*c* = 3) in (A) is shown with blue circles, and compared to experimental data (10). The force vs. displacement for the center red bead in (B) is shown with red diamonds. (**D**) The deformed configuration with the surface strain map for the 1.944 × 1.944 *μm*^2^ membrane (*k*_*m*_ = 20*k*_*b*_*T*) and twenty-one crosslinkers when the center crosslinker (*c* = 11) was pulled with the 281 nm displacement (left). An expanded view of the membrane-crosslinker complex with the surface map for the sum of normalized *M*_*c*_ (right). (**E**) A deformed configuration with the surface strain map for the membrane-crosslinker complex in (D) by considering 75% 50*k*_*b*_*T* lipids and 25% 20*k*_*b*_*T* lipids. The corresponding lipid sorting map is shown at right. (**F**) The force vs. displacement of the red bead at the center (*c* = 11) in (D) is shown with blue circles and compared to experimental data (10). The force vs. displacement for the center red bead in (E) is shown with red diamonds. See Figs. S17 and S18 for the indication of DOFs. Movies S4-S7 visualize all membrane-crosslinker complexes for the analyses in this figure.

Investigations for the integrin-membrane-talin complex were composed of five parts with different inputs (Fig. S4 and Fig. 3). The first simulation (sim1, data point index 1–28) shown with black traces was performed by pulling the crosslinker with *c* = 2. This simulation corresponds to the integrin-ligand interaction during the formation of initial adhesion. The lateral size of membrane tenting was microscopic (Fig. 3D and Fig. S5B). Forces applied on the crosslinker with *c* = 2 are plotted in Fig. 3A and Fig. S6A (black circles). Here, the final force about 6.5 pN was within one significant force population in 5.5–7 pN estimated for the integrin-ligand interaction by using fluorescence resonance energy transfer (FRET) sensors (24). By using the given φ_1_, φ_2_, and 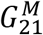 values, forces applied on the talin crosslinkers were 2.3 pN (for *c* = 1) and 2.2 pN (for (*c, p*) = (2,1)) at the end of sim1.

The second simulation (sim2, data point index 29-32) shown with green traces in Fig. S4 and Fig. 3A-C was performed by pulling the talin dimer (*c* = 1 and (*c, p*) = (2,1)) to the cytoplasmic direction. Forces applied on talin in 2.39–2.94 pN were similar to the minimum estimated talin force about 2.65 pN in a previous FRET experiment (25). Forces applied on the integrin crosslinker were still within the measured force population in 5.5–7 pN (Fig. 3A and Fig. S6A green circles) (24).

In the following simulation (sim3, red traces in Fig. S4 and Fig. 3, data point index 33–52), the fixed point for the talin dimer was randomly displaced within the area radially defined by 100 nm from the center. In this part, the average separation of two membrane targeting ends of the talin dimer was 162.38 nm when projected on the X–Y plane. This value is consistent with a previous measurement using super resolution microscopy (26). The separation was 72.9 nm when the point was randomly displaced in the area defined by 20 nm from the center. The result suggests that the motility of underlying actin networks is critical for the talin dimer separation. Finally, together with the surface map plots for *M*_*c*_ (bottom panel of Fig. 3D, E and Fig. S7A, B), the ∫ *M*_*c*_*dA* plot in Fig. S8 demonstrates that the model finds the solution, i.e. the deformed configuration of the membrane-crosslinker complex, by varying both the spatial distribution and the integrated total amount of the elastic modulus density *M*_*c*_ with given fixed φ_*c*_ and 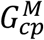 values.

The fourth simulation (sim4, data point index 53-54) shown with cyan traces was performed by increasing φ_1_ and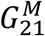. Since φ_*c*_ is the intrinsic material constant, the change of the value may correspond to structural modification of the crosslinker including its significant refolding, unfolding, and overstretching when the membrane interface is unchanged. For the case of talin, this can also occur with the change of force acting sites due to binding of vinculin along the talin rod (27-29). The change of 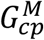 modifies the relative magnitude of the elastic modulus density of the corresponding crosslinker within its group. By increasing φ_1_ and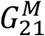, as shown with cyan traces in Fig. S4D and E to mimic the effect of talin fastening with vinculin, the initially tented membrane was able to be contracted under the X–Y plane (Fig. 3F and Fig. S5D).

In the following simulation (sim5, data point index 55–74) shown with blue traces, the shared point for the talin dimer was separated and each monomer was randomly displaced, as in Fig. S4A–C. The integrin forces were in the range from 11.2 pN to 11.9 pN. The calculations were consistent with the previous FRET experimental result that vinculin is required for the generation of integrin forces greater than 7 pN (24). With significant membrane deformation here, forces on two talin rods in 5.5–6.2 pN were also within measured values using the FRET sensors (25, 30). In sim2-5, extension values of two talin monomers were within the experimentally identified range from living cells (31).

According to previous works, integrins in living cell surfaces can undergo confined diffusion in the length scale of a hundred or hundreds of nanometers (32, 33). Furthermore, a recent experiment directly visualized that the initially freely mobile integrin was confined in a similar length scale with the talin colocalization (34). The observations support the crosslinker mechanism for the confined integrin diffusion. Remarkably, areas defined by half of the maximum *M*_*c*_ value in Fig. 3B and C were consistent with the confined integrin diffusion area in the experiments (32-34) (see SI for the calculation of the half max *M*_*c*_ area).

Stretching and sliding of crosslinkers with the deformation of membranes are visualized in Movie S2 for all data in Fig. 3. In Figs. S9, S10, and Movie S3, molecular machineries with four adhesion sites are investigated by using the same membrane and parameter values. The results were largely consistent with those from Fig. 3. Overall, the presented analyses demonstrate a remarkable ability of the model in predicting biological responses for the formation of the nascent adhesion at the single-molecule level.

### The effect of the number and distribution of crosslinkers and the composition of lipids in nanomechanical responses of membrane-crosslinker complexes

To reproduce recently identified nanomechanical responses of cell membranes (10), five crosslinkers were defined in a 1.944 × 1.944 μm^2^ membrane where the distance from one at the center to the other four crosslinkers is about 488.8 nm (Fig. 4A and Fig. S12A). The φ value for the crosslinker located at the center (*c* = 3) was 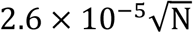. φ for the other crosslinkers (c = 1, 2,4,5) that represent the membrane-cytoskeleton linkage was 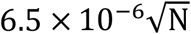. This value was selected to investigate whether φ smaller than that of the full-length talin monomer can generate the nanomechanical responses. To calculate the force vs. displacement response, the 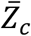 value for the center crosslinker (*c* = 3) was increased. The force vs. displacement curve in Fig. 4C (blue circles) showed a good agreement with a measured nanomechanical response of cell membranes (10).

To reproduce another type of nanomechanical response, twenty-one crosslinkers were compactly inserted in a region that spans 305.5 nm from the center of the 1.944 × 1.944 μm^2^ membrane (see Fig. 4D and Fig. S12B). The shortest distance to a crosslinker from the center was 61.1 nm (Fig. S12B). 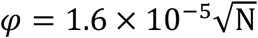 for the centered crosslinker (c = 11) while 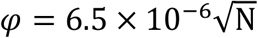 for the others were used. The increase of the number of crosslinkers within a reduced area resulted in a more localized deformation shape of the membrane. The crescent shape of the area for the elastic modulus density *M*_*c*_ around the sharp membrane curvature (c = 7, 8,14,15) demonstrates the mechanistic and hydrostatic interaction between the fluid membrane and the crosslinkers connected to mobile membrane inclusions (Fig. S13). With the given twenty-one crosslinkers, the force vs. displacement response showed zero stiffness at an intermediate displacement (Fig. 4F blue circles). A comparison between the calculated zero stiffness response and experimental data from a previous work showed a good agreement (Fig. 4F) (10). According to Fig. S14 and Movie S8, reducing the distance to the membrane-cytoskeleton linker from the center without a sufficient increase of the number of crosslinkers can result in instabilities to break the symmetry of the deformation of the membrane-crosslinker complex.

How the lipid composition affects the generation of the nanomechanical response was also investigated. The force vs. displacement responses with five and twenty-one crosslinkers were investigated again by using two different lipids (*k*_*m*_ = 50*k*_*b*_*T* and *k*_*m*_ = 20*k*_*b*_*T*) in Fig. 4C and F, respectively (red diamonds). With mechanical pulling of the center protein, two lipid classes were sorted and patterns were generated in the membrane for both cases, as shown in Movies S5 and S7. The central region with high membrane curvatures was filled with the lipid of *k*_*m*_ = 20*k*_*b*_*T* for both cases. Unlike the sorting investigation in Fig. 2, the collapse of the boundary of two lipid domains was identified as the lateral membrane stretching becomes significant with larger displacements (Fig. 4B right and E right; Movies S5 and S7). Notably, for both cases, the response in the high displacement region was shifted as the lipid of *k*_*m*_ = 50*k*_*b*_*T* was considered together with the 20*k*_*b*_*T* lipid (Fig. 4C and F). This shifting effect is consistent with the effect of reducing the size of lipid reservoirs in mechanical stretching of membranes, as analyzed in a previous work (10). According to the presented analysis, therefore, the reduced size of lipid reservoirs in membrane stretching is generated due to the presence of different lipid classes and sorting of the lipids in the deformed membrane.

### The law of current classical elasticity is not fully applicable for the crosslinker interacting with the fluid membrane

Classical mechanic laws such as Hooke’s law and nonlinear elasticity state that the force on a one-dimensional elastic spring is proportional to its extension (or a function of the extension). According to the model in this work, forces applied on a crosslinker can be different depending on the mobility condition applied at its one end. With a same extension and a φ value, the calculated force was much higher when the membrane-targeting end of the crosslinker is immobilized compared to the force with the mobile end on theoretical frictionless membranes (Fig. S3). This result suggests that the law of current classical elasticity might not be universally applicable for the crosslinker interacting with the fluid membrane―4.33 pN was expected to be applied for the 300 nm talin interacting with the frictionless membrane but only 4.3 × 10^−4^ − 4.6 × 10^−4^ pN was applied according to Fig. S3. This reduced force is the result of model calculations by additionally considering the confined Brownian motion of the crosslinker together with its extension and relaxation mechanism.

In Fig. 5, the 1.944 × 1.944 μm^2^ membrane and one crosslinker at its center were used to further investigate characteristics of the crosslinker interacting with the fluid membrane. With four different φ values, responses of the crosslinkers were significantly varied (Fig. 5A). A single φ value collectively defines the elastic property of a crosslinker as well as the viscosity for the membrane inclusion connected to the crosslinker in this model framework. Therefore, Fig. 5A suggests the possibility that force vs. extension responses of two same crosslinkers can differ if their membrane-inserted molecules are different with each other. The argument suggests that estimating the crosslinker force from the crosslinker extension by using current classical elasticity might not be always valid when the crosslinker is interacting with the fluid membrane. This interpretation is in line with the identification from Fig. S3.

**Fig. 5.**
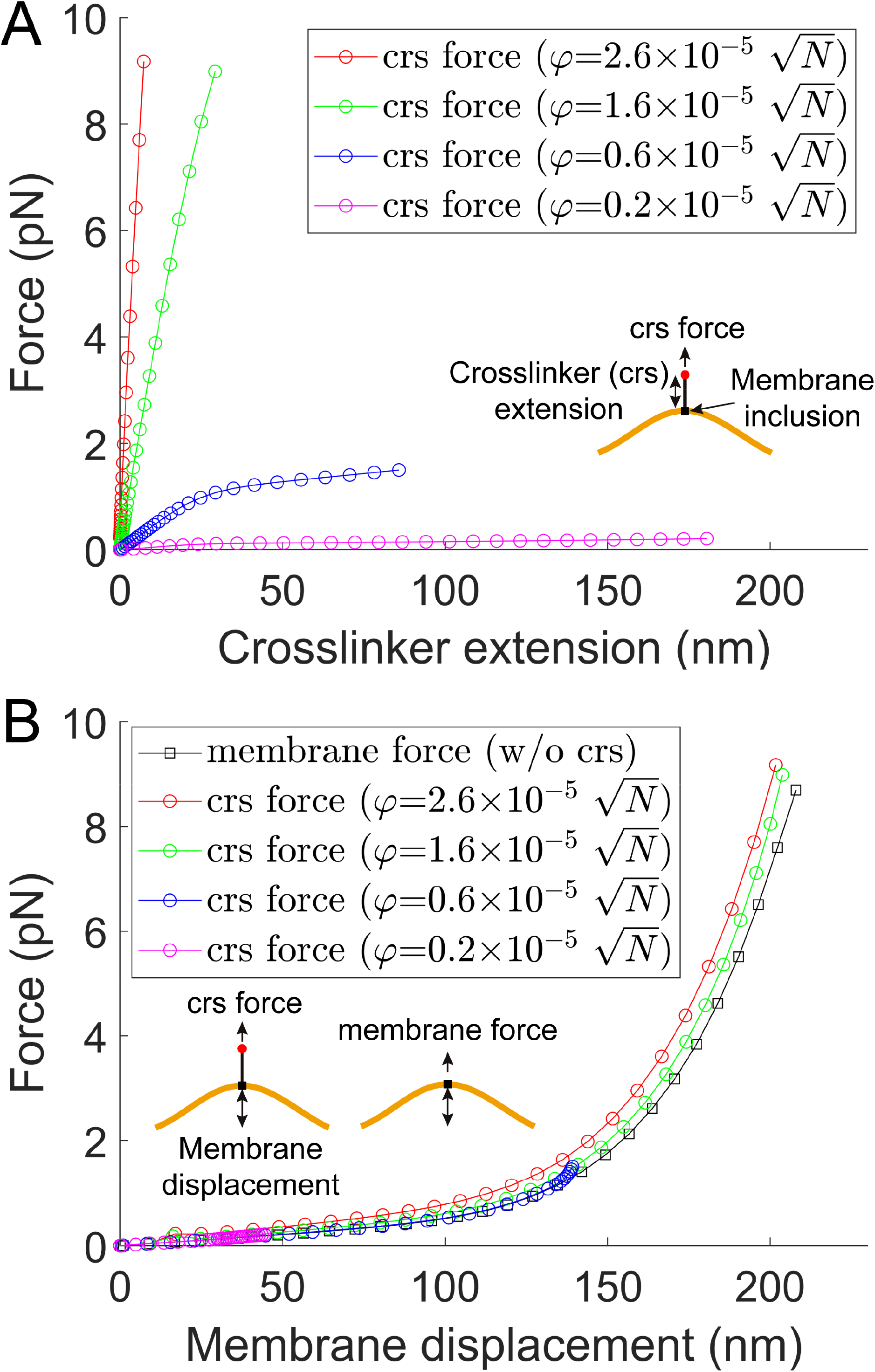
An analysis for mechanics of the elastic crosslinker interacting with the fluid membrane. (**A**) The 1.944 × 1.944 μm^2^ membrane was stimulated by using four different complexes of crosslinkers and membrane inclusions 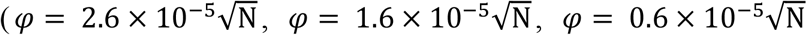, and 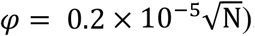. Forces applied on the crosslinkers were plotted with the extension of the crosslinkers. (**B**) The forces on the crosslinkers in (A) were plotted with the displacement of the membrane. Here the response by pulling the membrane without the center crosslinker was additionally plotted (black empty squares). See Fig. S19 for the indication of DOFs.

According to Fig. 5B, however, crosslinker force vs. membrane displacement responses were overlapped with each other for all used φ values. They were also consistent with the membrane force response directly obtained by taking the derivative of the membrane energy without the consideration of the pulling crosslinker (black empty squares in Fig. 5B). The results here suggest that the membrane deformation and Newton’s third law can be used to approximate the force on the crosslinker regardless of its elastic property and membrane-inserted interface, as similarly conducted in previous works (10, 35-37). Overall, the results in Fig. S3 and Fig. 5 suggest that current classical elasticity is not fully valid and the presented quantum-compatible classical mechanics model is a more general form to explain one-dimensional elastic materials. Finally, the results emphasize the importance of characterizing the φ value for various complexes of crosslinkers and their membrane-inserted partner molecules.

## 3. Discussion

The Brownian mobility has been considered as a quantum effect (38-40). In this work, by providing ideas potentially compatible with the quantum wavefunction but using grammars in classical mechanics modeling, the confined Brownian motion and mechanical deformation of crosslinkers interacting with deforming cell membranes were modeled. For the crosslinker interacting with the solid membrane, a one-dimensional force based on the law of current classical elasticity can be used as the crosslinker force applied exclusively on a point in the membrane (corresponding to Fig. 1A left). However, the fluid membrane cannot be exactly the same as the solid membrane because chances to apply the crosslinker force to multiple places on the membrane can arise instantaneously with lateral sliding of the crosslinker. In this work, the crosslinker was considered as an elastic constraint collectively applied on a continuous area (Fig. 1A right). The concept of the elastic modulus density for the crosslinker was introduced. Here the two orthogonal DOFs *K*_1_ and *K*_2_ for the modulus density *M* correspond to real and imaginary parts of the wavefunction.

The consistency of the calculated results with experimental data suggests that the mobile and force-bearing crosslinker might be better explained with this new approach. First, the area defined by half of the maximum *M* value for the force-bearing talin was similar to the measured area for the integrin under confined diffusion (32-34). The separated distance between two membrane-targeting ends of the talin dimer was consistent with a previous experiment (26). Finally, the model directly reproduced measured nonlinear nanomechanical responses of living cell membranes (10). These agreements collectively support the validity of the model.

The possibility of generating different force values for the same crosslinker with the same extension but with different membrane interfaces may question the validity of FRET force sensors interacting with the fluid membrane, because the technology parameterizes the extension of sensor molecules to estimate the force (24, 25, 30). In this work, however, calculated forces for talin were similar to FRET measurements from living cells (25, 30). Calculated forces for integrin-ligand interactions at adhesions were also consistent with experimental FRET data (24). These agreements suggest that the interface property between the mobile integrin and the cell membrane might be relatively less critical in determining the forces on talin in real cells. Nevertheless, more direct experimental investigations for the validity of the FRET force sensor with respect to various membrane proteins and lipid environments might be required.

One particular interest in this work was sorting of lipids and the generation of lipid nanodomains due to membrane curvatures and stretching. The size and shape of the nanodomains identified in this work were similar to those of membrane compartments identified by tracking the diffusion of lipids in living cell membranes. The picket-fence model suggests that anchored transmembrane proteins serve as physical barriers against the diffusion of lipids (17, 41). However, recently identified CD44 proteins do not fully fence a closed area in the membrane while they are responsible for the membrane compartmentalization (4, 15).

The result may ask an alternative mechanism where the compartment can be made with few picket proteins. The presented analyses suggest that a lipid can diffuse only into energetically favored nanodomains, i.e. the domains with same lipid parameter values, and that might be responsible for the generation of the observed compartmentalized diffusion. In this alternative paradigm, CD44 simply serves as the membrane-cytoskeleton crosslinker to generate membrane curvatures and stretching. The presented interpretation can also explain the hop diffusion observed in the supported lipid bilayer where the picket-fence proteins do not exist (20, 42, 43). The lipid sorting study in this work also explained how the presence of different lipid classes reduces the size of lipid reservoirs in mechanical deformation of cell membranes (10, 44-46).

As mentioned above, the demonstration of the general applicability of the modulus density theory to other systems might be important. In this regard, it would be interesting to investigate atomic orbitals by using the theory. The electron is mobile in the three-dimensional space and energetically attracted to the nucleus. Also, multiple electrons can physically constrain their motion by pushing from each other. In addition, the kinetic energy of electrons can take the electrons away from the nucleus. Therefore, the physical and mathematical characteristics of the atomic orbital may share similarities with those of the membrane-crosslinker complex investigated in this work. An electron cloud in the conventional model can be a volume with the modulus density value for the electron-nucleus interaction in this model paradigm. Likewise, the presented theory might be useful in modeling various flow phenomena with respect to liquids, information, cash, and so on. Also, conventional theories based on the spring-like, one-dimensional representation for the interaction between two fluid particles such as molecular fluid dynamic models might need to be reformulated according to this work.

In summary, the area of the confined Brownian motion for force-bearing crosslinkers was investigated by introducing a continuum theory potentially compatible with the quantum wavefunction. The variational modulus density theory for crosslinkers was developed and coupled into the continuum model for cell membranes. A novel finite element method for the combined model was developed. The finite element method not only reproduced numerous measured data but also provided invaluable predictions on the lateral organization of the cell membrane and the membrane-crosslinker interaction. The model also suggested the physical and mathematical identity of the quantum wavefunction. The theoretical model in this work is expected to provide improved capabilities for the realistic simulation of classical and quantum biomechanical aspects of cell membranes and membrane crosslinkers.

## Materials and Methods

### 1. Descriptions for the membrane model

In Equation 2 of the main text, the mean and Gaussian curvatures at a certain point on the membrane surface can be expressed as 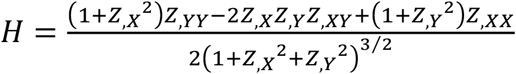 and 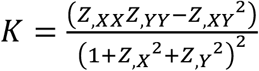, respectively (11). Here *X*, *Y*, and *Z* are displacement functions for the membrane in the three-dimensional space (Fig. 1 and Fig. S1). *Z*_,*X*_ and *Z*_,*Y*_ indicate derivatives of *Z* with respect to *X* and *Y*, respectively. Similarly, *Z*_,*XX*_, *Z*_,*XY*_, and *Z*_,*YY*_ are derivatives of *Z*_,*X*_, *Z*_,*X*_, and *Z*_,*Y*_ with respect to *X, Y*, and *Y*, respectively. The area element *dA* can be expressed as 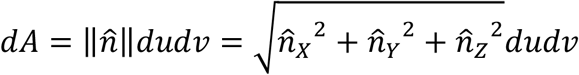, where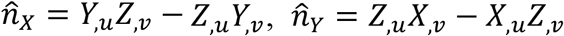, and 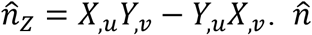 is the normal vector at a certain point on the membrane surface. *u* and *v* are parametric coordinates defined on the surface of the membrane (Fig. 1C and Fig. S1). Parametric derivatives for *X, Y*, and *Z* used for the expansion of Equation 1 are summarized in Equations S1-S30 in Supporting Information (SI).

*E*^*st*^ in Equations 3, 4 defines the strain energy density of the area *dA*. The equations were derived from two smooth expressions for the surface tension, 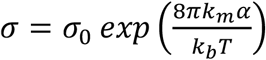 for 0 ≤ *α* ≤ *α*_*cross*_ and *σ* = *K*_*app*_(*α* − *α*_*cut*_) for *α* > *α*_*cross*_ (10, 47). Here, *α* is the lateral strain. *α*_*cut*_ and *α*_*cross*_ are the cut-off and crossover strains, respectively, that can be obtained from continuity conditions for the two surface tension expressions. *σ*_0_ is the surface tension with zero strain. *K*_*app*_ is the apparent area stretching modulus, and 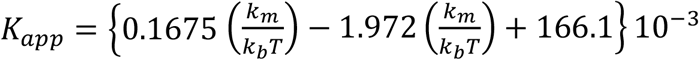 was used to define the value (47). *k*_*b*_ is the Boltzmann constant and *T* is the temperature in Kelvin. The constants *c*_1_ and *c*_2_ in Equations 3, 4 are 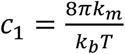 and 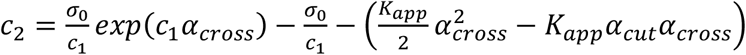, respectively (10, 47). *k*_*m*_ is the bending modulus of the membrane for 0 ≤ *α* ≤ *α*_*cross*_. Throughout this article, all calculations were performed in the range 0 ≤ *α* ≤ *α*_*cross*_ since the bending energy in Equation 2 might not be valid in *α* > *α*_*cross*_ (47).

## 2. Finite element methods for membrane-crosslinker complexes

### Variational equations for finite element modeling

The equilibrium shape of the membrane-crosslinker complex can be found by defining the variation of the given functional in Equation 1 as being equal to zero. To this end, Equation 1 can be rewritten with respect to the parametric coordinates *u* and *v* as in Equations 8-13.

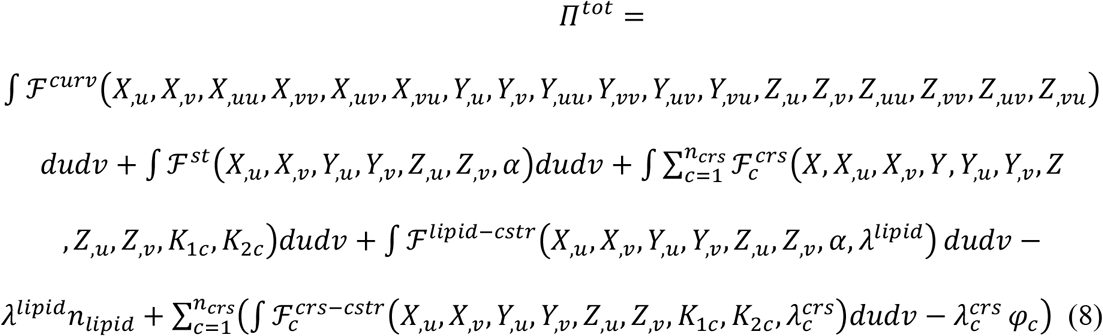

where

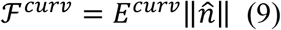

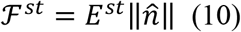

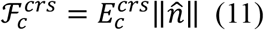

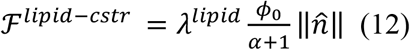

and

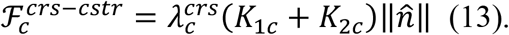

The variation of the functional i.e., δ*Π*^*tot*^, can then be written as follows.

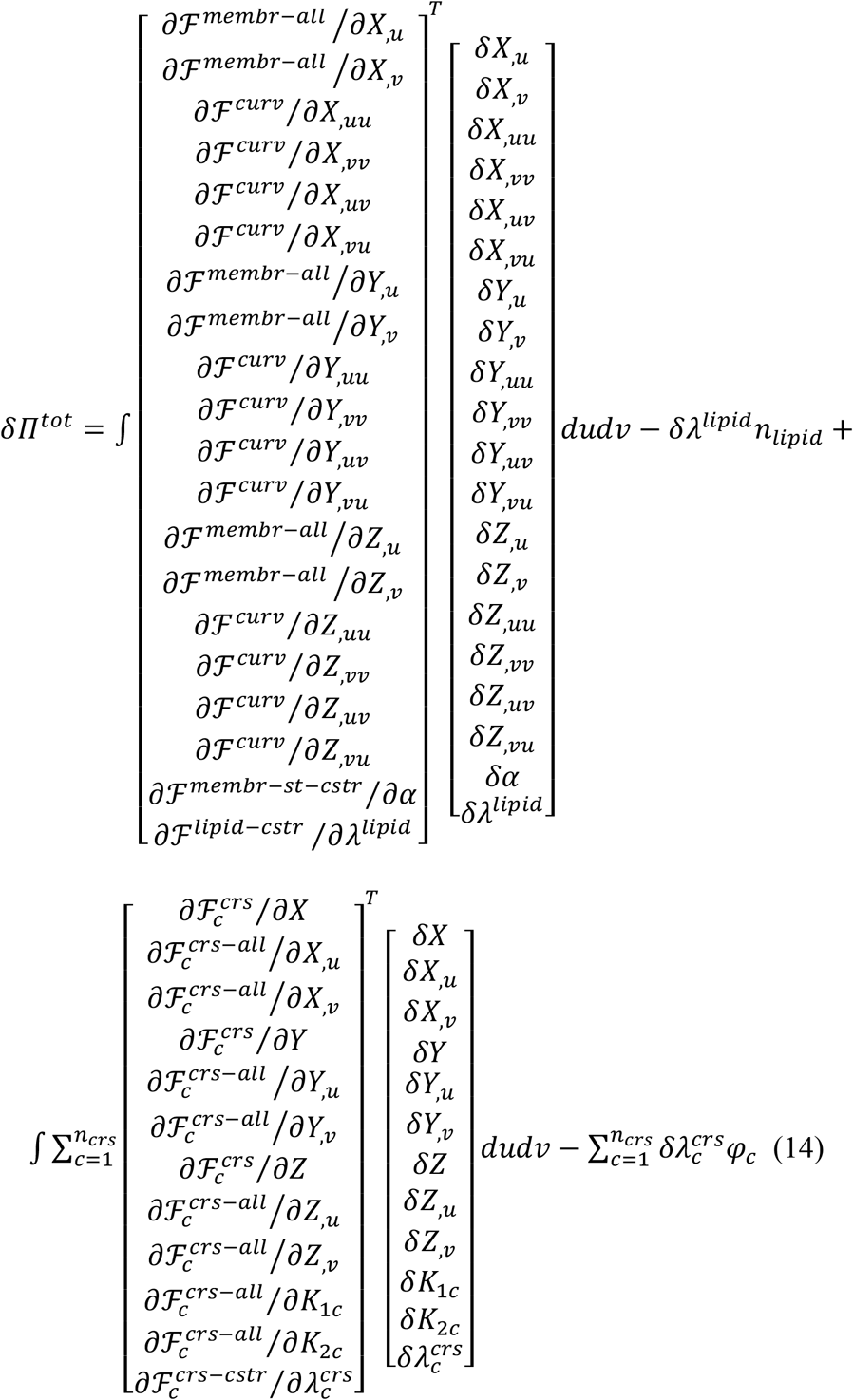

where ℱ^*membr*−*all*^ = ℱ^*curv*^ + ℱ^*st*^ + ℱ^*lipid*−*cstr*^; ℱ^*membr*−*st*−*cstr*^ = ℱ^*st*^ + ℱ^*lipid*−*cstr*^; and 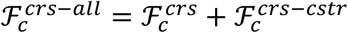. In Equation 14, δ*X*, δ*Xu*, δ*X*_*v*_, δ*X*_*uu*_, δ*X*_*vv*_, δ*X*_*uv*_, δ*X*_*vu*_ are the variation of *X, X*_,*u*_, *X*_,*v*_, *X*_,*uu*_, *X*_,*vv*_, *X*_,*uv*_, *X*_,*vu*_, respectively. The variation of the functions *Y, Z* and their derivatives; the variation of the lipid number strain *α*; the variation of *K*_1*c*_ and *K*_2*c*_; the variation of Lagrange multipliers *λ*^*lipid*^ and 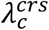 are similarly defined in Equation 14.

### B-spline based parameterization

Within the framework of parametric finite element methods, the shape of functions *X, Y, Z, α, K*_1*c*_, and *K*_2*c*_ was approximated by using the B-spline function (10). First, the function *X* was parameterized as follows.

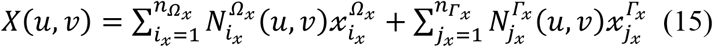

Here, Ω_*x*_ indicates the domain with the unknown 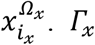 denotes the domain with the fixed boundary value 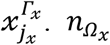 and 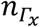 are the total number of elements for 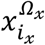 and the total number of elements for 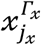, respectively. *i*_*x*_ and *j*_*x*_ are indices for 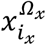 and 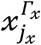, respectively. *N*(*u, v*) is the two-dimensional B-spline function. See Fig. S1C for nine basis functions used in this work, i.e. *B*_1_ to *B*_9_, to define *N*(*u, v*). Individual equations for *B*_1_ to *B*_9_ with respect to the parent domain ξ[−1,1] and *η*[−1,1] are listed in Equations S31-S39 in SI.

The surface tangential degree of freedom can cause numerical instabilities for lipid membranes. To avoid this zero-energy mode, the weight 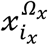 for the corresponding 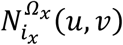 function was determined by projection of the unknown variable 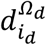 to the *X* axis. Here, 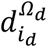was defined to the normal direction of a reference surface (10, 48). Since the membrane was stimulated in a stepwise fashion, the reference surface was derived directly or indirectly (after remeshing) from the solution of the previous step. Therefore, Equation 15 can be rewritten as follows.

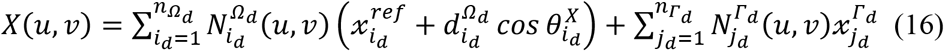

Ω_*d*_ is the domain with the unknown 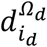 and *Γ*_*d*_ is the domain with the fixed boundary value 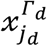 (see Fig. S1A). 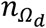 is the number of elements for 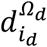, where 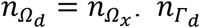 is the number of elements for 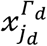, where 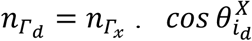 was defined by 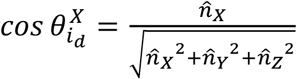 at the center of each element in the reference surface. 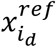 is the constant value obtained from the reference surface. Without the remeshing process, the 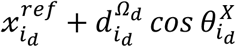 value becomes 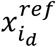 of the next step (see “Remeshing process for the finite element model” in SI for how 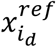is defined with the remeshing process). Similarly, the functions *Y* and *Z* can be written as follows.

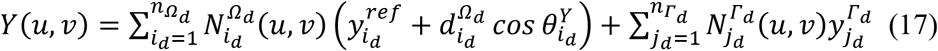

and

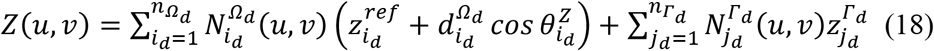

where 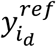 and 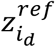 are constant values defined similarly with 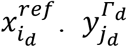 and 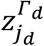 are prescribed boundary constants. 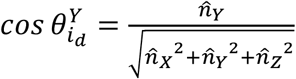 and 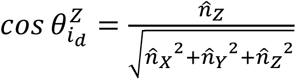 were used at the center of each element. In Equations 16-18, *i*_*d*_ is the index for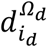, where *i*_*d*_ = *i*_*x*_. Similarly, *j*_*d*_ is the index for 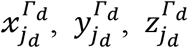 where *j*_*d*_ = *j*_*x*_. Derivatives for the functions *X, Y, Z* with respect to *u* and *v* can be defined by taking derivatives of *N*(*u, v*) from Equations 16-18. Finally, the functions *α, K*_1*c*_, and *K*_2*c*_ were written as follows.

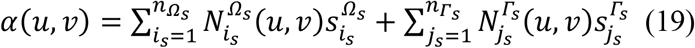

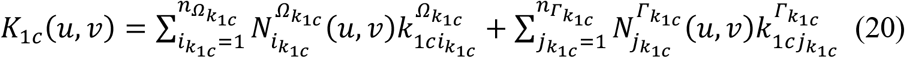

and

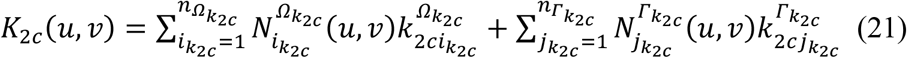

*c* is the index for the group of crosslinkers (see main text and Fig. 1E). 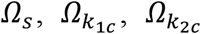 indicate the domains with the unknowns 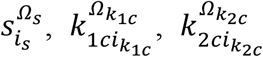, respectively. 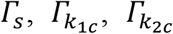 are the domains with the fixed boundary values 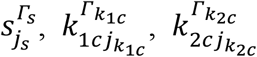, respectively. 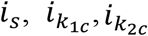 are indices for the unknowns. 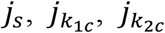 are indices for the boundary values. 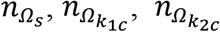 are the number of elements for 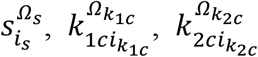, respectively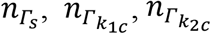 are the number of elements for 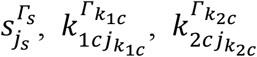, respectively.

The variation of the functions *X, Y, Z, α, K*_1*c*_, *K*_2*c*_ i.e., δ *X*, δ *Y*, δ *Z*, δ *α*, δ *K*_1*c*_, δ *K*_2*c*_, was similarly defined as written in Equations 22-27.

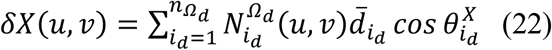

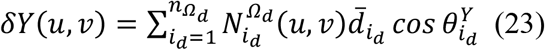

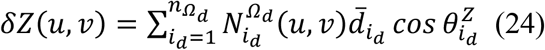

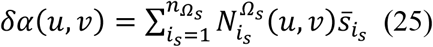

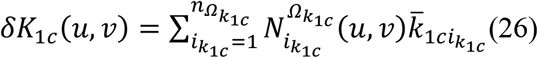

and

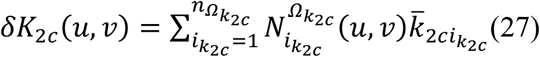

Here, 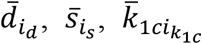, and 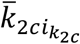 are unknown values. Derivatives of δ *X*, δ *Y*, and δ *Z* can be defined by taking derivatives of *N*(*u, v*) with respect to *u* and *v* from Equations 22-24.

### The Newton-Raphson method for nonlinear simultaneous equations

The condition to find finite element solutions is that the variation of the functional is equal to zero, i.e. δ *Π*^*tot*^ = 0. To this end, the parameterized functions *X, Y, Z* (Equations 16-18); the lipid number strain *α* (19); *K*_1*c*_ and *K*_2*c*_ (20, 21); and their variations (Equations 22-27) were substituted into Equation 14. The condition δ *Π*^*tot*^ = 0 and the arbitrariness of 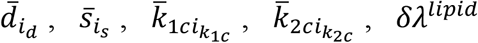, and 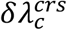 resulted in coupled *n*_*equ*_ nonlinear simultaneous equations where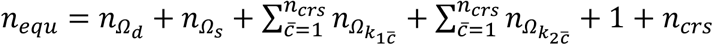.

To use the Newton-Raphson method, the residual vector 𝒢 was defined as follows.

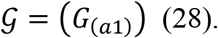

In addition, the following definitions were further made. *D*_1_ = *X, D*_2_ = *X*_,*u*_, *D*_3_ = *X*_,*v*_, *D*_4_ = *X*_,*uu*_, *D*_5_ = *X*_,*vv*_, *D*_6_ = *X*_,*uv*_, *D*_7_ = *X*_,*vu*_, *D*_8_ = *Y, D*_9_ = *Y*_,*u*_, *D*_10_ = *Y*_,*v*_, *D*_11_ = *Y*_,*uu*_, *D*_12_ = *Y*_,*vv*_, *D*_13_ = *Y*_,*uv*_, *D*_14_ = *Y*_,*vu*_, *D*_15_ = *Z*, *D*_16_ = *Z*_,*u*_, *D*_17_ = *Z*_,*v*_, *D*_18_ = *Z*_,*uu*_, *D*_19_ = *Z*_,*vv*_, *D*_20_ = *Z*_,*uv*_, *D*_21_ = *Z*_,*vu*_,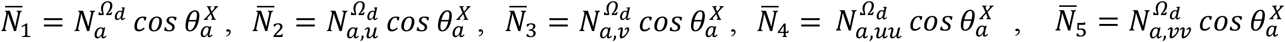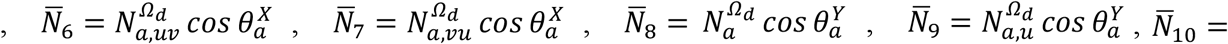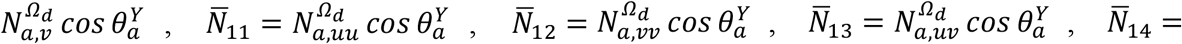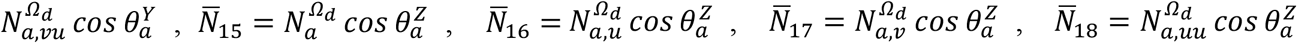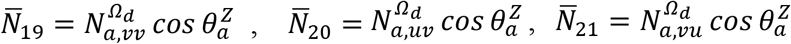 Elements of the vector 𝒢 for 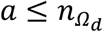 can then be defined as follows.

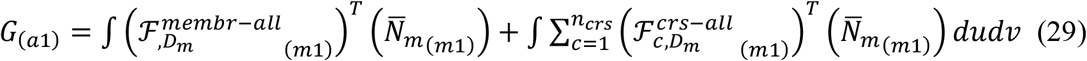

where 1 ≤ m ≤ 21. See Equation S40 in SI for the expansion of Equation 29.

Elements of 𝒢 for 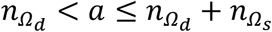 is

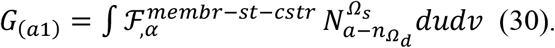

Elements of 𝒢 for 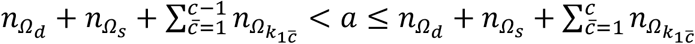 is

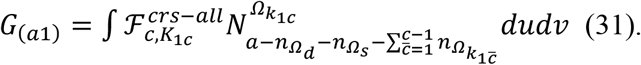

Elements of 𝒢 for 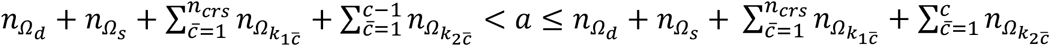 is

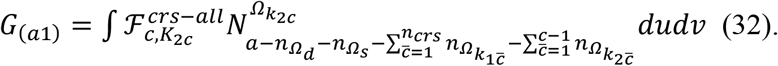

Elements of 𝒢 for 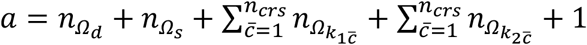 is

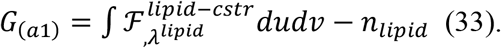

Elements of 𝒢 for 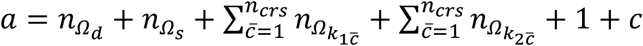 is

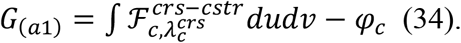

The *n*_*equ*_-by-*n*_*equ*_ Jacobian matrix 𝒥 that serves as the tangential operator in using the Newton-Raphson method is defined as follows to solve the nonlinear equations iteratively.

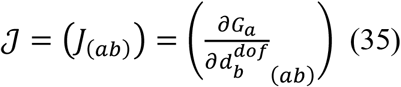

Here, *I*_(*ab*)_ indicates the element in the *a*^*th*^ row and *b*^*th*^ column of 𝒥 that can be either calculated from expressions in Equations 36-55 or defined to be zero. *G*_*a*_ is equal to *G*_(*a*1)_ in Equation 28. 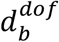 is the unknown for DOFs. 1 ≤ *m* ≤ 21 and/or 1 ≤ *n* ≤ 21 hold for Equations 36-42, 45, 48, 51, and 53.

For 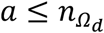 and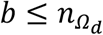,

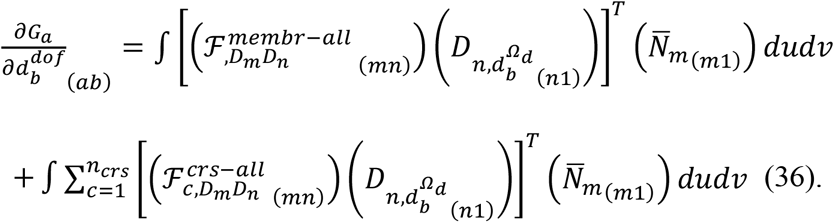

See Equation S41 for the expansion of Equation 36.

For 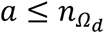 and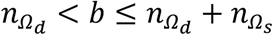,

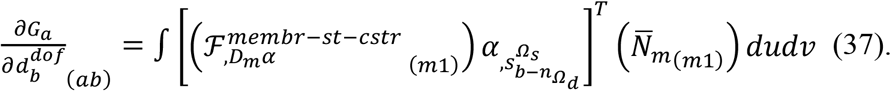

See Equation S42 for the expansion of Equation 37.

For 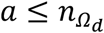 and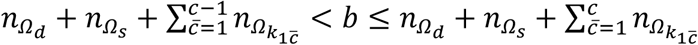,

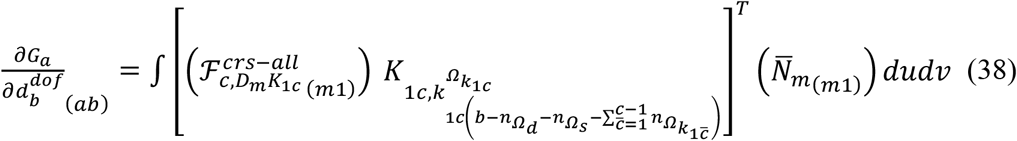

See Equation S43 for the expansion of Equation 38.

For 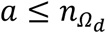 and 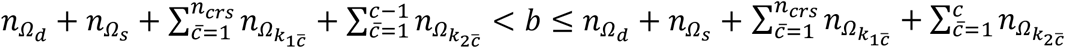,

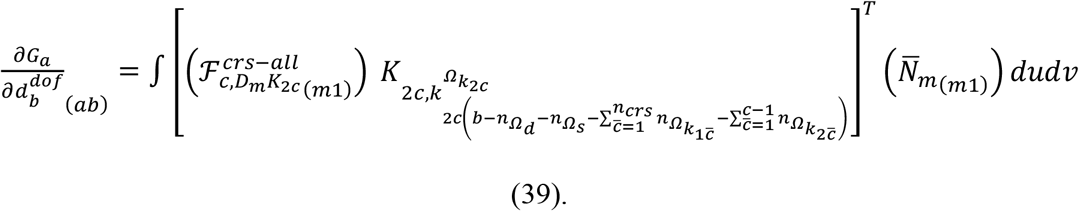

See Equation S44 for the expansion of Equation 39.

For 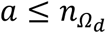 and 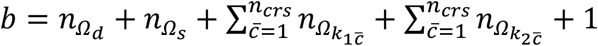,

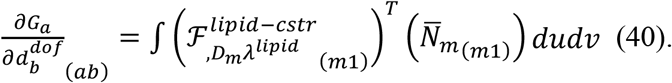

See Equation S45 for the expansion of Equation 40.

For 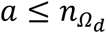 and 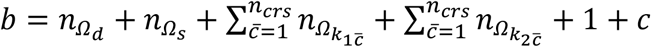

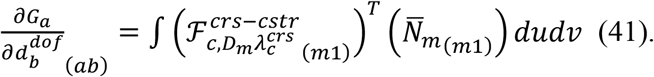

See Equation S46 for the expansion of Equation 41.

For 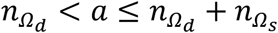 and 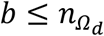

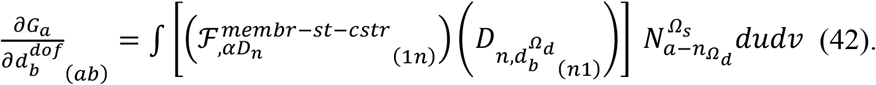

See Equation S47 for the expansion of Equation 42.

For 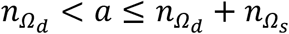 and 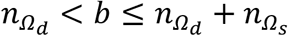,

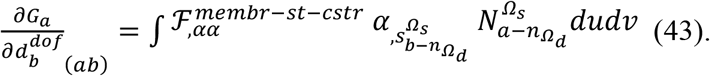

For 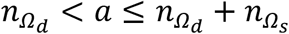 and 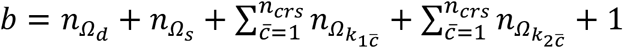,

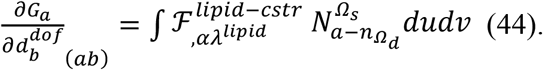

For 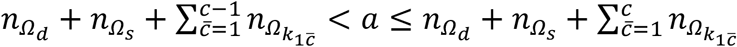 and 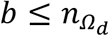,

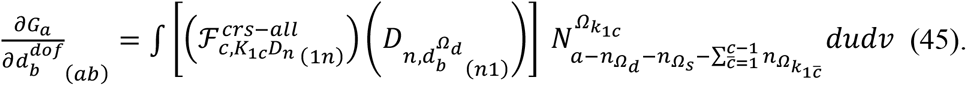

See Equation S48 for the expansion of Equation 45.

For 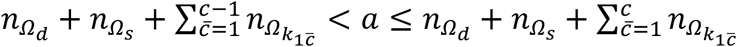 and 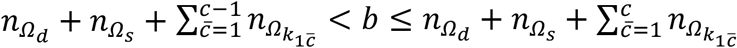,

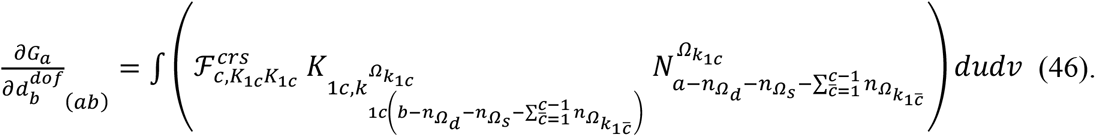

For 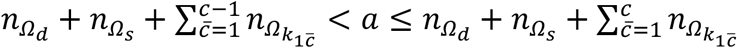 and 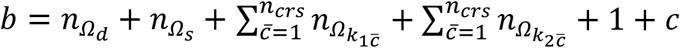,

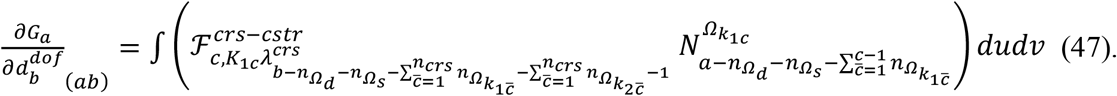

For 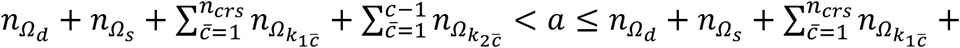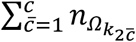, and 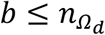,

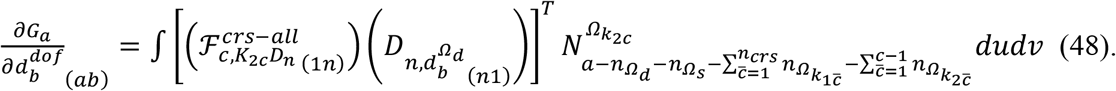

See Equation S49 for the expansion of Equation 48.

For 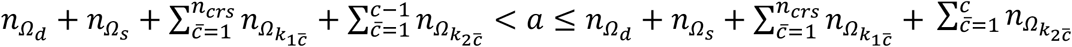 and 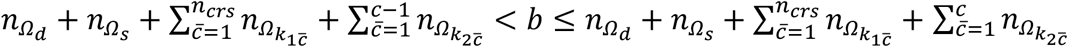,

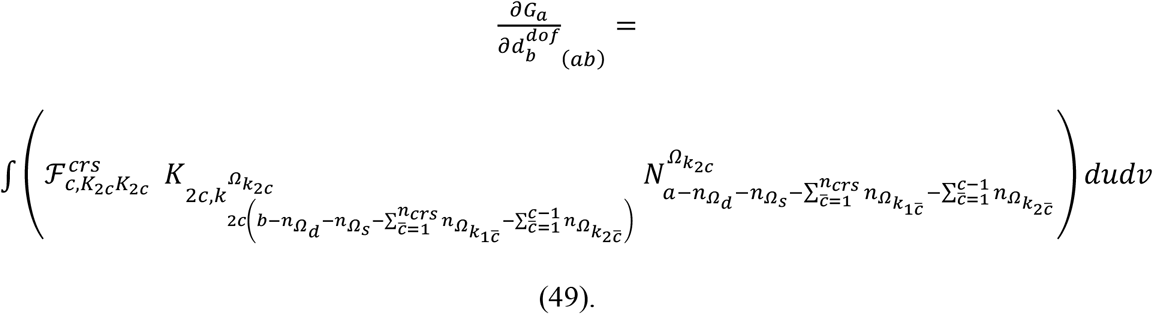

For 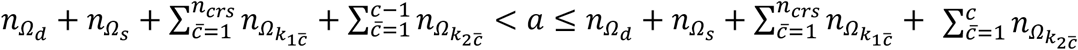 and 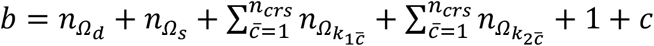,

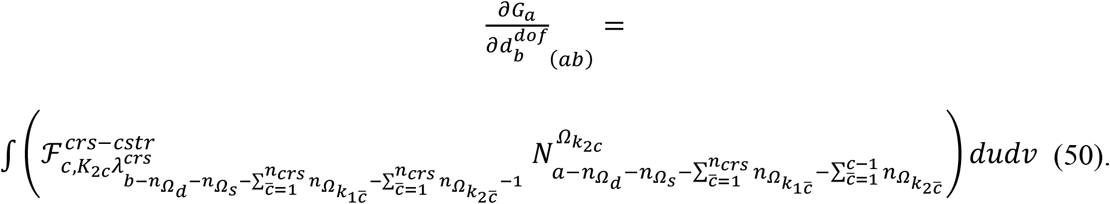

For 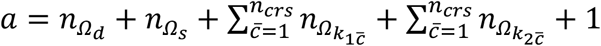 and 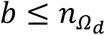,

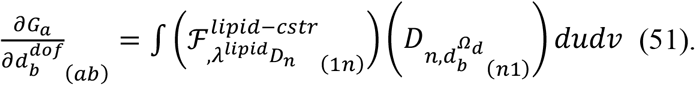

See Equation S50 for the expansion of Equation 51.

For 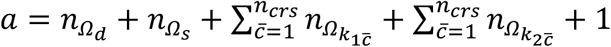 and 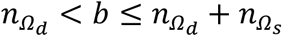

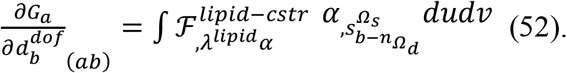

For 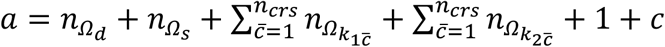 and 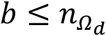,

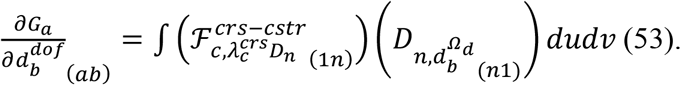

See Equation S51 for the expansion of Equation 53.

For 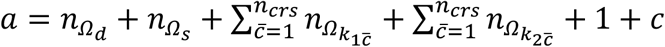 and 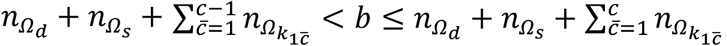,

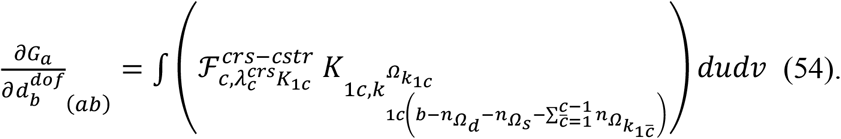

For 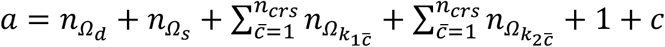 and 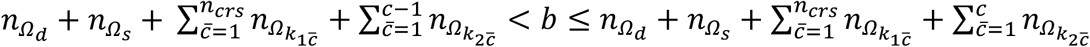,

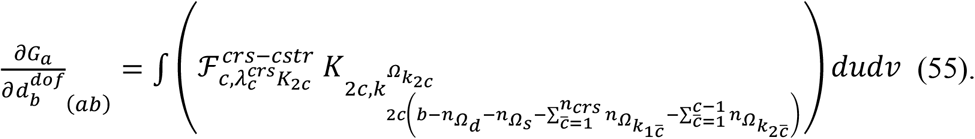

Note that the function evaluation and integration in Equations 36-55 can be performed locally only for corresponding elements since each element unknown weights the element shape function locally in finite element methods.

### Function evaluation for elements in the residual vector and the Jacobian matrix

To evaluate the residual vector and the Jacobian matrix, functions *X*^*e*^, *Y*^*e*^, *Z*^*e*^, *α*^*e*^, 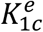, and 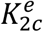 can be defined as given in Equations 56-61 from the element point of view.

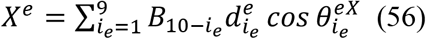

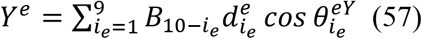

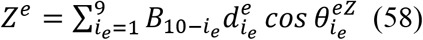

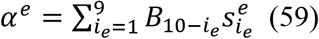

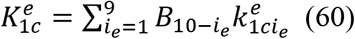

and

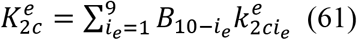

*B*_1_ to *B*_9_ are nine basis B-spline surface functions in the parent domain defined by [™1,1] and η[™1,1] (see Equations S31–S39 and Fig. S1D). 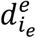 (Fig. S1B), 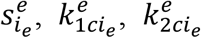 are unknowns. 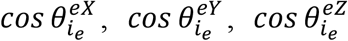 can be calculated from 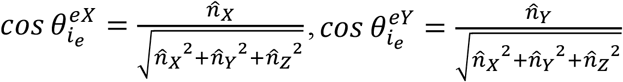, 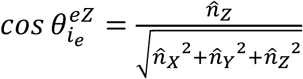, respectively, at the center of each element.

To perform the numerical integration, the 5 × 5 Gaussian rule was used. Therefore, the B-spline functions and their derivatives per element were priorly calculated at twenty-five Gauss quadrature points.

### Iterative calculations

With the given boundary values 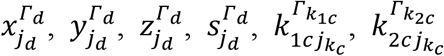 and the given fixed positions 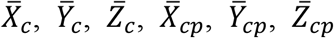 for crosslinkers, initial guesses for the unknowns 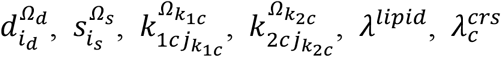 were substituted into the residual vector 𝒢 and the Jacobian matrix 𝒥. The vector 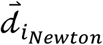 for the unknowns can be defined and calculated from 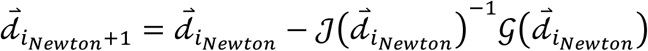, where *i*_*Newton*_ is the index for the iteration in the Newton-Raphson method. This iterative process was continued until Euclidean norms for the difference of two consecutive solutions for 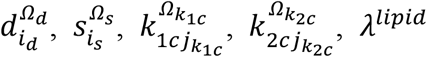 and 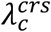 become smaller than predefined values.

### Forces applied on the fixed point for crosslinkers

Forces applied on the fixed point (Fig. 1 red bead) of crosslinkers can be defined by taking the derivative of the total energy with respect to 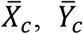, and 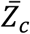 (or 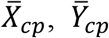, and 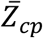)., *Y*, and *Z* components of the force on the crosslinker indexed with *c* can be expressed as follows.

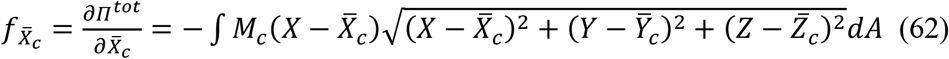

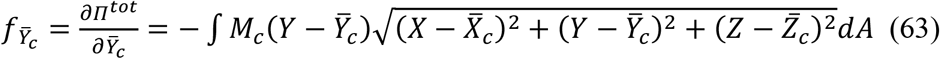

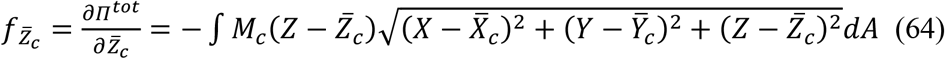

The total crosslinker force can be 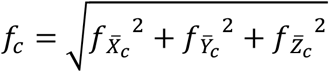. Calculated solutions from the finite element model can be substituted into Equations 62-64 to compute the force values. To compute 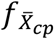 from Equation 62, 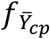 from Equation 63, 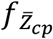 from Equation 64, and 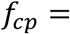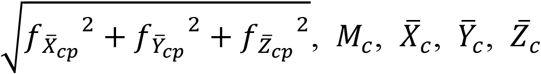 in Equations 62-64 can be replaced with 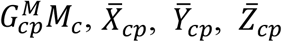, respectively.

### Programing and simulation environments

The model was coded by using MATLAB R2022a and its curve fitting and symbolic math toolboxes. The code was parallelized by using the parallel computing toolbox of MATLAB. A function central_diff.m (by Robert A. Canfield) from MATLAB Center File Exchange was also used for data analyses. The computation was conducted in the Windows 11 Home environment by using an Intel Core i9-10980XE processor and 128 GB usable memory.

## Acknowledgements

The author thanks Glenn Manarin (KAIST) for the proofreading.

## Code availability

MATLAB codes for the finite element model and the data analysis will be freely available upon publication in the journal.

## Data availability

Calculated raw data will be available upon publication in the journal.

## Supporting Information (SI) Supporting Equation

### Parametric derivatives

Parametric derivatives with respect to the parametric coordinates *u* and *v* are summarized below. The comma symbol indicates partial derivatives.

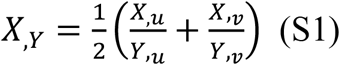

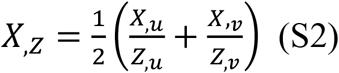

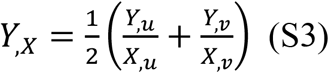

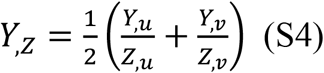

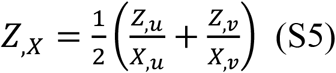

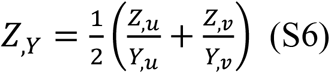

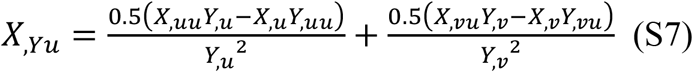

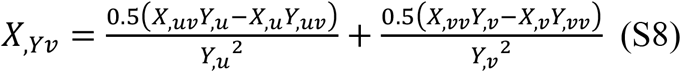

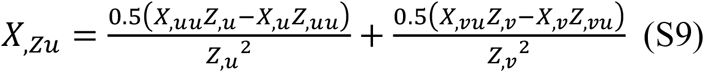

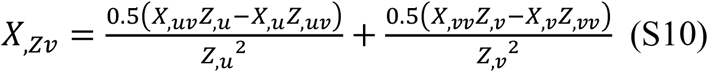

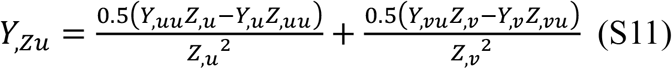

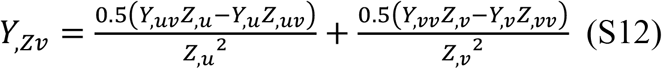

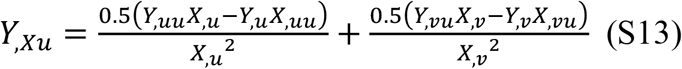

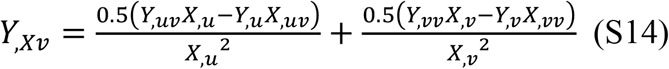

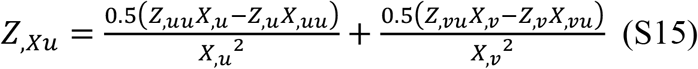

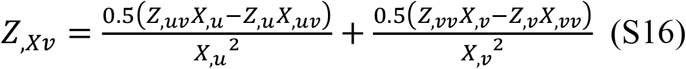

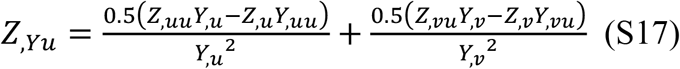

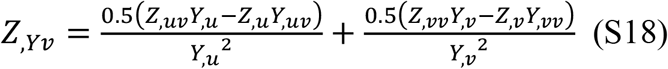

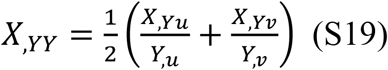

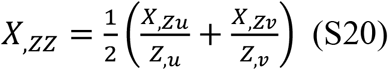

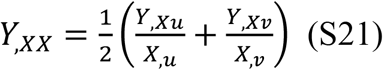

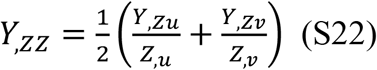

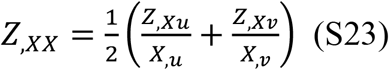

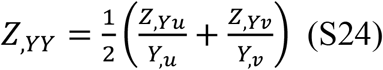

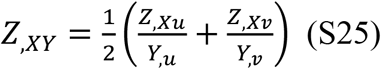

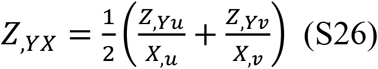

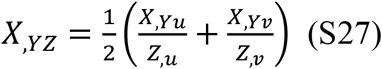

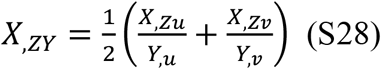

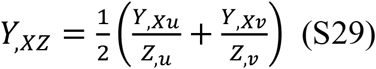

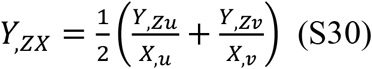

### B-spline surface functions used in this work for prior function evaluation at the Gauss quadrature points

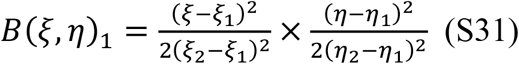

where (*ξ*_1_, *ξ*_2_) = (−1, 1) and (*η*_1_, *η*_2_) = (−1, 1).

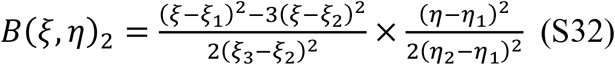

where (*ξ*_1_, *ξ*_2_, *ξ*_3_) = (−3, −1, 1) and (*η*_1_, *η*_2_) = (−1, 1).

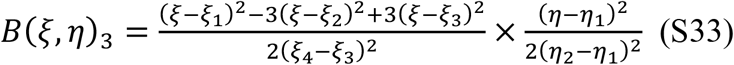

where (*ξ*_1_, *ξ*_2_, *ξ*_3_, *ξ*_4_) = (−5, −3, −1, 1) and (*η*_1_, *η*_2_) = (−1, 1).

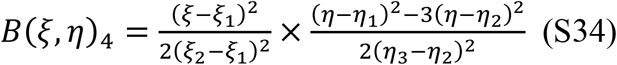

where (*ξ*_1_, *ξ*_2_) = (−1, 1) and (*η*_1_, *η*_2_, *η*_3_) = (−3, −1, 1).

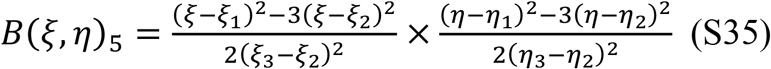

where (*ξ*_1_, *ξ*_2_, *ξ*_3_) = (−3, −1, 1) and (*η*_1_, *η*_2_, *η*_3_) = (−3, −1, 1).

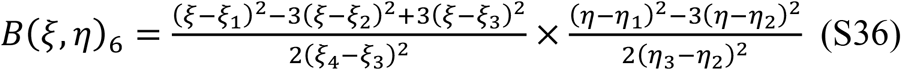

where (*ξ*_1_, *ξ*_2_, *ξ*_3_, *ξ*_4_) = (−5, −3, −1, 1) and (*η*_1_, *η*_2_, *η*_3_) = (−3, −1, 1).

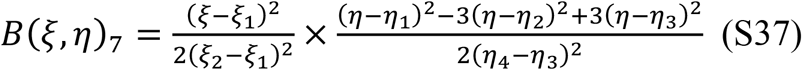

where (*ξ*_1_, *ξ*_2_) = (−1, 1) and (*η*_1_, *η*_2_, *η*_3_, *η*_4_) = (−5, −3, −1, 1).

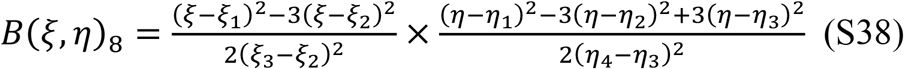

where (*ξ*_1_, *ξ*_2_, *ξ*_3_) = (−3, −1, 1) and (*η*_1_, *η*_2_, *η*_3_, *η*_4_) = (−5, −3, −1, 1).

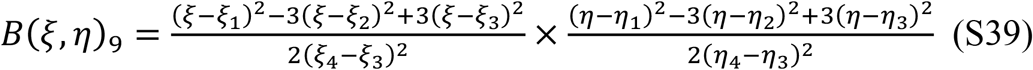

where (*ξ*_1_, *ξ*_2_, *ξ*_3_, *ξ*_4_) = (−5, −3, −1, 1) and (*η*_1_, *η*_2_, *η*_3_, *η*_4_) = (−5, −3 − 1, 1).

### Expansion for the element of the residual force vector

Equation 29 can be expanded as follows.

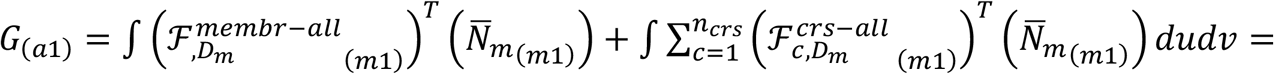

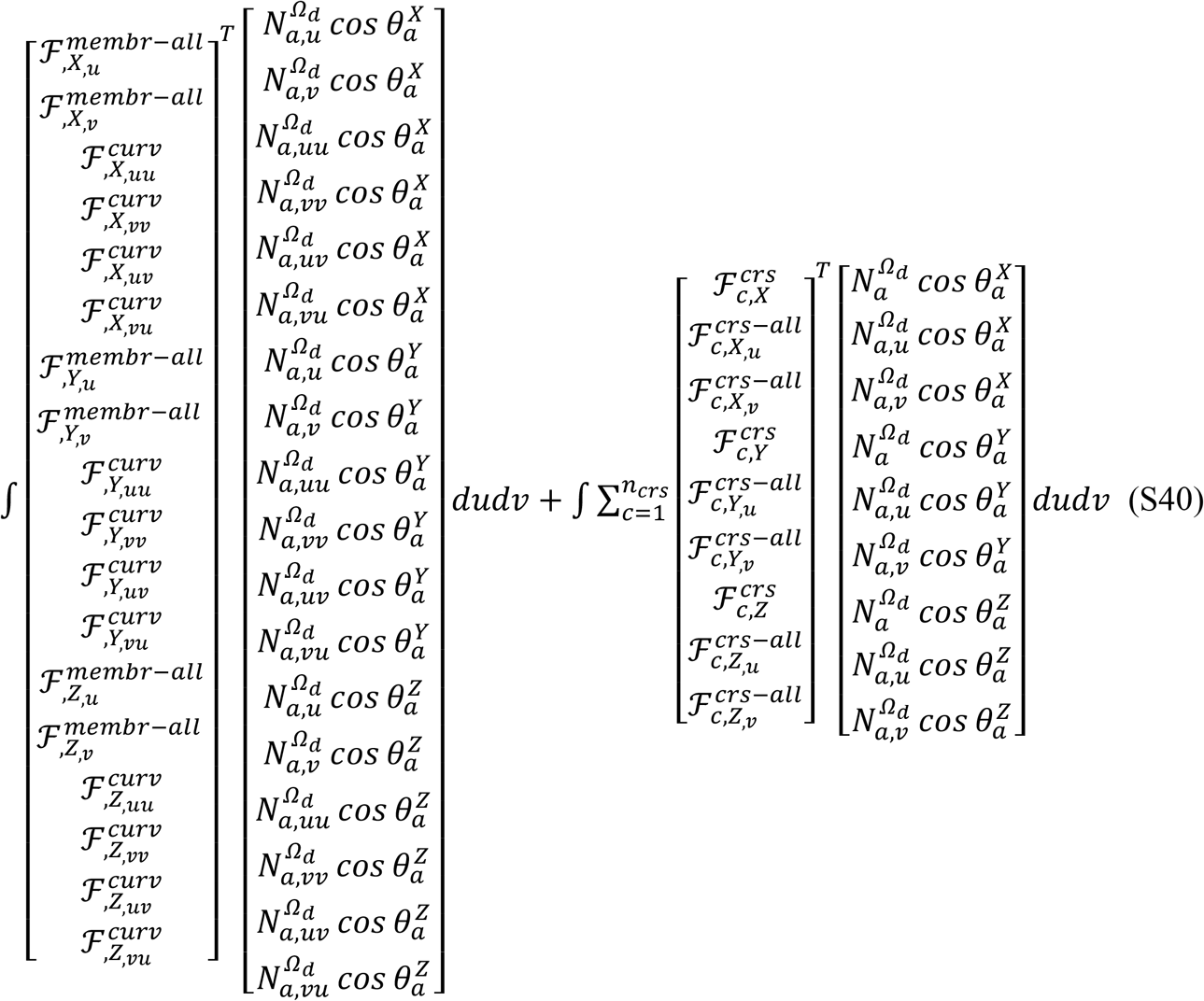

### Expansion for the element of the Jacobian matrix

Equation 36 can be expanded as follows.

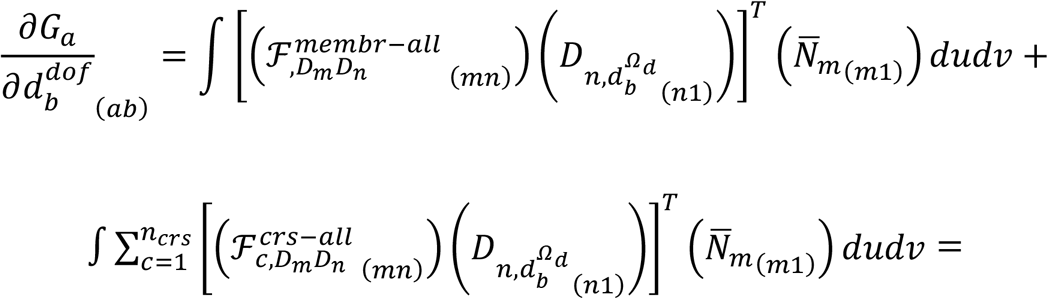

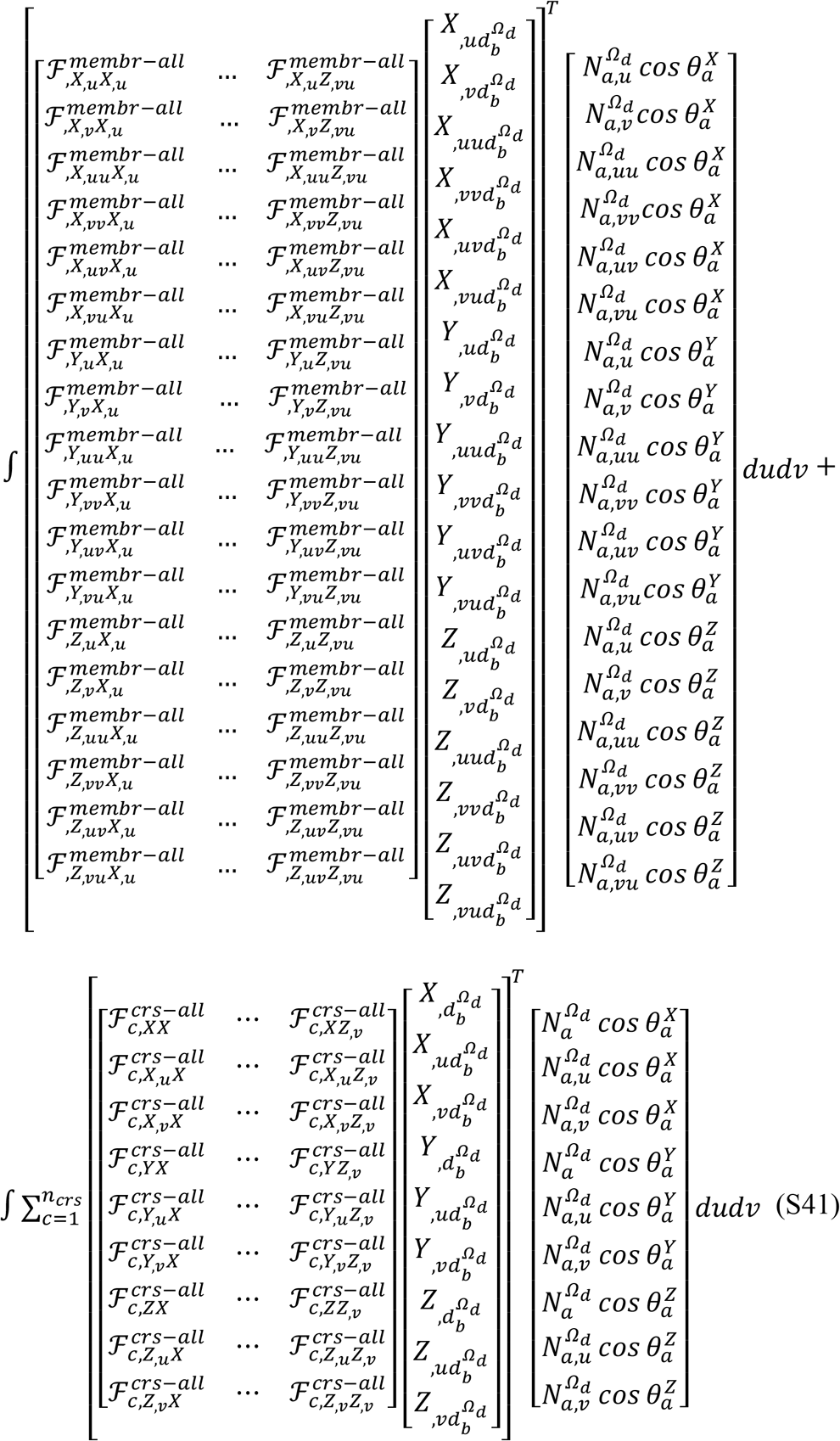

Equation 37 can be expanded as follows.

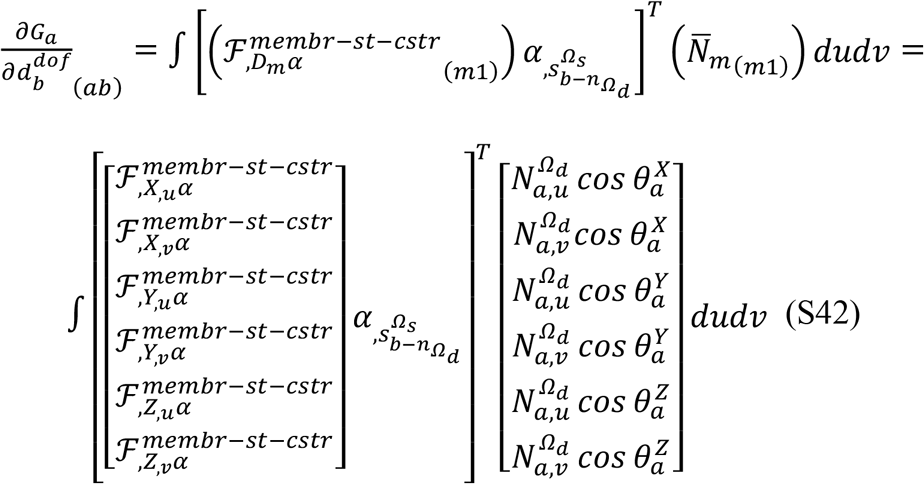

Equation 38 can be further expanded as follows.

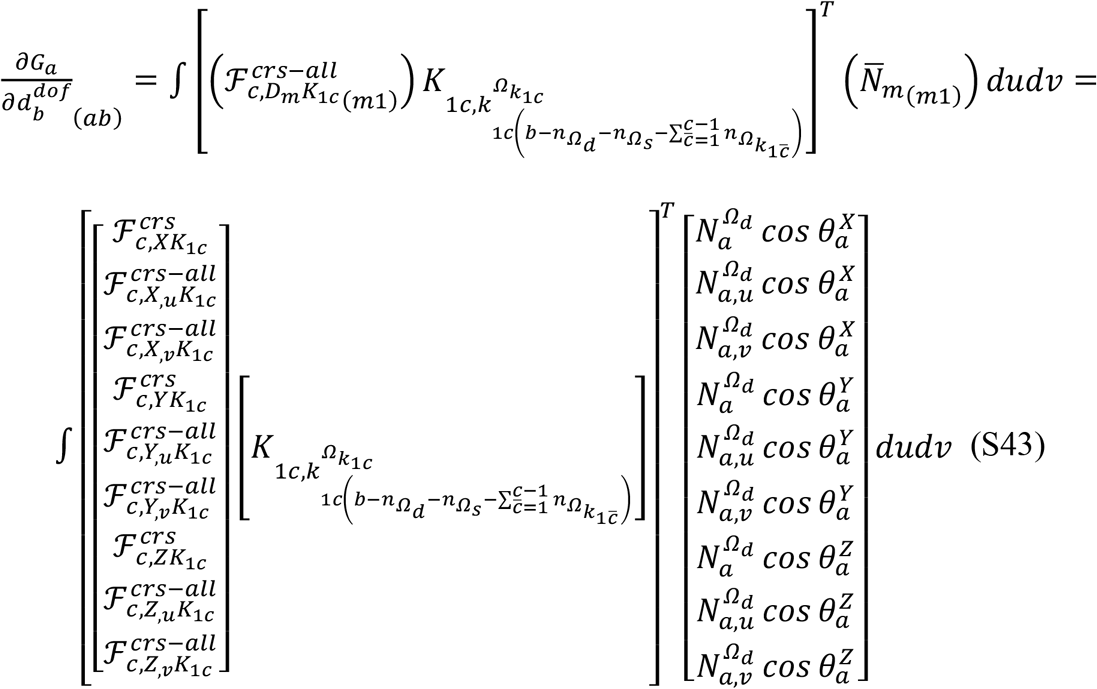

Equation 39 can be further expanded as follows.

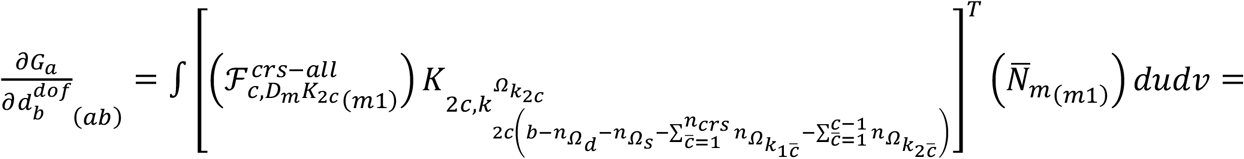

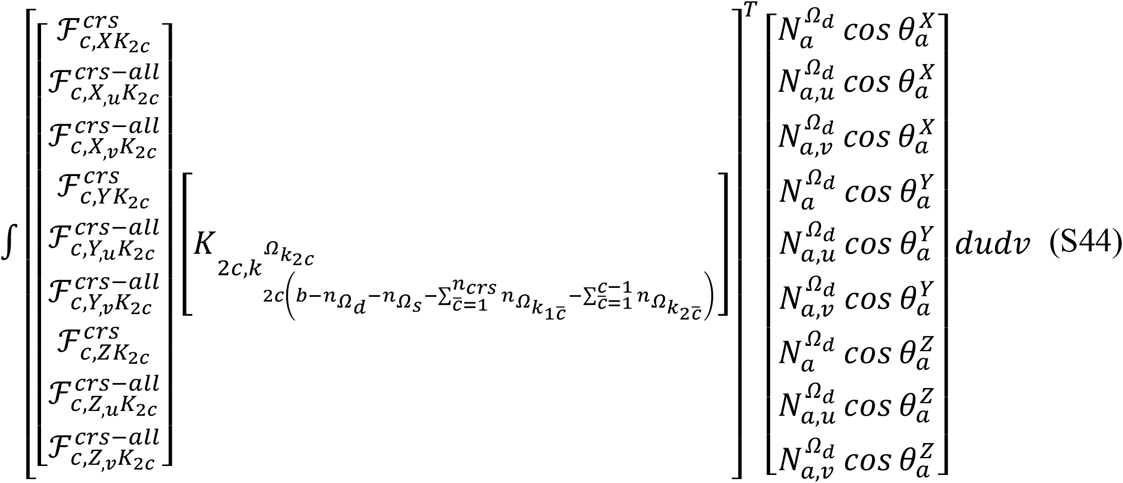

Equation 40 can be expanded as follows.

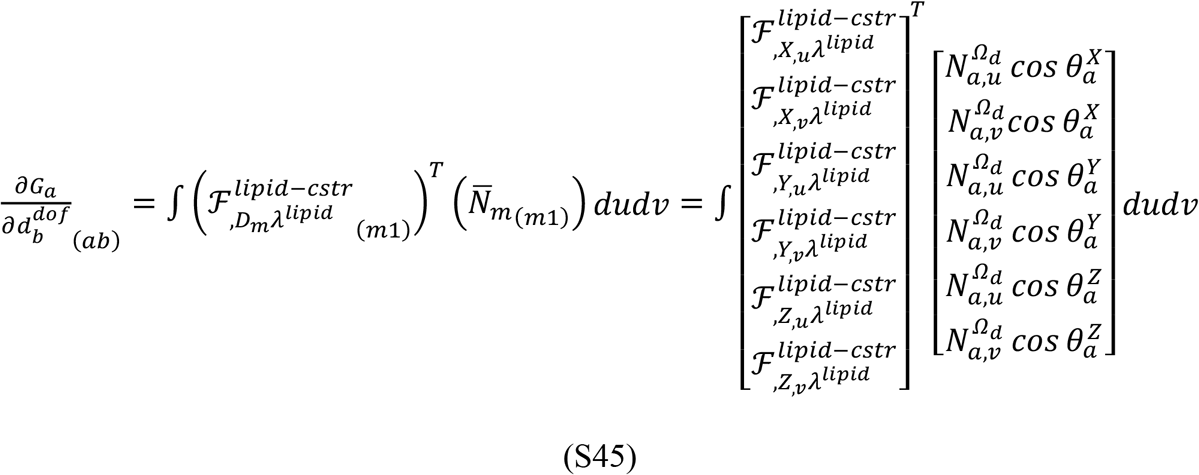

Equation 41 can be expanded as follows.

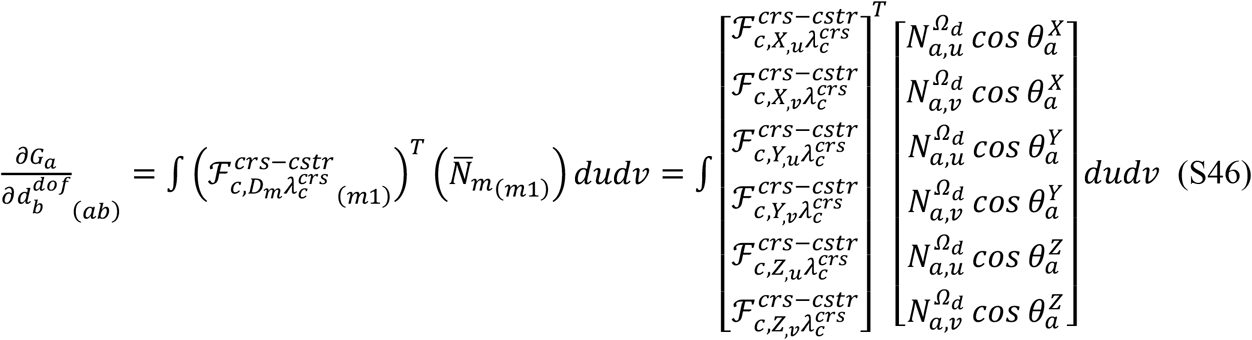

Equation 42 can be expanded as follows.

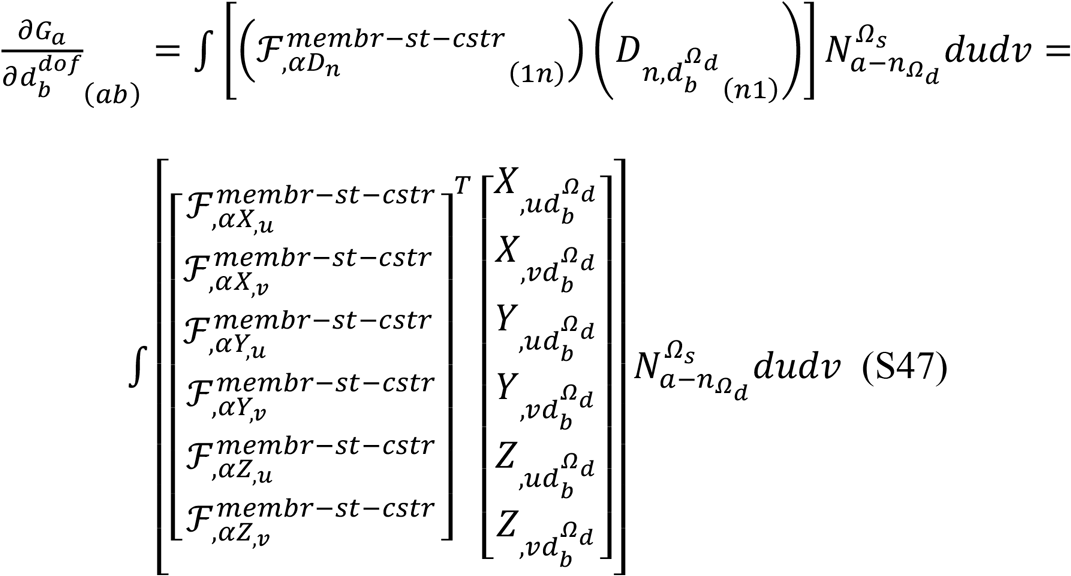

Equation 45 can be expanded as follows.

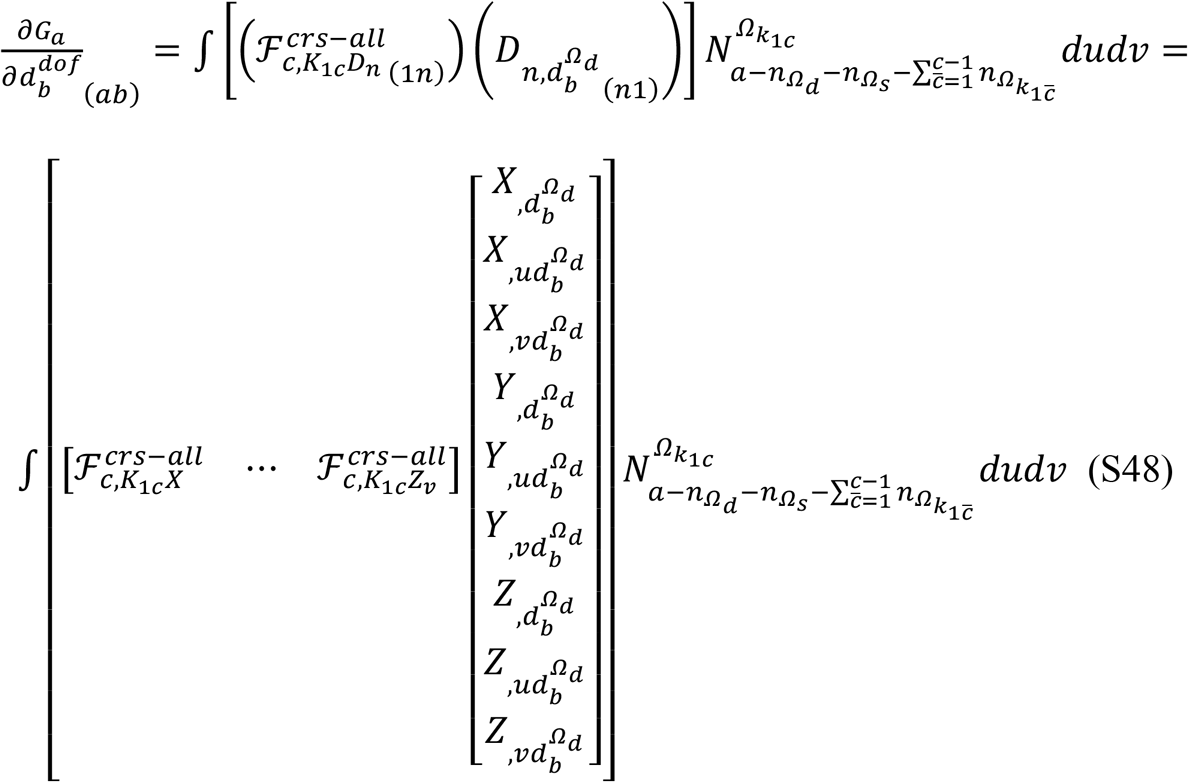

Equation 48 can be expanded as follows.

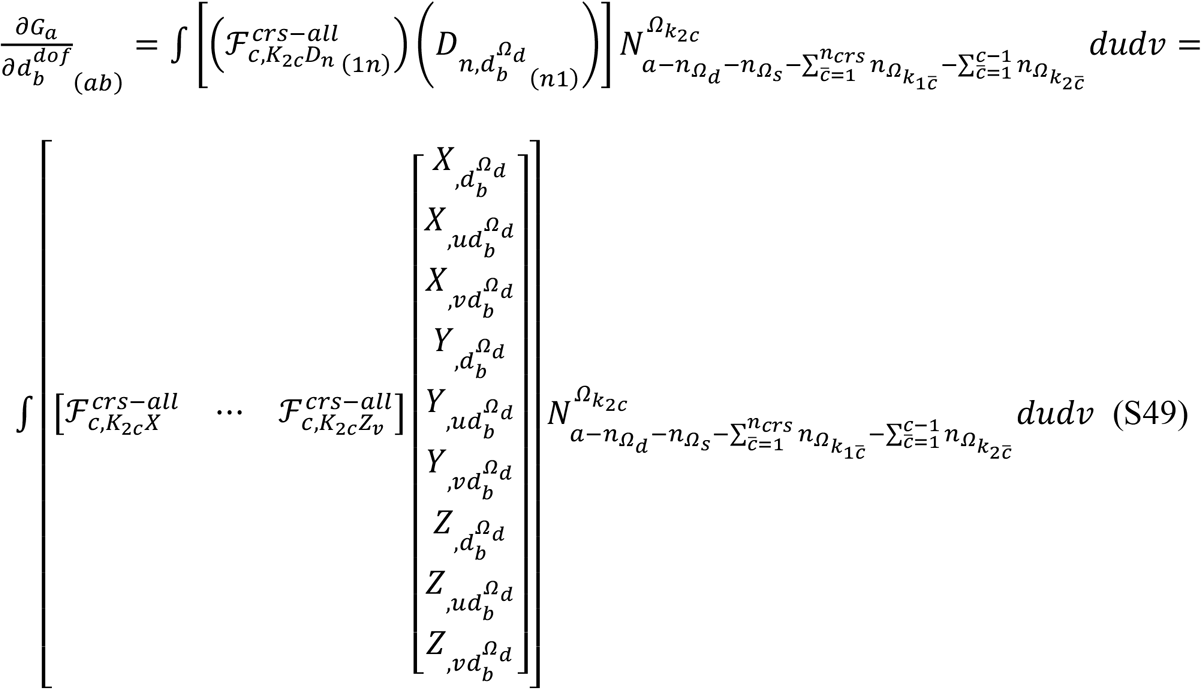

Equation 51 can be expanded as follows.

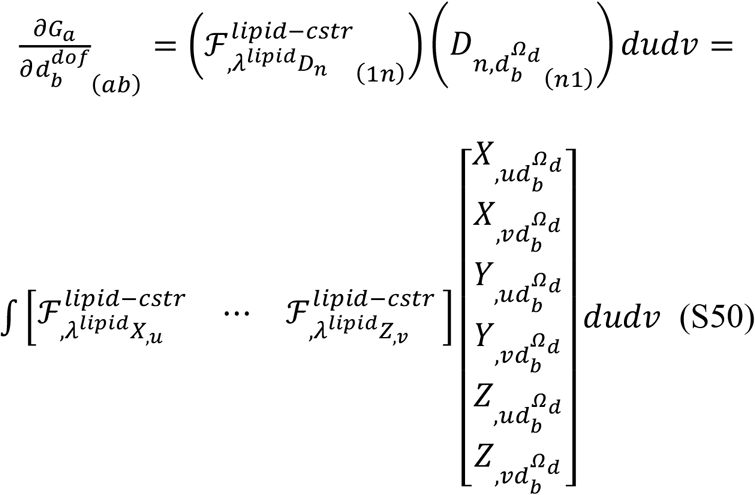

Equation 53 can be expanded as follows.

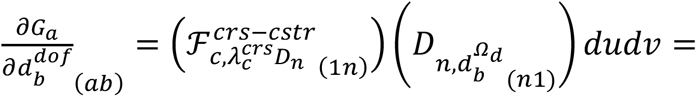

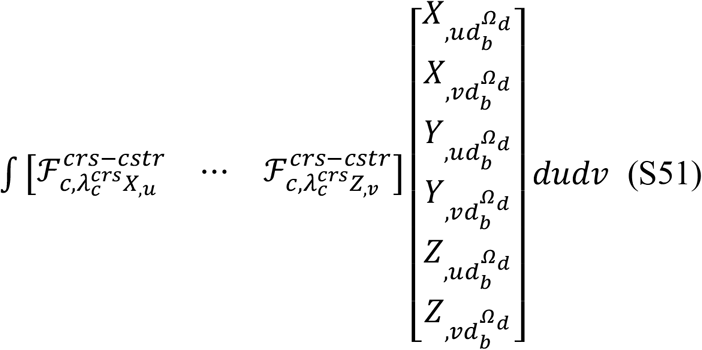

### Derivation of Equation 7 in the main text

From Equation 5, the elastic energy for talin with two fixed ends can be written as follows.

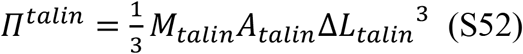

*M*_*talin*_ and *A*_*talin*_ *are t*he elastic modulus density of talin and the area of the fixed region of talin, respectively. Since the fixed end can be considered as a point, φ = (*K*_1_ + *K*_2_)_*talin*_ *can b*e held. To obtain the minimum П^*talin*^ *by ha*ving the minimum *M*_*tali*_, φ *=* (*K*_1_ + *K*_2_)_*talin*_ *= 2K*_1_*A*_*talin*_ *can b*e held. Therefore, *M*_*talin*_ *can b*e written as follows.

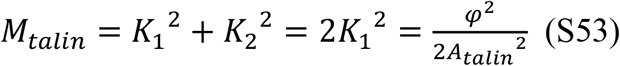

By substituting Equation S53 into Equation S52 and by taking the derivative of Equation S52 with respect to Δ*L*_*talin*_, *the* force for the fixed talin can be defined as follows.

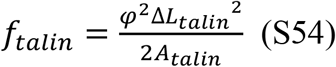

## Supporting Notes

### Lipid sorting simulation

According to previous works, a variety of different lipid species and classes can collectively form cellular membranes. For example, a lipidomic study performed for human islets detected 329 lipid species across eight major lipid classes (49). A recent study on single cell lipid profiling identified 81 species from six major lipid classes (50). Therefore, eight lipids with different bending moduli were considered for the calculations in Fig. 2.

Regularly defined nine crosslinkers in a 1.547 × 1.547 μm^2^ planar membrane were anti parallelly stimulated, as shown in Fig. 2A and B. In this calculation, the displacement of elements in the boundary region of the square membrane was also prescribed, as shown in Fig. 2A, by sampling calculated values from the interior region (see Fig. S15 for the indication of the boundary region).

To implement lipid sorting in the membrane, the combined membrane energy density for mean curvatures and lateral strain values of each element was calculated at the end of each calculation performed by prescribing the displacement. The Gaussian curvature energy density was not considered in this process based on the Gauss-Bonnet theorem. With the given energy densities, a lower bending modulus was assigned to the element with a higher energy density value. By counting the number of cumulated lipids assigned for certain elements, the process was repeated and continued until all elements for the membrane are assigned with a bending modulus value. This set of processes was repeated for each step performed by varying the displacement.

The described lipid sorting process was identically applied for the calculations in Fig. 4.

### Segmentation algorithm to define lipid nanodomains for calculations in Fig. 2

A segmentation algorithm was implemented to define lipid nanodomains in the finite element model framework after lipid sorting. With a given tetrahedral element in the membrane, four adjacent elements that share an edge with the given center element were evaluated with respect to whether their bending moduli are the same as the bending modulus of the center element. This process was repeated with the growth of a nanodomain and completed when no adjacent element was found with the same bending modulus. Here, elements with a share vertex (without a shared edge) were not included in the same lipid nanodomain. Note that the nanodomains identified with the most outer boundary elements of the square membrane were not considered in calculating mean areas in Fig. 2O because of the possibility that the nanodomains can continue beyond the boundary. MATLAB codes are available for the segmentation analysis.

### The half max *M*_*c*_ area

To estimate the area for the confined Brownian motion of crosslinkers, the area defined by half of the maximum *M*_*c*_ was calculated. For the physical domain of the membrane made by (*n*_*grid_u*_ *−* 2) × (*n*_*grid_v*_ *−* 2) elements, where *n*_*grid_u*_ and *n*_*grid_v*_ are the number of grid regions on parametric coordinate *u* and *v*, respectively, *M*_*c*_ values at 4(*n*_*grid_u*_ *−* 2)(*n*_*grid_v*_ *−* 2) + 2(*n*_*grid_u*_ *−* 2) + 2(*n*_*grid_v*_ *−* 2) + 1 number of regularly defined points in the membrane were tested with respect to whether they are equal or greater than the half max *M*_*c*_ value. From the sampled points, the area was calculated by using alphaShape and surfaceArea functions in MATLAB.

### Remeshing process for the finite element model

For all calculations presented in this article, unless mentioned otherwise, the regulation of mesh grids was conducted for the solution of the previous step before using that as the initial guess in the Newton-Raphson method. A simple algorithm was implemented as follows. First, by using the calculated membrane-crosslinker solution, all membrane arclengths on the parametric mesh grid to the direction of *u* were uniformly discretized. Second, this process was repeated for the arclengths on the mesh grid to the direction of *v*. Third, the control point values, i.e. the weights of the B-spline function, for the displacement functions *X, Y*, and *Z* were adjusted until the corresponding membrane mesh became close enough to the uniformly regulated membrane mesh. This was implemented by comparing the adjusted *X, Y*, and *Z* values at the center of elements and the uniformly regulated point values. By doing this, 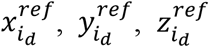 in Equations 16-18 can be determined. The same process was repeated for *α, K*_1c_, and *K*_2c_.

For all data in this work, the first three displacement steps were calculated without the remeshing procedure. Additionally, data between the 34^th^ point and the 61^st^ point in Fig. 4F blue trace were obtained without the remeshing process because the Newton-Raphson method did not provide a converged solution, perhaps due to poor initial guesses after the remeshing process. Also, the remeshing was omitted as it was not relevant for all calculations in Fig. S2.

### Calculations for the lipid number strain *α* around membrane-inserted proteins

A set of calculations was performed to test whether non-negligible gradients of lipid number strain *α* can be formed around membrane-inserted proteins. The strain *α* was calculated for a rectangular membrane where eight membrane proteins are considered, as shown in Fig. S2. To define the protein insertion points, two-by-two elements were selected to generate the zero-strain condition at their shared vertex (see Fig. S2C). To increase the strain in the rectangular region, the total number of lipids (*n*_*lipid*_ in Equation 1) was reduced. The strain values for the boundary of the rectangular membrane were also prescribed (see Fig. S2C).

Two different inter-protein distances were tested. The distance was about 0.9 nm and 0.9 Å in Fig. S2A and B, respectively. The number of elements was the same for both cases. A steeper gradient was generated with the reduced inter-protein distance. According to data in Fig. S2B, the length scale of the gradient can be smaller than 1 Å. The results in Fig. S2 allowed us to assume that the strain gradient around the membrane-inserted protein can be negligible.

## Supporting Figures

**Fig. S1.**
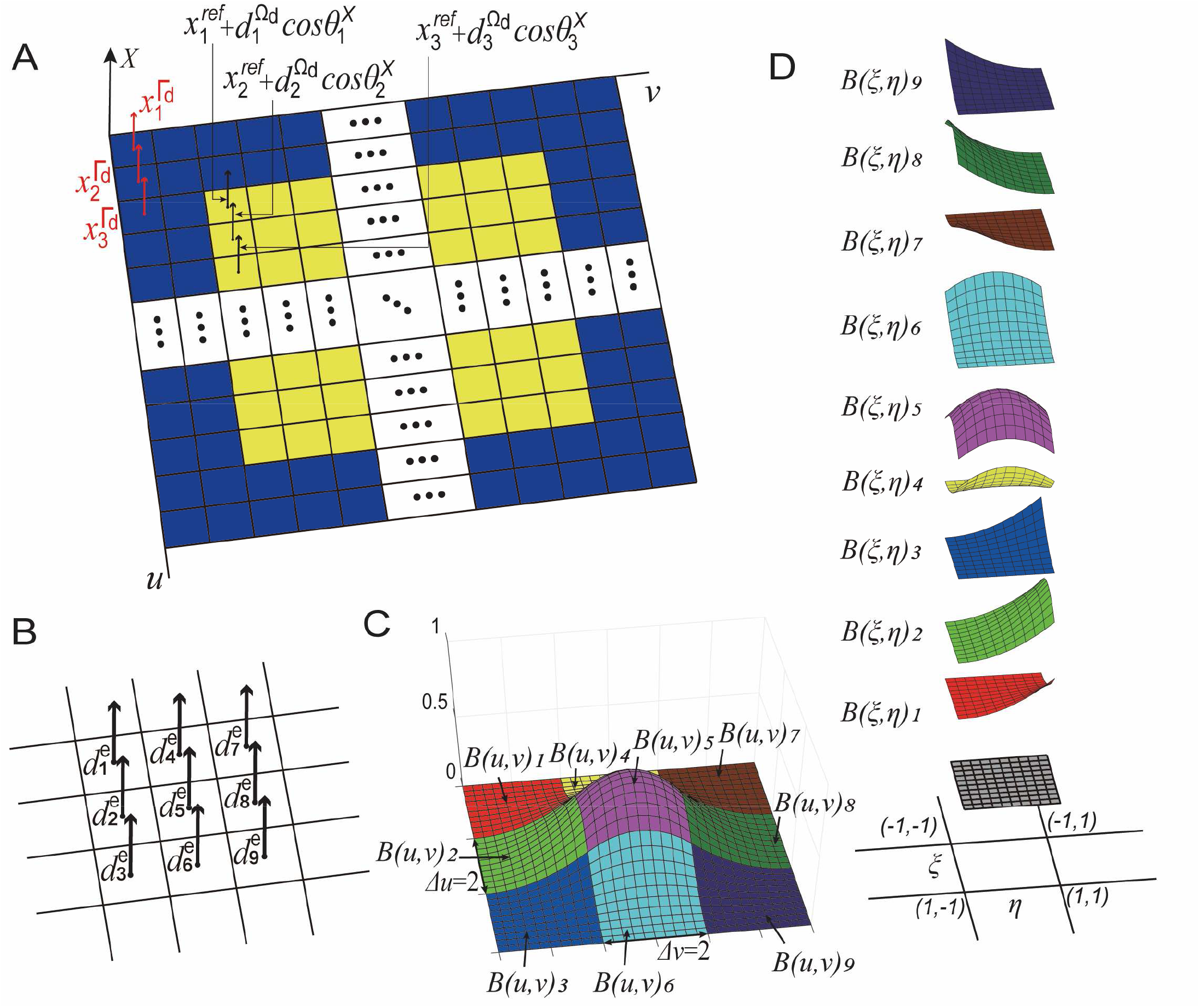
Description of the finite element model. (**A**) Parametric coordinates *u* and *v* defined for the displacement function *X*. See Equation 16 for how weights shown with black and red arrows are applied for the B-spline shape functions. The domain for the fixed boundary values (*Γ*_*d*_) is shown in navy blue. The domain for the unknown variables (*Ω*) is shown in yellow. The displacement functions *Y* and *Z*; the function for the lateral strain *α*; and the functions *K*_1*c*_ and *K*_2*c*_ can be defined similarly. (**B**) Nine unknowns for *X, Y*, and *Z* of an element are shown from the element point of view. The B-spline functions used in this work require nine control point weights to define the local function values for *X, Y, Z, α, K*_1*c*_, and *K*_2*c*_. (**C**) The B-spline basis functions used in this model. (**D**) The nine basis functions in the parent domain defined by *ξ*[−1,1] and **η**[−1,1] (See Equations S31-S39).

**Fig. S2.**
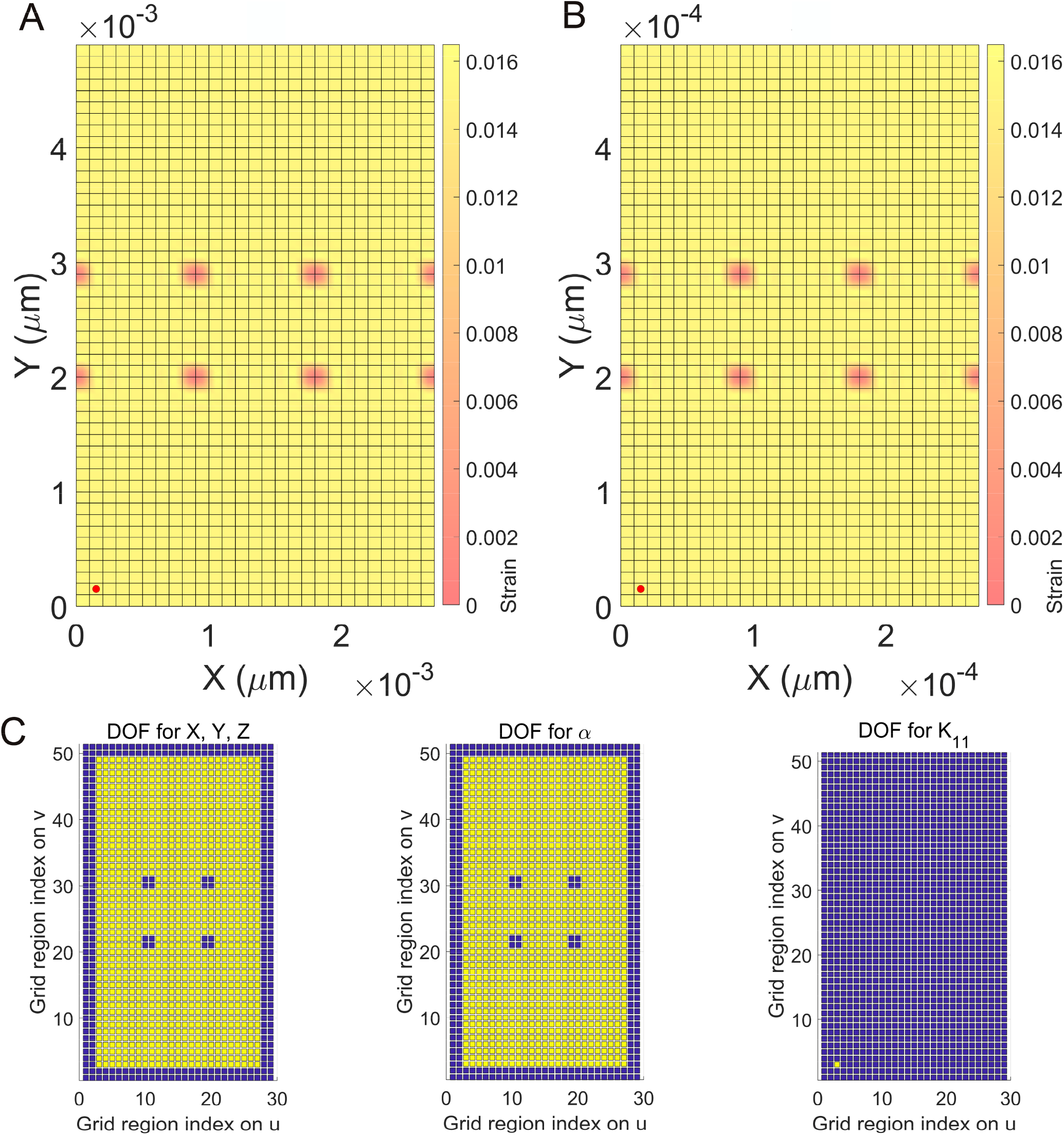
The profile of the lipid number strain around membrane-inserted proteins. Eight points (vertices) were assumed for the membrane-inserted proteins where the lateral strain *α* is zero. Calculations were performed by reducing the *n*_*lipid*_ value. (**A**) A calculation for the lateral strain profile when the protein-protein distance is about 0.9 nm. (**B**) Another calculation for the strain profile when the protein-protein distance is about 0.9 Å. The number of total elements is the same for (A) and (B). A crosslinker with one element for DOFs and a negligible *φ* value (*φ* = 2 × 10^−25^) was considered at the bottom left corner due to a programming issue. (**C**) Indication of DOFs for each element. Yellow: regions with unknowns. Navy blue: regions with fixed boundary values. The plot for *K*_21_ is identical to the plot for *K*_11_ in (C right).

**Fig. S3.**
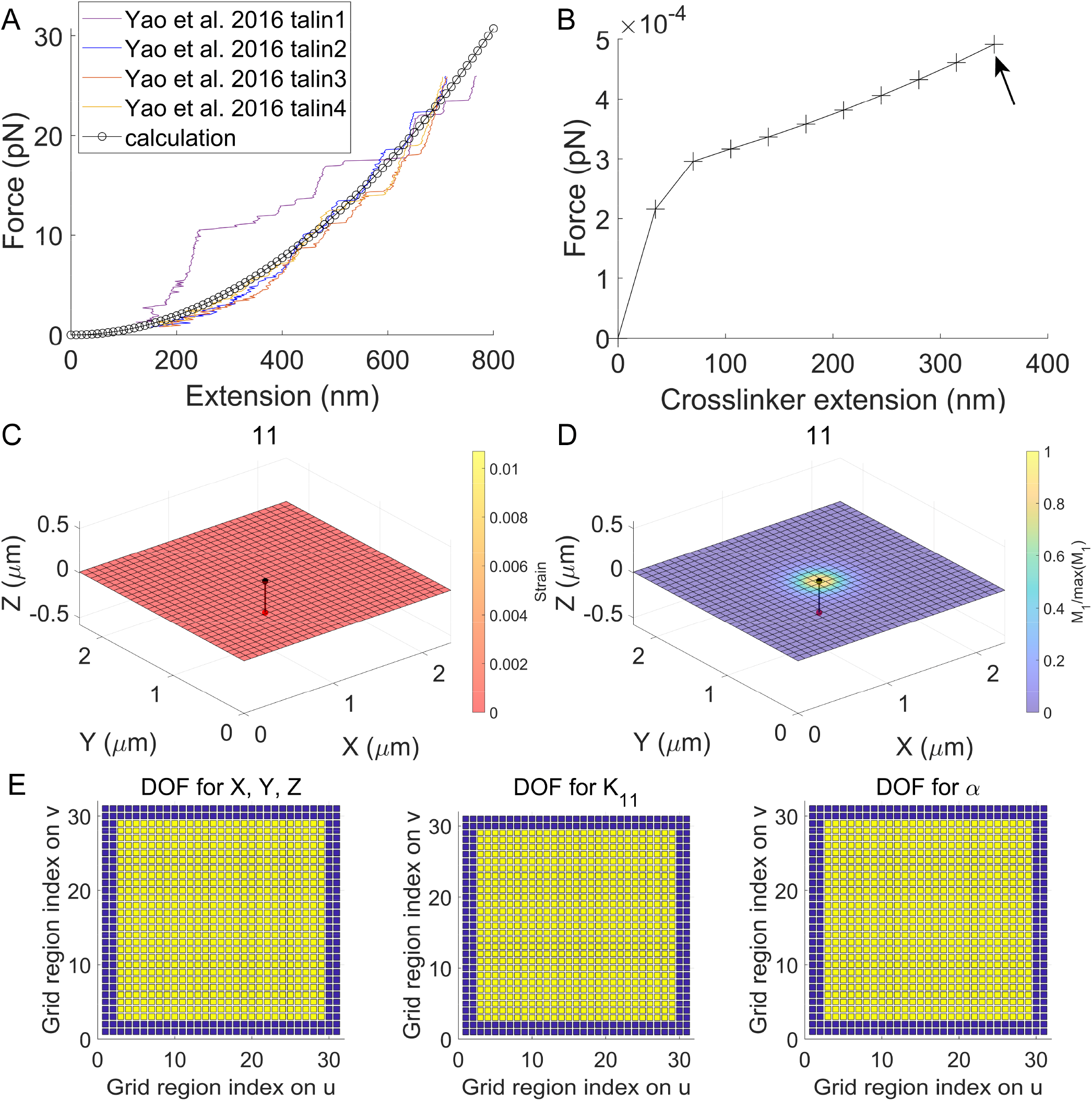
Different crosslinker responses depending on the mobility condition. (**A**) Comparisons between magnetic tweezer measurements (23) and calculations for the force vs. extension response of single full length talin monomers interacting with vinculin molecules in the solution. Two ends of the talin monomer are assumed to be fixed in the calculation. 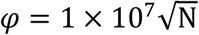 and A_talin_ = 10.2 nm^2^ were assumed to use Equation 7. (**B-D**) Calculations for the crosslinker where one end is mobile on the membrane surface. 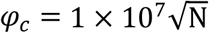 was also used. The extension of the crosslinker was defined from the maximum *M*_*c*_ point (black bead) to the fixed point (red bead). Membrane-crosslinker configurations for the final data point (arrow) are shown in (C, D). (**E**) Indication of DOFs for each element. Yellow: regions with unknowns. Navy blue: regions with fixed boundary values. The plot for *K*_21_ is identical to the plot for *K*_11_.

**Fig. S4.**
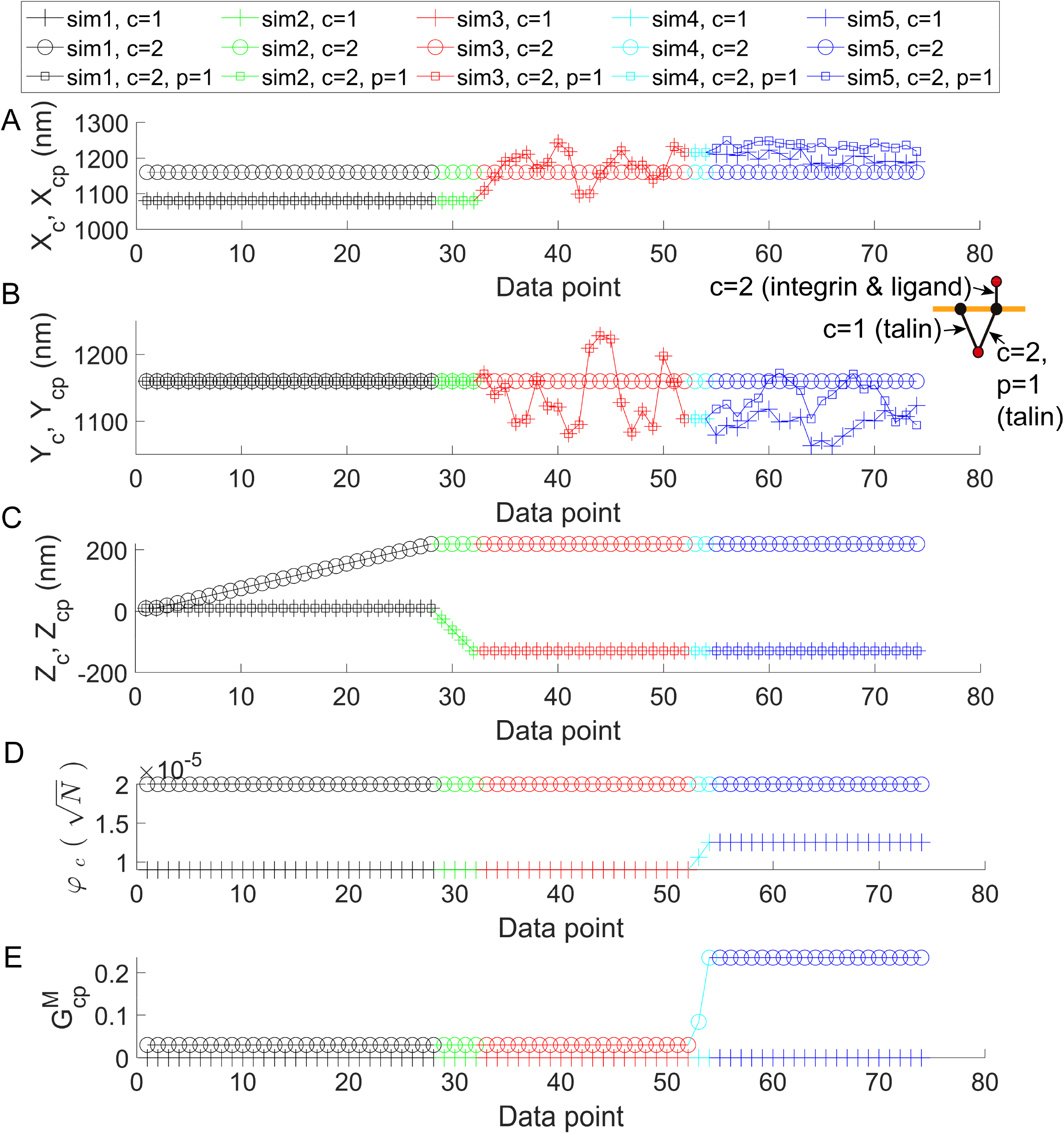
Inputs for calculations in Fig. 3. (**A-E**) Inputs for a set of five simulations in Fig. 3. Different simulations were plotted by using different colors.

**Fig. S5.**
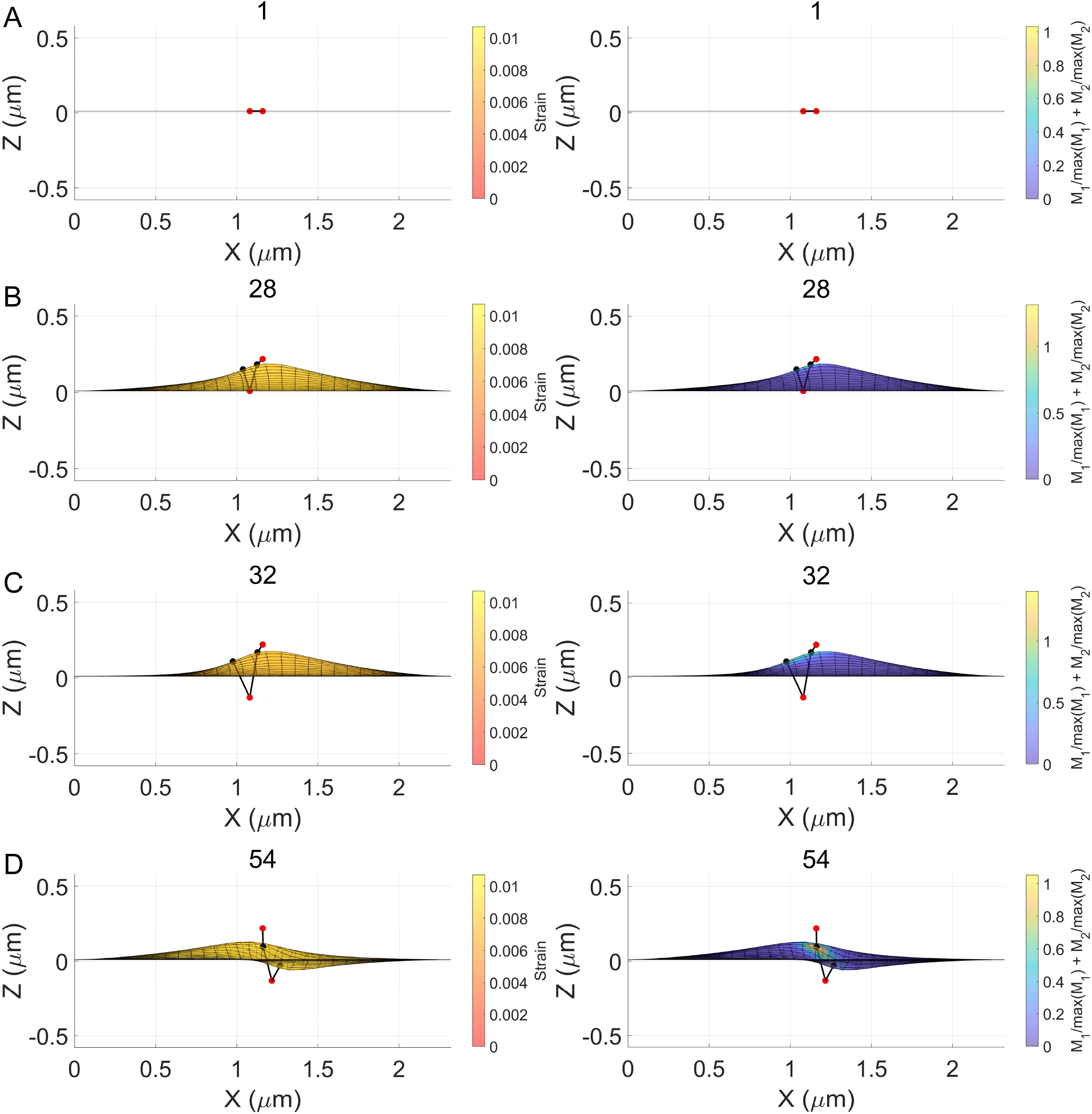
The side view of the membrane-crosslinker complex in Fig. 3. The number on top of each plot is the index for data points.

**Fig. S6.**
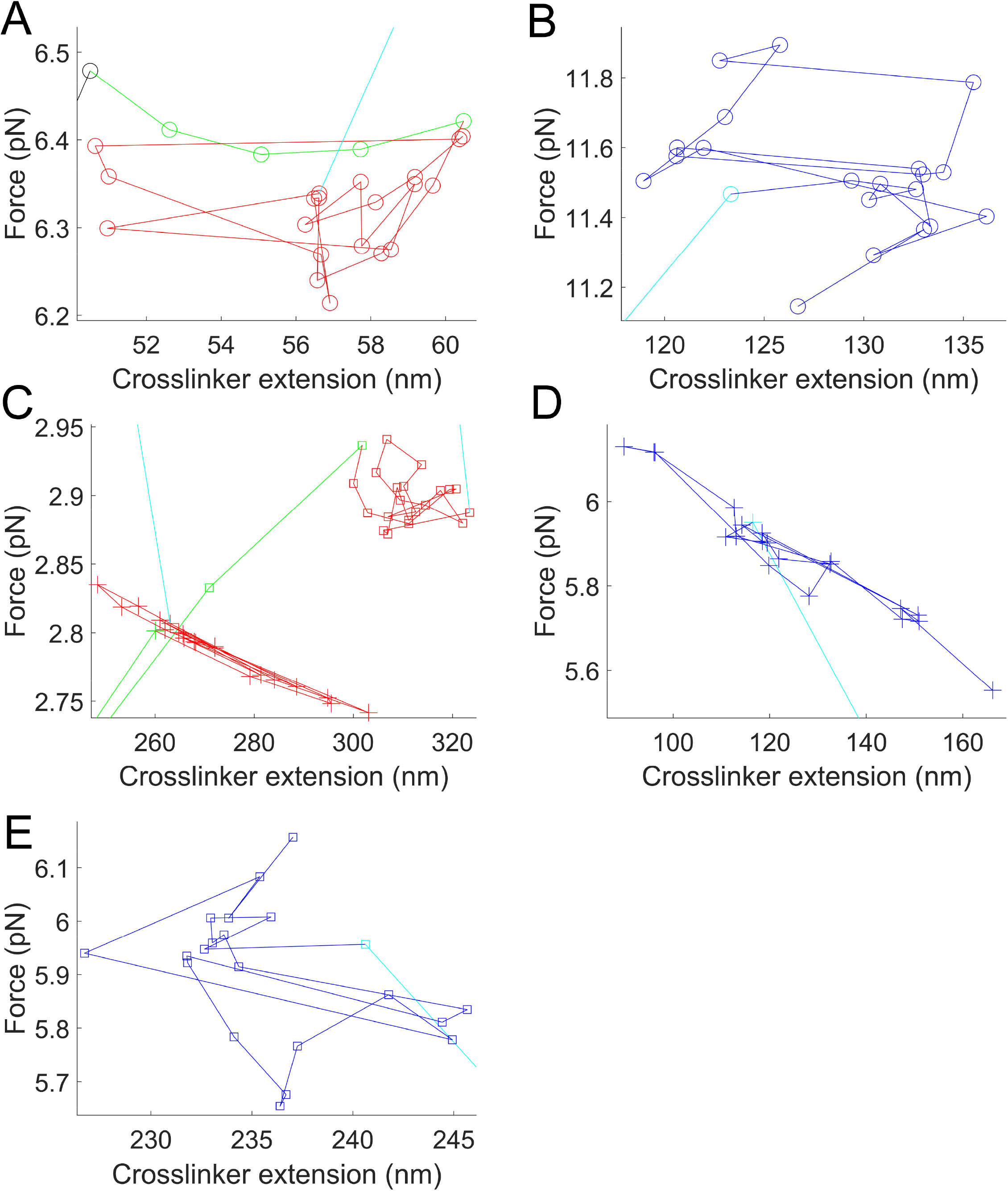
Expanded view for marked regions in Fig. 3A.

**Fig. S7.**
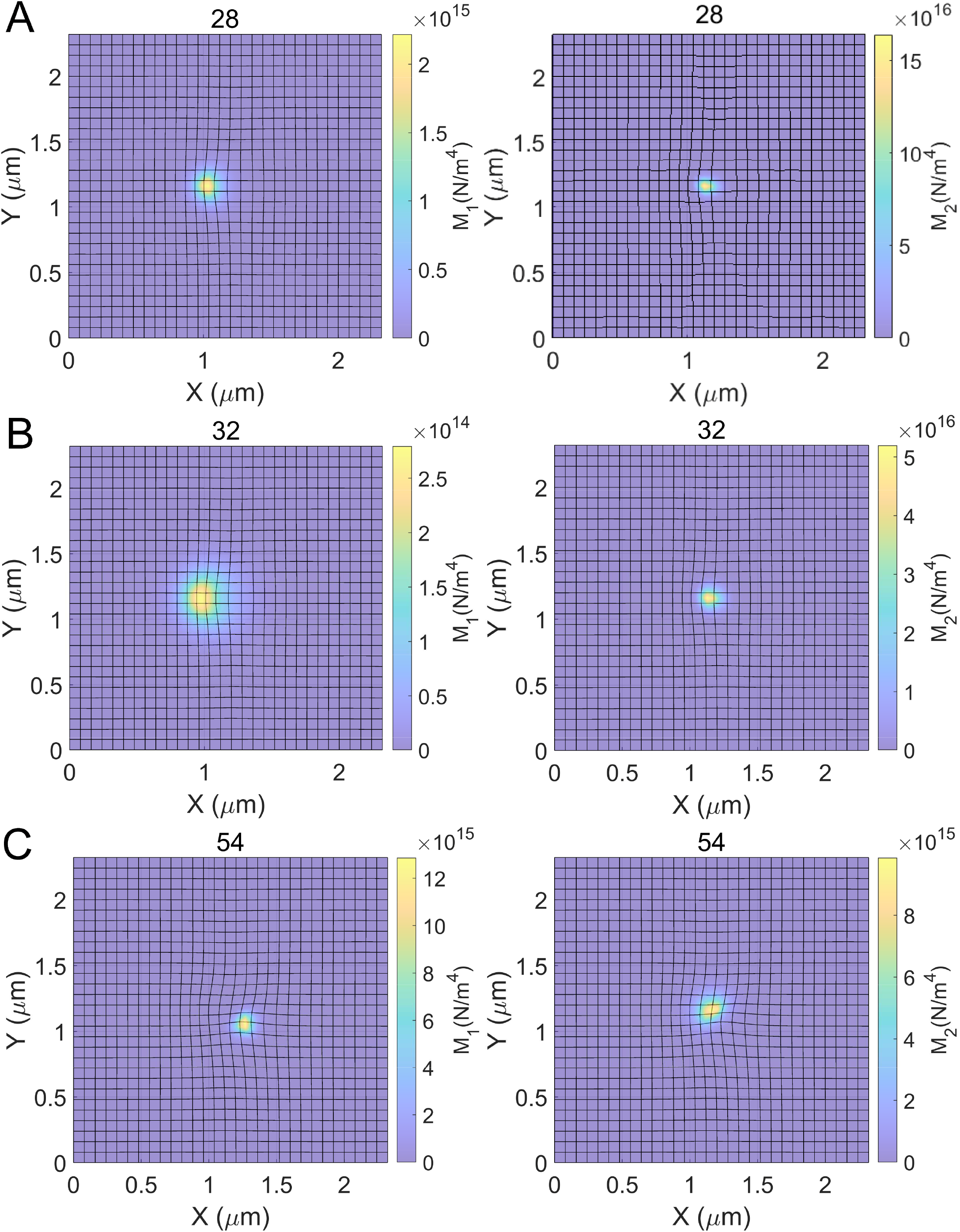
Top view of the membrane-crosslinker complex in Fig. 3 with the surface map of the elastic modulus density. The number on top of each plot is the index for data in Fig. 3. The elastic modulus density for (*c, p*) = (2,1) can be simply 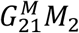.

**Fig. S8.**
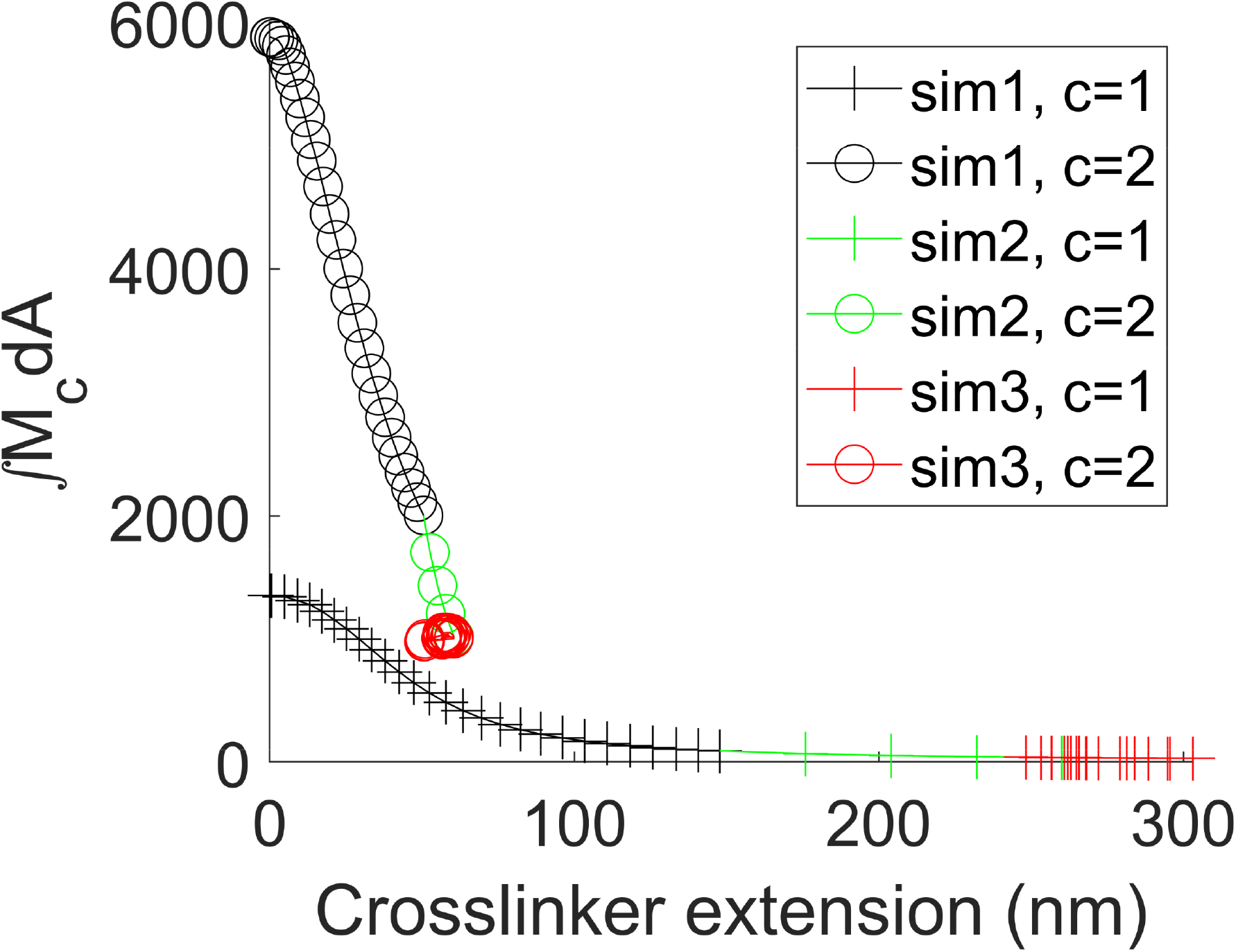
Total elastic modulus density ∫ *M*_*c*_*dA* for sim1-3 in Fig. 3. 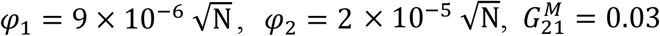.

**Fig. S9.**
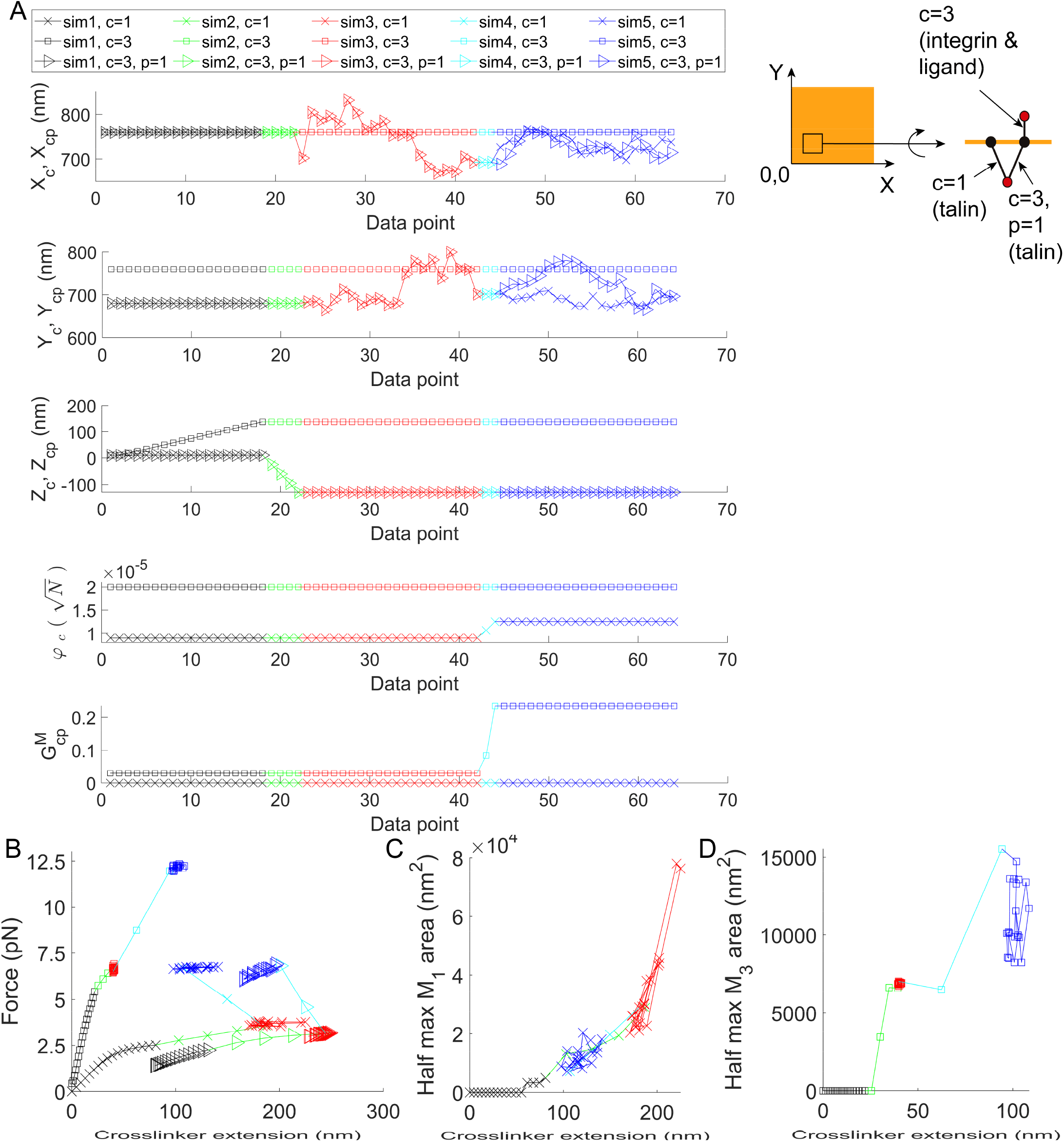
Responses of crosslinkers by considering four integrin-adhesion sites. See Movie S3 for all membrane-crosslinker configurations. The initial size of the membrane as well as parameter values for the membrane, talin, and the integrin-ligand complex are identical to those used in Fig. 3. Responses of crosslinkers at the bottom left corner of the membrane are presented (see inset and Movie S3). (**A-D**) Inputs (A) and responses (B-D) of crosslinkers with *c* = 1 (talin), *c* = 3 (integrin-ligand complex), and (*c, p*) = (3,1) (talin). See Fig. S11 for the indication of DOFs.

**Fig. S10.**
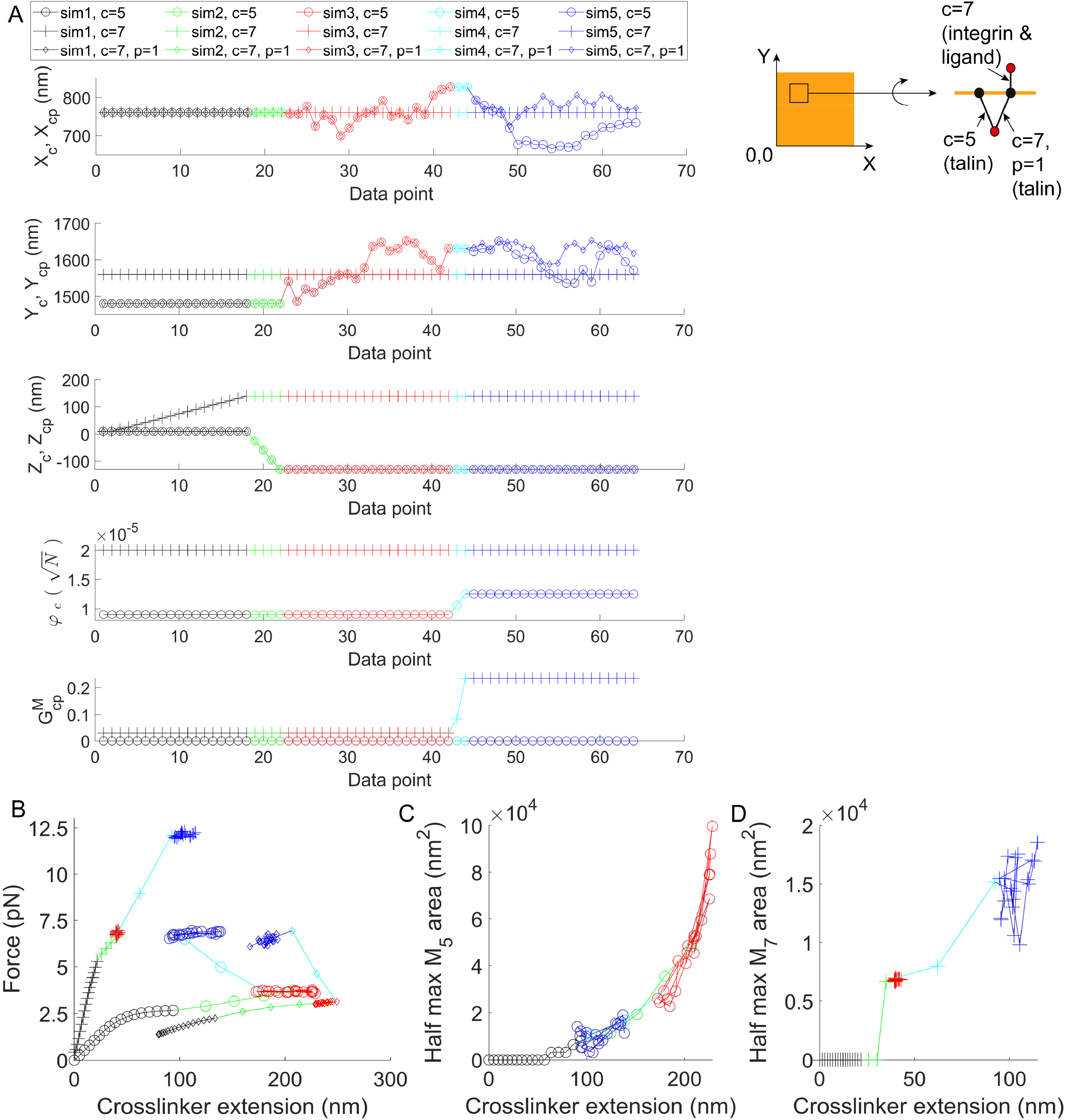
Responses of other crosslinkers in the membrane-crosslinker complex for Fig. S9 and Movie S3. As a continuation of the analyses in Fig. S9, responses of crosslinkers at the top left corner of the membrane are presented in this figure (see inset and Movie S3). (**A-D**) Inputs (A) and responses (B-D) of crosslinkers with *c* = 5 (talin), *c* = 7 (integrin-ligand complex), and (*c, p*) = (7,1) (talin).

**Fig. S11.**
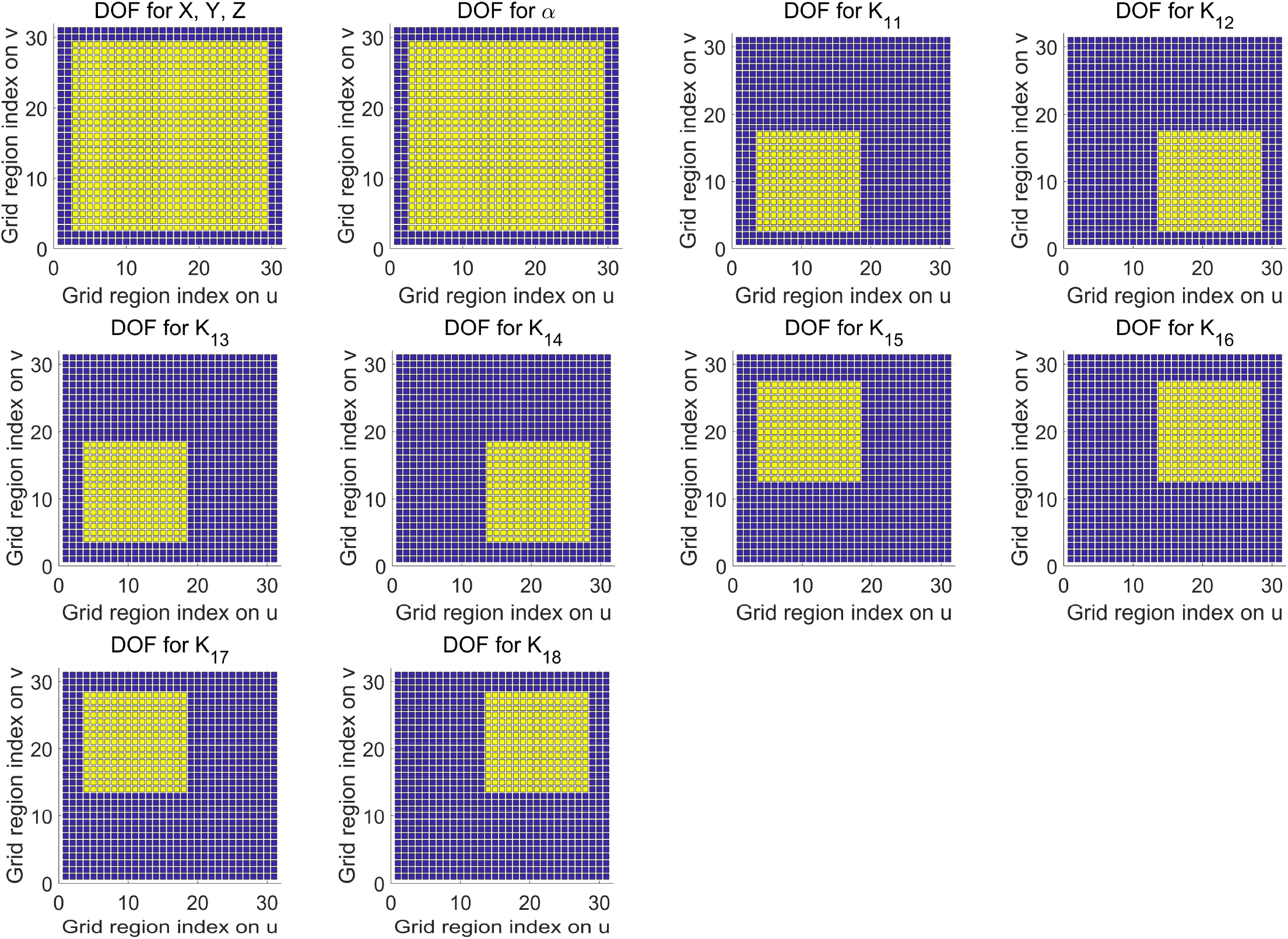
Indication of DOFs for the membrane-crosslinker complex in Figs. S9 and S10. Yellow: regions with unknowns. Navy blue: regions with fixed boundary values. The plot for *K*_2*c*_ is identical to the plot for *K*_1*c*_.

**Fig. S12.**
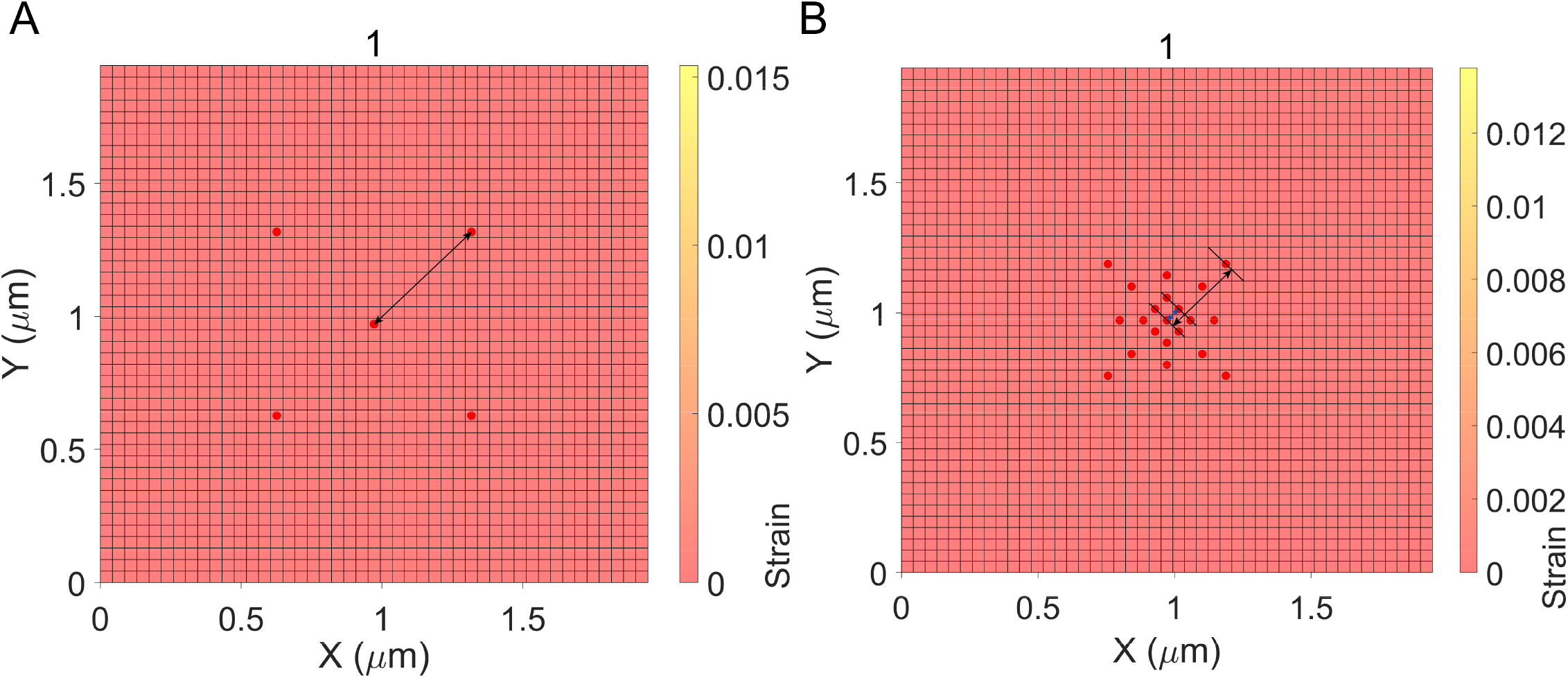
Initial membrane-crosslinker configurations for Fig. 4. (**A, B**) Top view of the initial configuration for membrane-crosslinker complexes in Fig. 4. (A) and (B) correspond to Fig. 4A and Fig. 4D, respectively. The distances indicated with arrows are 488.8 nm for (A); 61.1 nm for (B blue); and 305.5 nm for (B black).

**Fig. S13.**
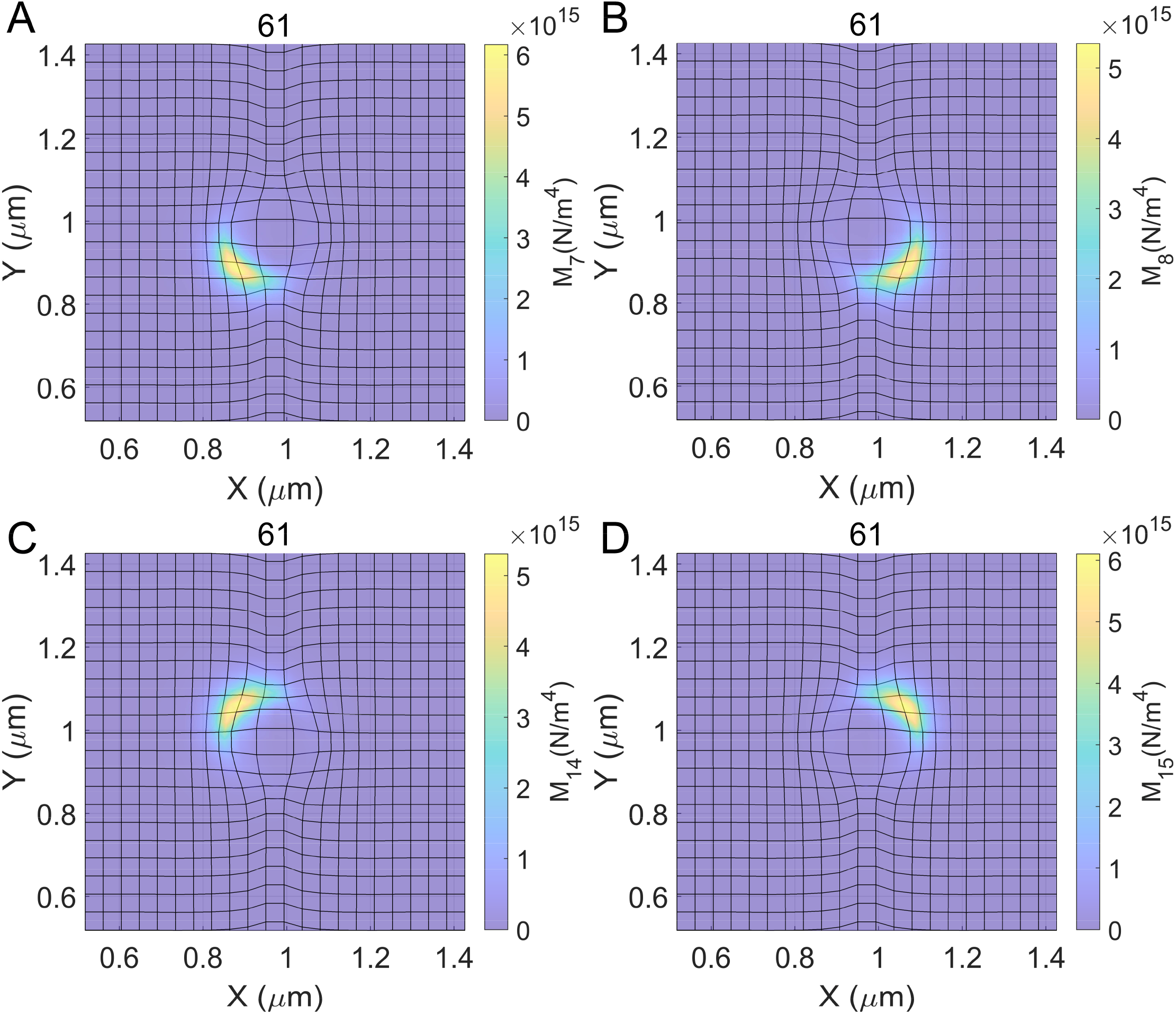
Expanded top view of the surface elastic modulus density for the membrane-crosslinker complex in Fig. 4D. The density profiles for crosslinkers with *c* = 7, 8,14,15 are plotted.

**Fig. S14.**
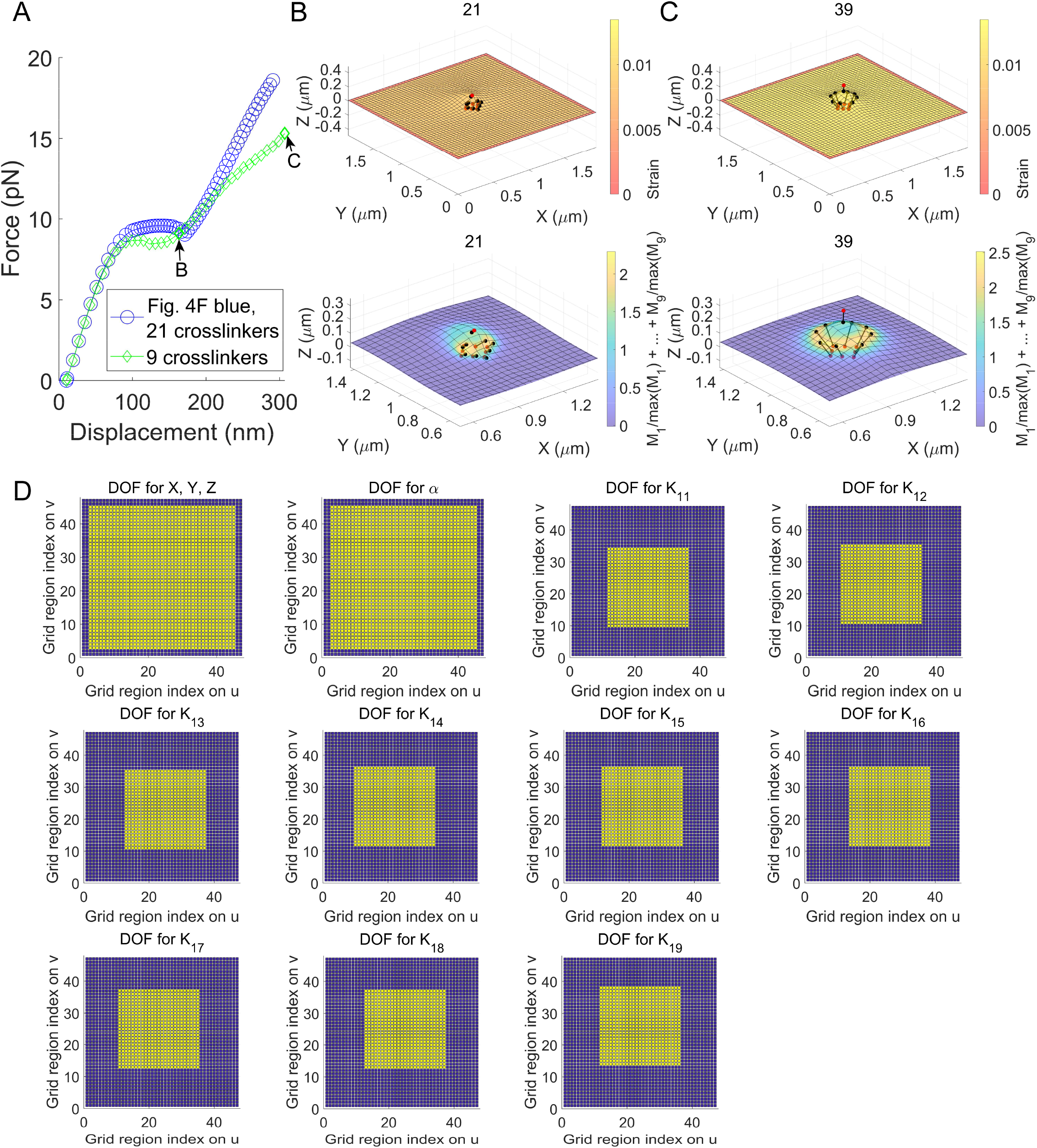
Demonstration of broken deformation symmetry. The symmetry for the membrane deformation can be broken when the total number of membrane-cytoskeleton crosslinkers clustered within a small area is not enough. (**A**) Comparison between force vs. displacement responses with 21 (Fig. 4F blue) and nine crosslinkers. The crosslinker located at the center of the membrane is displaced by varying *Z*. (**B, C**) Calculated membrane-crosslinker configurations for the green diamonds in (A). See Movie S8 for all calculated membrane-crosslinker configurations. (**D**) Indication of DOFs for data shown with green diamonds in (A). Yellow: regions with unknowns. Navy blue: regions with fixed boundary values. The plot for *K*_2*c*_ is identical to the plot for *K*_1*c*_.

**Fig. S15.**
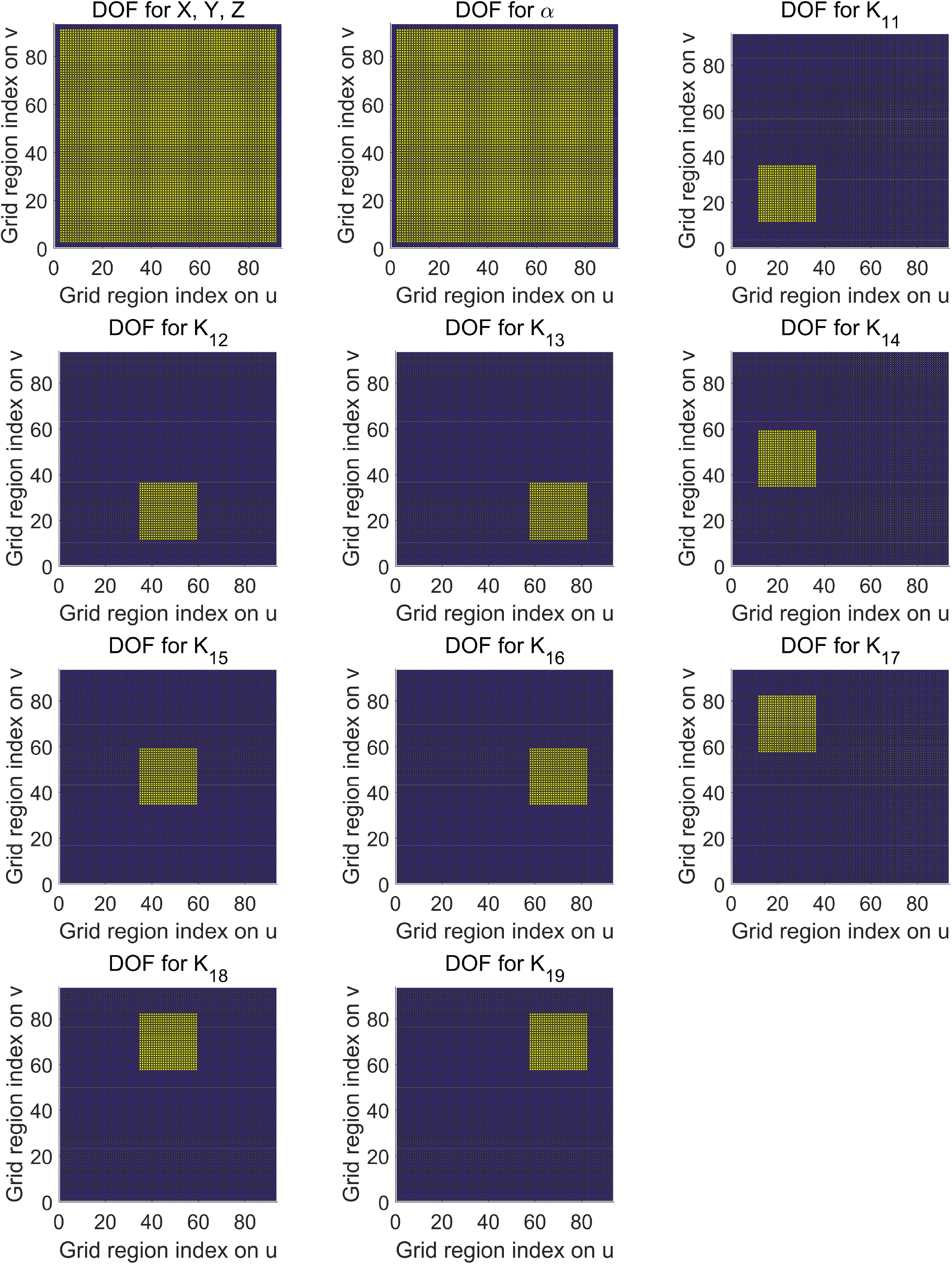
Indication of DOFs for Fig. 2. Yellow: regions with unknowns. Navy blue: regions with fixed boundary values. The plot for *K*_2*c*_ is identical to the plot for *K*_1*c*_.

**Fig. S16.**
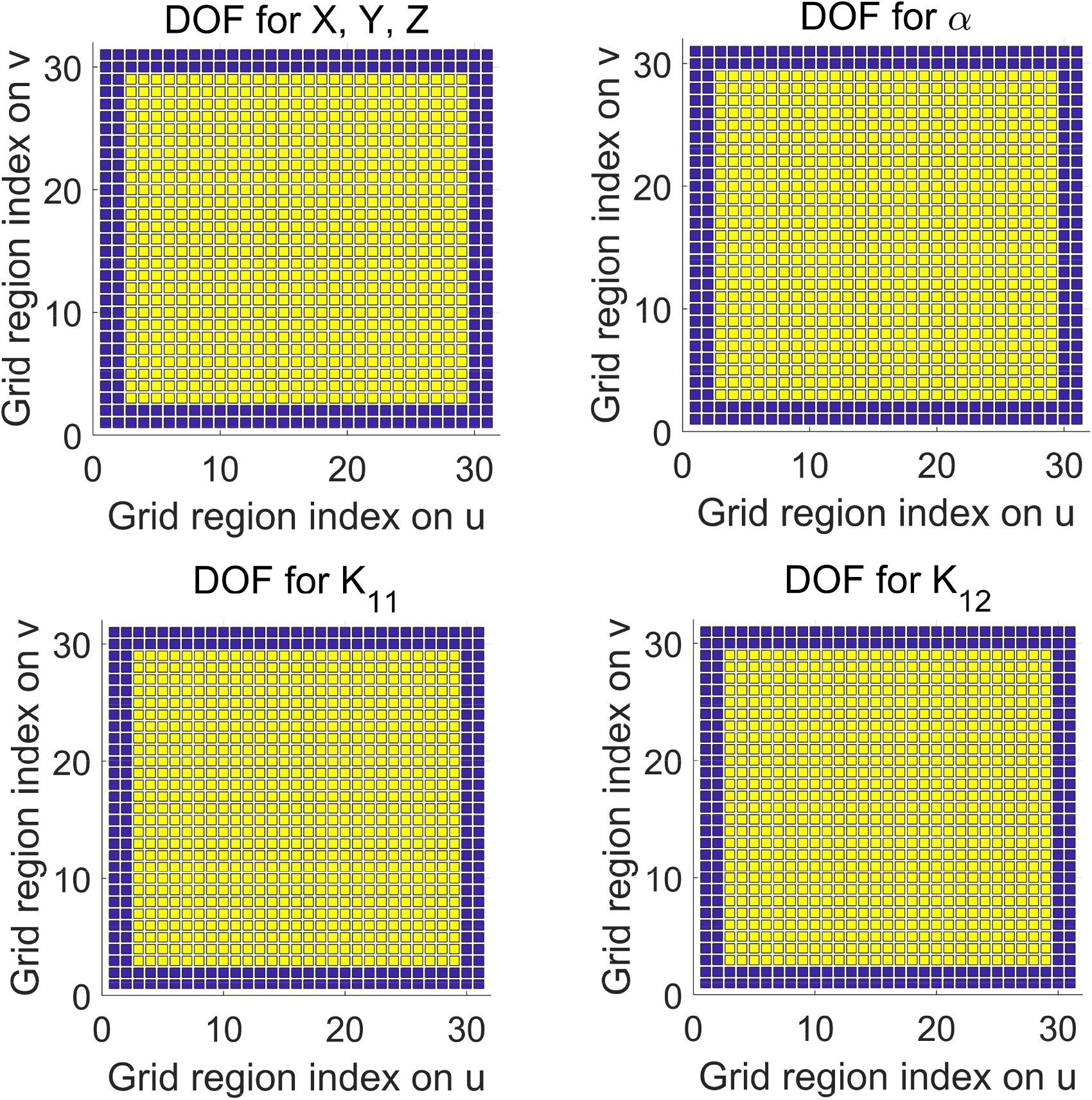
Indication of DOFs for Fig. 3. Yellow: regions with unknowns. Navy blue: regions with fixed boundary values. The plot for *K*_2*c*_ is identical to the plot for *K*_1*c*_.

**Fig. S17.**
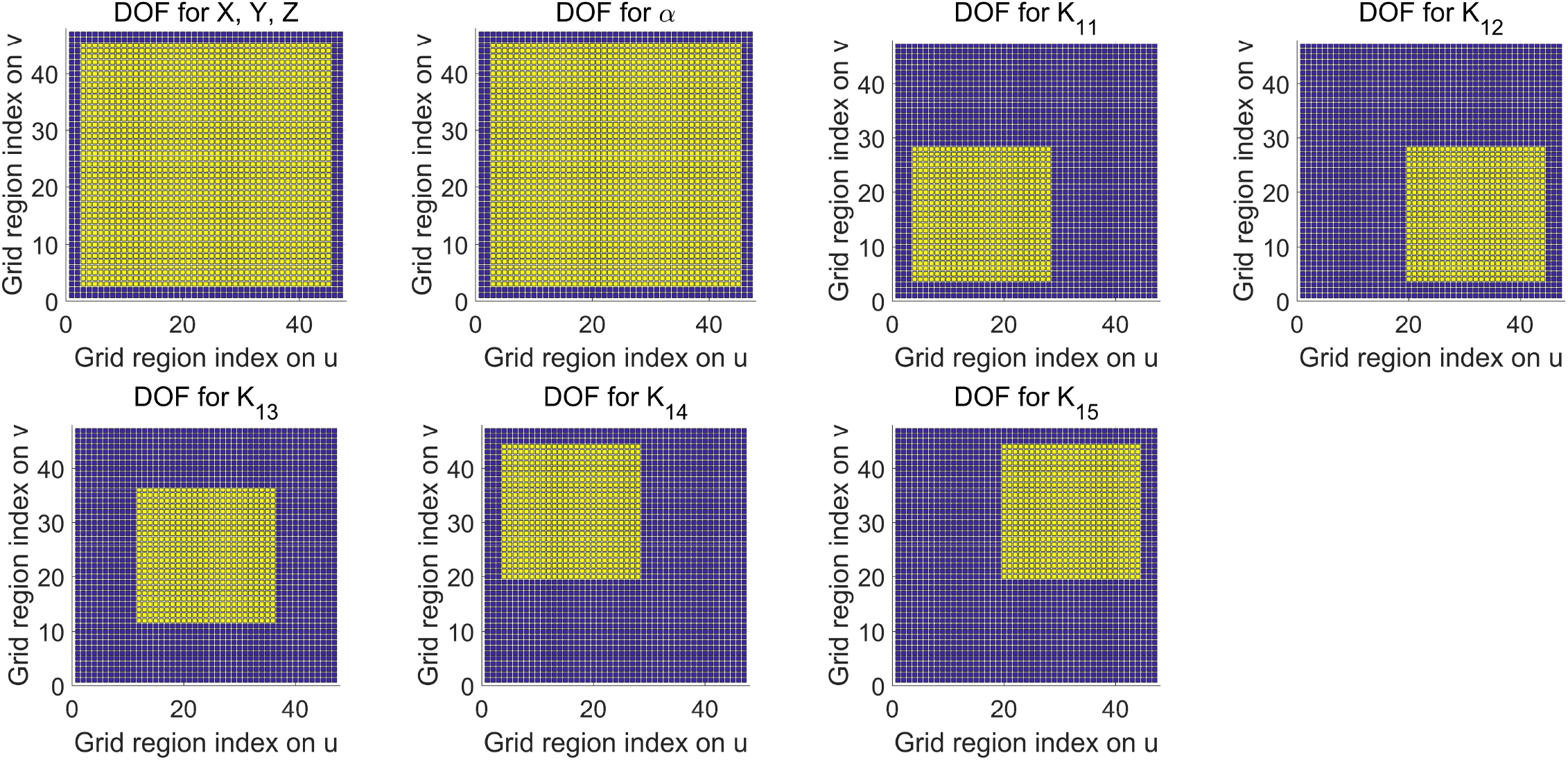
Indication of DOFs for Fig. 4A-C. Yellow: regions with unknowns. Navy blue: regions with fixed boundary values. The plot for *K*_2*c*_ is identical to the plot for *K*_1*c*_.

**Fig. S18.**
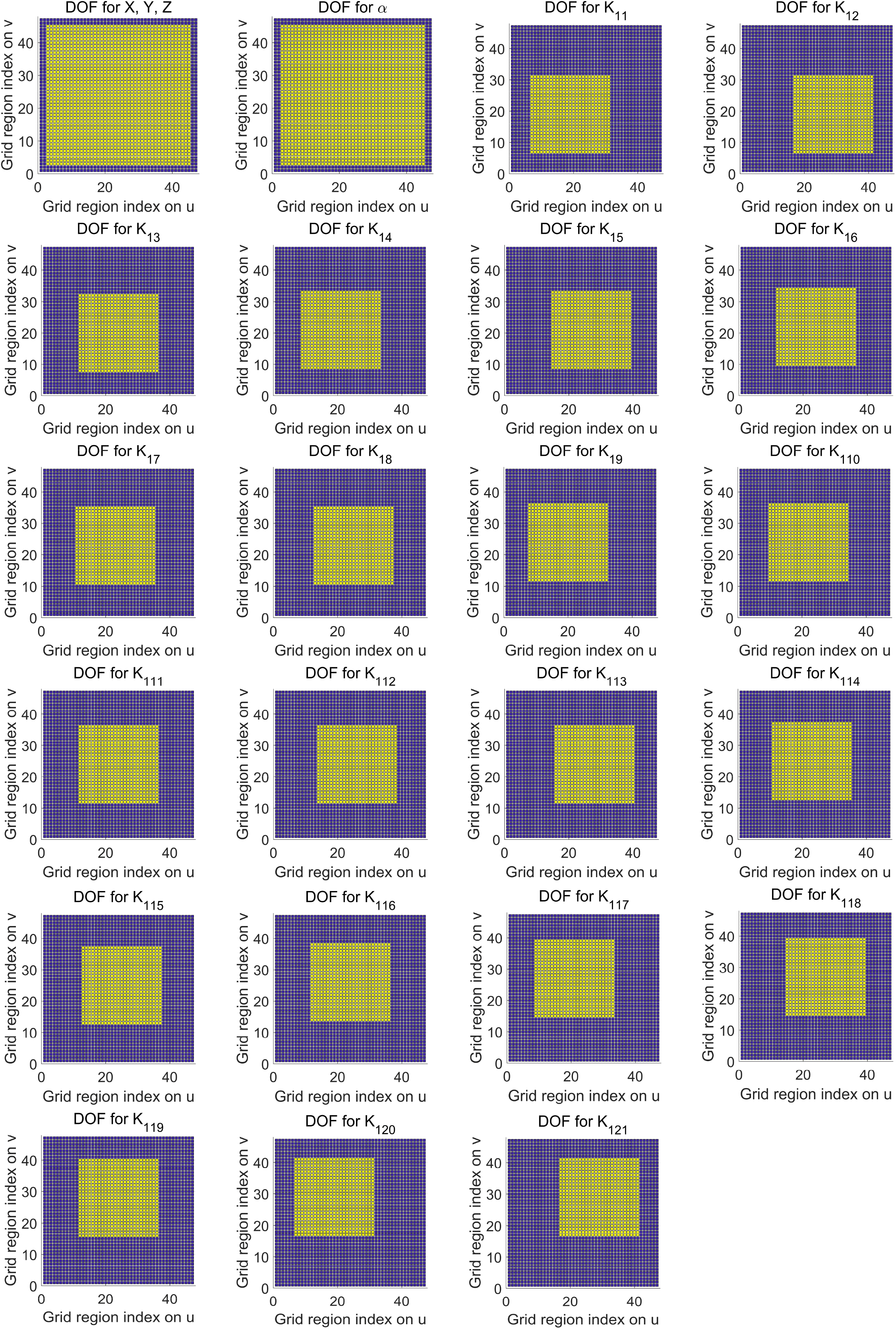
Indication of DOFs for Fig. 4D-F. Yellow: regions with unknowns. Navy blue: regions with fixed boundary values. The plot for *K*_2*c*_ is identical to the plot for *K*_1*c*_.

**Fig. S19.**
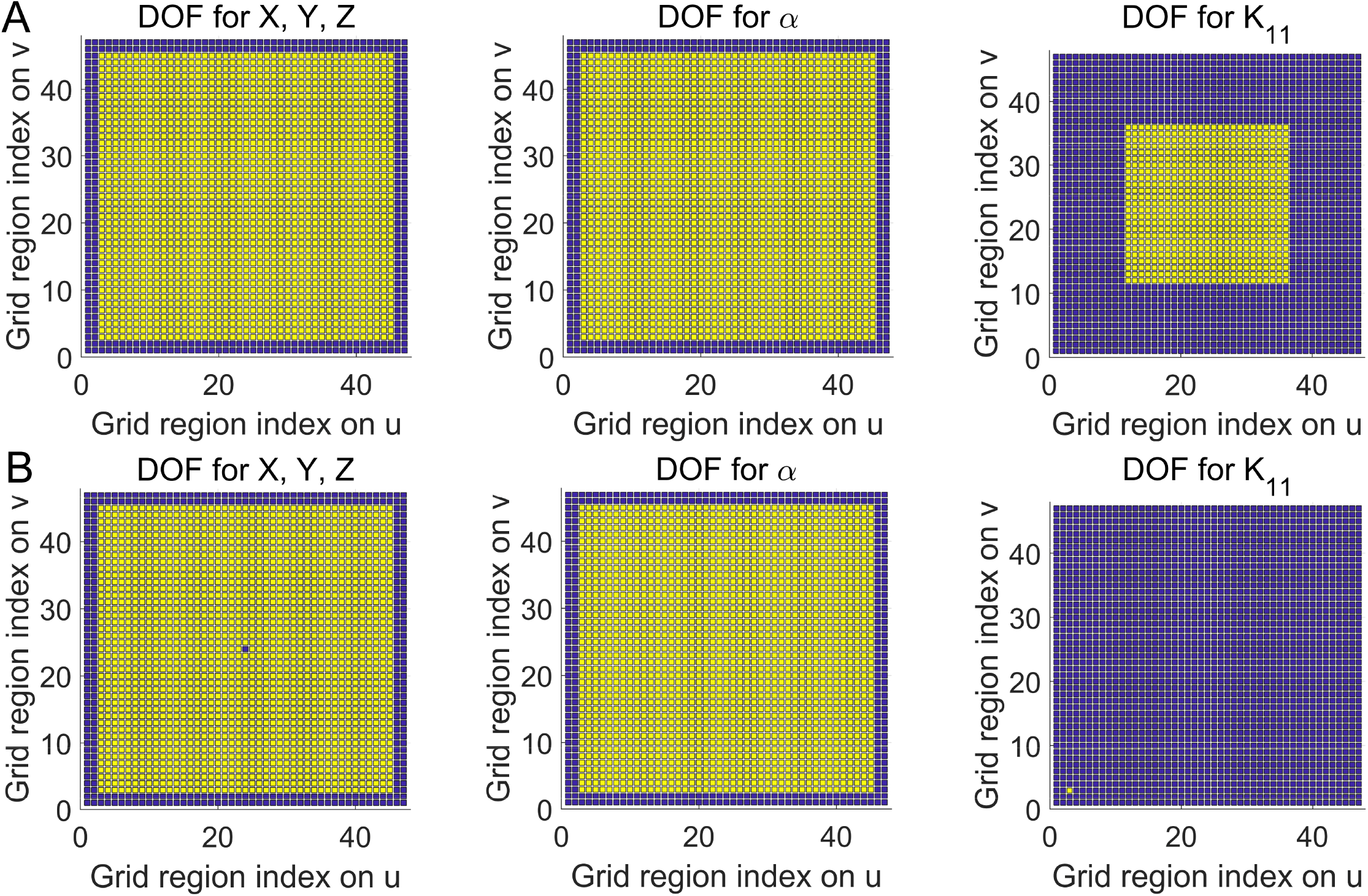
Indication of DOFs for data in Fig. 5. (**A**) DOFs for the calculations with the center crosslinker (crs force in Fig. 5). (**B**) DOFs for the calculation without the center crosslinker (membrane force in Fig. 5, black squares). A crosslinker with one element for DOFs and a negligible *φ* value (*φ* = 1 × 10^−35^) was assumed at the bottom left corner due to a programming issue in (B). Yellow: regions with unknowns. Navy blue: regions with fixed boundary values. The plot for *K*_2*c*_ is identical to the plot for *K*_1*c*_.

## Supporting Table

**Table S1.** Summary of membrane parameters used in this work. (**A**) *k*_*b*_ = 1.3806488 × 10^−23^ *J*/*K. T* = 300 *K*. (**B**) *n*_*lipid*_ was calculated from *n*_*lipid*_ = (1 − 10^−10^)_0_*A*_*initial*_ for Fig. 2, Fig. 3, and Figs. S9 and S10. *n*_*lipid*_ = *Φ*_0_*A*_*initial*_ was used for the other calculations. *A*_*initial*_ is the initial membrane area.

| Parameters (description, references, note) | Used values (Figure) |
| --- | --- |
| $k_m$ (bending modulus of lipid membranes, (47), A) | $10k_bT$, $15.7k_bT$, $21.4k_bT$, $27.1k_bT$, $32.9k_bT$, $38.6k_bT$, $44.3k_bT$, $50k_bT$ (Fig. 2); $20k_bT$ (Figs. 3 and 5; Figs. S2, S3, S9, S10, and S14; $20k_bT$ and $50k_bT$ (Fig. 4) |
| $\sigma_0$ (membrane surface tension with zero strain, (10, 37)) | $\exp(-10) \times 10^{-3}$ (all) |
| $K_{app}$ (apparent area stretching modulus, (47)) | $K_{app} = \left\{ 0.1675 \left( \frac{k_m}{k_bT} \right) - 1.972 \left( \frac{k_m}{k_bT} \right) + 166.1 \right\} 10^{-3}$ (all) |
| $\phi_0$ (reference lipid number density for zero strain, (51), B) | $\frac{1000}{629} \times 10^{18}$ (all) |

## Supporting Movies

**Movie S1**. Calculated membrane-crosslinker configurations for the analyses in Fig. 2.

**Movie S2**. Calculated membrane-crosslinker configurations for the analyses in Fig. 3.

**Movie S3**. Calculated membrane-crosslinker configurations for the analyses in Figs. S9 and S10.

**Movie S4**. Calculated membrane-crosslinker configurations for the data indicated with blue circles in Fig. 4C.

**Movie S5**. Calculated membrane-crosslinker configurations for the data indicated with red diamonds in Fig. 4C.

**Movie S6**. Calculated membrane-crosslinker configurations for the data indicated with blue circles in Fig. 4F.

**Movie S7**. Calculated membrane-crosslinker configurations for the data indicated with red diamonds in Fig. 4F.

**Movie S8**. Calculated membrane-crosslinker configurations for the data indicated with green diamonds in Fig. S14A.

## References

1. X. Pang et al., Targeting integrin pathways: mechanisms and advances in therapy. Signal Transduction and Targeted Therapy 8, 1 (2023).

2. W. H. Ziegler, A. R. Gingras, D. R. Critchley, J. Emsley, Integrin connections to the cytoskeleton through talin and vinculin. Biochemical Society Transactions 36, 235–239 (2008).

3. D. A. Calderwood et al., The talin head domain binds to integrin β subunit cytoplasmic tails and regulates integrin activation. Journal of Biological Chemistry 274, 28071–28074 (1999).

4. S. A. Freeman et al., Transmembrane pickets connect cyto-and pericellular skeletons forming barriers to receptor engagement. Cell 172, 305-317. e310 (2018).

5. T. Mori et al., Structural Basis for CD44 Recognition by ERM Proteins*. Journal of Biological Chemistry 283, 29602–29612 (2008).

6. P. B. Canham, The minimum energy of bending as a possible explanation of the biconcave shape of the human red blood cell. Journal of theoretical biology 26, 61–81 (1970).

7. W. Helfrich, Elastic properties of lipid bilayers: theory and possible experiments. Zeitschrift für Naturforschung C 28, 693–703 (1973).

8. E. Evans, W. Rawicz, Entropy-driven tension and bending elasticity in condensed-fluid membranes. Physical review letters 64, 2094 (1990).

9. W. Rawicz, K. C. Olbrich, T. McIntosh, D. Needham, E. Evans, Effect of chain length and unsaturation on elasticity of lipid bilayers. Biophysical journal 79, 328–339 (2000).

10. J. Kim, Probing nanomechanical responses of cell membranes. Scientific reports 10, 1–11 (2020).

11. F. L. Brown, Elastic modeling of biomembranes and lipid bilayers. Annu. Rev. Phys. Chem. 59, 685–712 (2008).

12. P. Alberto, C. Fiolhais, V. Gil, Relativistic particle in a box. European Journal of Physics 17, 19 (1996).

13. S. Di Martino et al., A quantum particle in a box with moving walls. Journal of Physics A: Mathematical and Theoretical 46, 365301 (2013).

14. A. Callan-Jones, B. Sorre, P. Bassereau, Curvature-driven lipid sorting in biomembranes. Cold Spring Harbor perspectives in biology 3, a004648 (2011).

15. K. AbuZineh et al., Microfluidics-based super-resolution microscopy enables nanoscopic characterization of blood stem cell rolling. Science Advances 4, eaat5304 (2018).

16. S. J. Singer, G. L. Nicolson, The Fluid Mosaic Model of the Structure of Cell Membranes: Cell membranes are viewed as two-dimensional solutions of oriented globular proteins and lipids. Science 175, 720–731 (1972).

17. T. Fujiwara, K. Ritchie, H. Murakoshi, K. Jacobson, A. Kusumi, Phospholipids undergo hop diffusion in compartmentalized cell membrane. The Journal of cell biology 157, 1071–1082 (2002).

18. M. P Clausen, B. Christoffer Lagerholm, The probe rules in single particle tracking. Current Protein and Peptide Science 12, 699–713 (2011).

19. Y.-J. Chai, C.-Y. Cheng, Y.-H. Liao, C.-H. Lin, C.-L. Hsieh, Heterogeneous nanoscopic lipid diffusion in the live cell membrane and its dependency on cholesterol. Biophysical Journal 121, 3146–3161 (2022).

20. S. Spindler et al., Visualization of lipids and proteins at high spatial and temporal resolution via interferometric scattering (iSCAT) microscopy. Journal of Physics D: Applied Physics 49, 274002 (2016).

21. E. C. Arnspang, J. Schwartzentruber, M. P. Clausen, P. W. Wiseman, B. C. Lagerholm, Bridging the gap between single molecule and ensemble methods for measuring lateral dynamics in the plasma membrane. PLoS One 8, e78096 (2013).

22. K. G. Suzuki, A. Kusumi, Refinement of Singer-Nicolson fluid-mosaic model by microscopy imaging: Lipid rafts and actin-induced membrane compartmentalization. Biochimica et Biophysica Acta (BBA)-Biomembranes 1865, 184093 (2023).

23. M. Yao et al., The mechanical response of talin. Nature communications 7, 1–11 (2016).

24. S. J. Tan et al., Regulation and dynamics of force transmission at individual cell-matrix adhesion bonds. Science Advances 6, eaax0317 (2020).

25. P. Ringer et al., Multiplexing molecular tension sensors reveals piconewton force gradient across talin-1. Nature methods 14, 1090–1096 (2017).

26. X. Hu et al., Cooperative vinculin binding to talin mapped by time-resolved super resolution microscopy. Nano letters 16, 4062–4068 (2016).

27. A. Del Rio et al., Stretching single talin rod molecules activates vinculin binding. Science 323, 638–641 (2009).

28. C. Grashoff et al., Measuring mechanical tension across vinculin reveals regulation of focal adhesion dynamics. Nature 466, 263–266 (2010).

29. P. Atherton et al., Vinculin controls talin engagement with the actomyosin machinery. Nature communications 6, 10038 (2015).

30. A. Kumar et al., Talin tension sensor reveals novel features of focal adhesion force transmission and mechanosensitivity. Journal of Cell Biology 213, 371–383 (2016).

31. F. Margadant et al., Mechanotransduction in vivo by repeated talin stretch-relaxation events depends upon vinculin. PLoS biology 9, e1001223 (2011).

32. O. Rossier et al., Integrins β1 and β3 exhibit distinct dynamic nanoscale organizations inside focal adhesions. Nature cell biology 14, 1057–1067 (2012).

33. X. H. Yang et al., CD151 restricts the α6 integrin diffusion mode. Journal of cell science 125, 1478–1487 (2012).

34. J. W. Yuan et al., Diffusion behaviors of Integrins in single cells altered by epithelial to mesenchymal transition. Small 18, 2106498 (2022).

35. J. Kim, Unconventional mechanics of lipid membranes: a potential role for mechanotransduction of hair cell stereocilia. Biophysical journal 108, 610–621 (2015).

36. H. Alimohamadi, R. Vasan, J. Hassinger, J. Stachowiak, P. Rangamani, The role of traction in membrane curvature generation. Molecular Biology of the Cell 29, 2024–2035 (2018).

37. J. Kim, A possible molecular mechanism for mechanotransduction at cellular focal adhesion complexes. Biophysical Reports 1, 100006 (2021).

38. E. Nelson, Derivation of the Schrödinger equation from Newtonian mechanics. Physical review 150, 1079 (1966).

39. W. T. Strunz, L. Diósi, N. Gisin, T. Yu, Quantum trajectories for Brownian motion. Physical Review Letters 83, 4909 (1999).

40. K. Mita, Schrödinger’s equation as a diffusion equation. American Journal of Physics 89, 500–510 (2021).

41. A. Kusumi et al., Paradigm shift of the plasma membrane concept from the two-dimensional continuum fluid to the partitioned fluid: high-speed single-molecule tracking of membrane molecules. Annu. Rev. Biophys. Biomol. Struct. 34, 351–378 (2005).

42. K. M. Spillane et al., High-speed single-particle tracking of GM1 in model membranes reveals anomalous diffusion due to interleaflet coupling and molecular pinning. Nano letters 14, 5390–5397 (2014).

43. R. Schmidt et al., MINFLUX nanometer-scale 3D imaging and microsecond-range tracking on a common fluorescence microscope. Nature Communications 12, 1478 (2021).

44. D. Raucher, M. P. Sheetz, Characteristics of a membrane reservoir buffering membrane tension. Biophysical journal 77, 1992–2002 (1999).

45. Z. Shi, Z. T. Graber, T. Baumgart, H. A. Stone, A. E. Cohen, Cell membranes resist flow. Cell 175, 1769-1779. e1713 (2018).

46. H. De Belly et al., Cell protrusions and contractions generate long-range membrane tension propagation. Cell 186, 3049-3061. e3015 (2023).

47. J. Kim, A Review of Continuum Mechanics for Mechanical Deformation of Lipid Membranes. Membranes 13, 493 (2023).

48. R. Rangarajan, H. Gao, A finite element method to compute three-dimensional equilibrium configurations of fluid membranes: Optimal parameterization, variational formulation and applications. Journal of Computational Physics 297, 266–294 (2015).

## Additional references

49. V. Petrenko et al., Type 2 diabetes disrupts circadian orchestration of lipid metabolism and membrane fluidity in human pancreatic islets. PLoS Biology 20, e3001725 (2022).

50. S. G. Snowden et al., Development and application of high-throughput single cell lipid profiling: a study of SNCA-A53T human dopamine neurons. Iscience 23 (2020).

51. H. I. Petrache, S. W. Dodd, M. F. Brown, Area per lipid and acyl length distributions in fluid phosphatidylcholines determined by 2H NMR spectroscopy. Biophysical journal 79, 3172–3192 (2000).

